# Substrate-Dependent Crosslinking by the Cytochrome P450 from Aminopyruvatide Biosynthesis

**DOI:** 10.64898/2026.05.07.723658

**Authors:** Chandrashekhar Padhi, Dinh T. Nguyen, Lingyang Zhu, Lide Cha, Jesse W. Wald, Douglas A. Mitchell, Wilfred A. van der Donk

**Affiliations:** Department of Chemistry and Howard Hughes Medical Institute, 600 South Mathews Avenue, University of Illinois at Urbana−Champaign, Urbana, Illinois 61801, USA; Carl R. Woese Institute for Genomic Biology, University of Illinois at Urbana-Champaign, 1206 West Gregory Drive, Urbana, Illinois, 61801, USA; School of Chemical Sciences NMR Laboratory, University of Illinois at Urbana-Champaign, Urbana, 61801, IL, USA; Departments of Biochemistry and Chemistry, Vanderbilt University School of Medicine, Nashville, TN, United States

**Keywords:** Biosynthesis, cytochrome P450, α-keto acid, macrocyclic scaffolds, protease inhibition

## Abstract

Cytochrome P450s catalyze an array of reactions including crosslinking of aromatic side chains in the biosynthesis of ribosomally synthesized and post-translationally modified peptides (RiPPs). ApyO is a cytochrome P450 that forms a C−C bond between two tyrosines in a YLY motif in the substrate ApyA, the precursor peptide of the RiPP aminopyruvatide. We utilized cell-free translation to generate ApyA variants and probe the substrate tolerance of ApyO. Through Alphafold-based modelling and *in vitro* assays, we show that ApyO accepts the 10 C-terminal residues of ApyA and requires a conserved Arg/Lys in the substrate. Inspired by substrate sequences in orthologous biosynthetic gene clusters, we substituted one of the tyrosine residues with a tryptophan and observed that ApyO catalyzed formation of an N−C bond between the indole of Trp and Cε2 of Tyr. ApyO unexpectedly catalyzed formation of a C−O bond between the two tyrosine residues when we substituted the leucine residue in the YLY motif with tyrosine or tryptophan. A peptide containing a biaryl linkage and C-terminal aminopyruvate displayed sub-nanomolar inhibition of select proteases with the aminopyruvate group critical for activity. Overall, this study demonstrates plasticity in the manner of macrocyclization catalyzed by the P450 ApyO.

## Introduction

Genes encoding cytochrome P450 enzymes are ubiquitous in all domains of life and play important roles in both primary and secondary metabolism [1]. The versatility of their catalytic potential has been explored extensively in biomedicine and biotechnology [2–5]. Strategies for bioengineering P450 enzymes include substrate diversification, enzyme engineering, and exploring non-canonical redox partners [6–9]. P450 enzymes incorporate a variety of post-translational modifications (PTMs) in ribosomally synthesized and post-translationally modified peptides (RiPPs). The array of RiPP modifications catalyzed by P450s include oxidative decarboxylation, hydroxylation, epoxidation, and macrocyclization [10–14].

Biarylitides and cittilins are RiPPs that employ a P450 enzyme to form macrocyclic products in scaffolds containing three or four amino acid residues resulting from the oxidative formation of C−C, C−N and C−O bonds between two aromatic rings (e.g. Figure 1) or between aromatic and aliphatic side chains [15–18]. Bioinformatic and machine learning-based genome mining approaches have highlighted the widespread occurrence of these crosslinked tri- and tetra-peptide scaffolds throughout the bacterial domain [19–23]. A related dipeptide scaffold is installed by a P450 during the biosynthesis of the lasso peptides nocapeptin and longipeptin [24]. Researchers have engineered a subset of these enzymes to alter the macrocyclic linkages expanding the chemical space of their products [16, 25].

**Figure 1.**
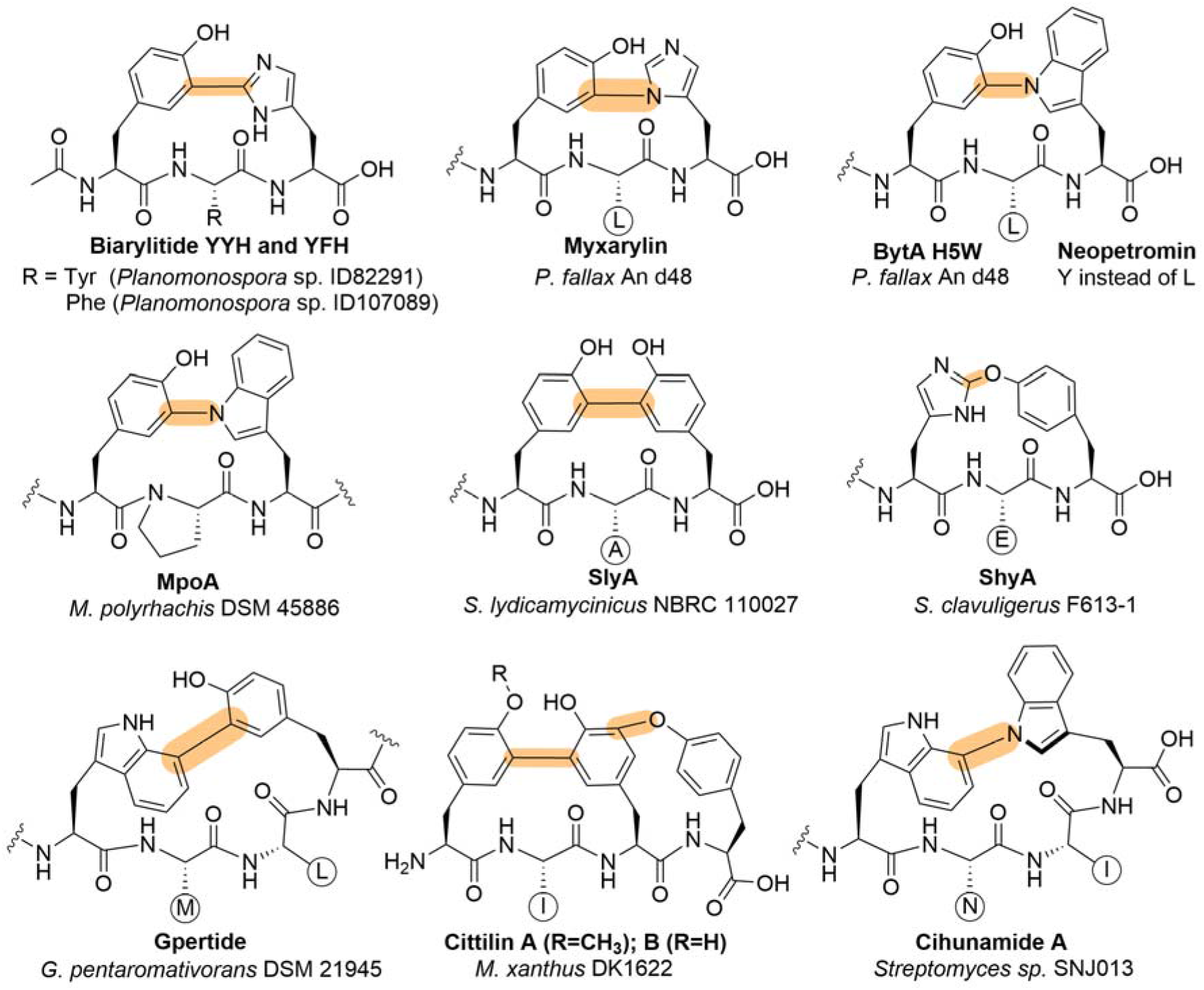
Examples of RiPP products with P450-mediated crosslinks. Examples of tri-/tetra-peptide scaffolds crosslinked by cytochrome P450 enzymes that feature C−C, C−O and C−N linkages between two aromatic residues. For brevity, the residues in between the crosslinked residues are annotated in a circle by their one letter amino acid code.

Previously, we reported a P450 enzyme involved in the biosynthesis of aminopyruvatides [26]. The aminopyruvatide biosynthetic gene cluster (BGC) in *Burkholderia thailandensis* E264 (*apy* cluster, Figure 2) occurs widely in *Burkholderia* genomes. It encodes a precursor peptide (ApyA) with a C-terminal YLYD motif. PTM enzymes encoded in the BGC include a multinuclear non-heme iron-dependent oxidation enzyme (MNIO; ApyH) that transforms the C-terminal Asp residue into an α-keto β-amino acid, and a P450 enzyme (ApyO) that crosslinks the two tyrosines by a Cε−Cε linkage.

**Figure 2.**
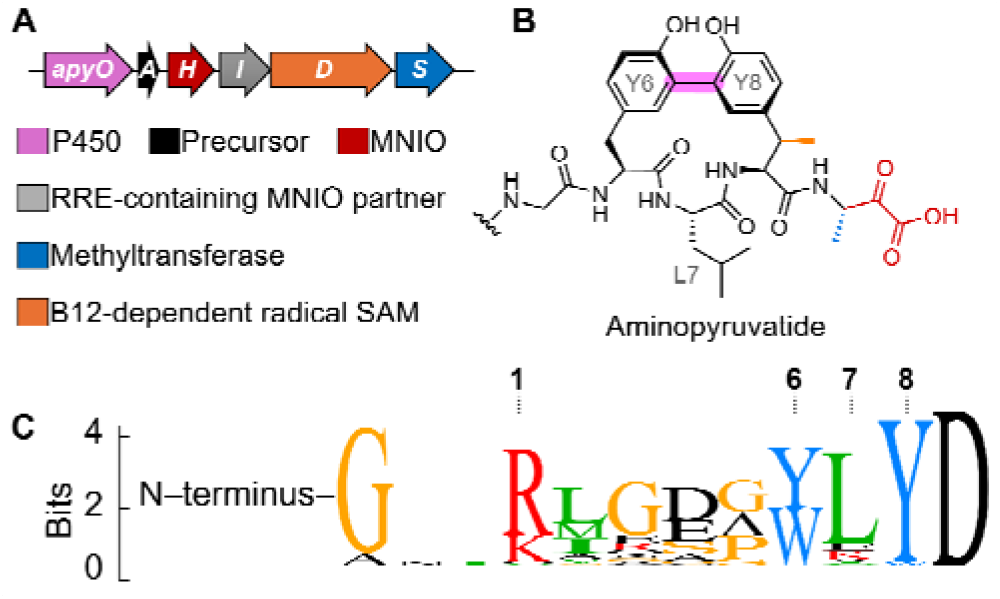
Aminopyruvatide BGC, the corresponding product and partial sequence logo of putative precursors from orthologous BGCs. (A) The *apy* BGC including the gene encoding the P450 enzyme ApyO (pink) that catalyzes the formation of a biaryl crosslink between the two C-terminal tyrosines (pink highlight in panel B) in the YLY motif of the precursor ApyA. Color coding is coordinated for panels A and B. Residue numbering is based on an AlphaFold model for ApyO in complex with ApyA derived peptides (vide infra). RRE, RiPP recognition element [29]; SAM, *S*-adenosyl methionine. (C) A partial sequence logo of the C-terminal region of homologous precursor peptides from orthologous BGCs highlighting the aromatic residues preceding the C-terminal Asp. Residues evaluated in this study are numbered in panels B and C. Colors for panel C are independent. For full length precursor peptide sequences, see Figure S1.

ApyO catalyzes the macrocyclization of unmodified ApyA [26], allowing investigation of its substrate tolerance regardless of whether other enzymes encoded in the BGC accept such changes. Putative precursor sequences in orthologous BGCs suggested conservation of one arginine (**Arg1**) and a leucine (**Leu7**) in the C-terminal **R**MGEGY**L**YD motif of ApyA (Figure S1). These sequences also suggested the possibility of alternative biaryl linkages beyond the C−C bond between two tyrosines (Tyr6 and Tyr8) because both WLY and YLW motifs occur naturally. Therefore, in this study we investigated the substrate scope of ApyO with respect to Arg1, Tyr6, Leu7, and Tyr8. First, we established an *in vitro* assay that allowed identification of a short peptide substrate and determined the kinetic parameters of crosslink formation. Then we utilized cell-free expression (CFE) to generate variants of the truncated substrate. Subsequent modification of the substrate variants by ApyO led to the discovery of an N−C linkage between Trp and Tyr residues and constitutional isomeric products of the Tyr−Tyr biaryl linkage that were macrocyclized either through the native C−C linkage or a non-native C−O linkage. While previous studies have typically substituted the residues participating in crosslinking to biosynthesize new molecules, in this study we show that altering the residues adjacent to the modified residues also can access alternate chemistry. This study also highlights that mass spectrometric analysis alone is not sufficient to structurally characterize the crosslinked products, as noted previously [18].

## Results and Discussion

### *In silico* analysis of aminopyruvatide-like BGCs

A previous bioinformatic analysis identified multiple aminopyruvatide-like BGCs encoding homologues of the precursor peptide (ApyA; NCBI accession: ABC34935.1) and the cytochrome P450 enzyme (ApyO; ABC35200.1) in addition to other modifying enzymes [26]. To identify additional precursor peptides of this type, the ApyA peptide sequence was subjected to Pattern Hit Initiated BLAST (PHI-BLAST) [27] analysis using the ApyA sequence as query and the YLY motif as the PHI pattern for 10 iterations at an inclusion threshold of 0.05. In total, 53 nonredundant sequences were identified containing a tripeptide motif with aromatic residues, *i*.*e*., YxY (n = 25), WxY (n = 26), and YLW (n = 2) (Figure S1). Among the 25 sequences containing a YxY motif, the x residue was mostly Leu (n = 17) followed by Lys (n = 4), Glu (n = 2) and Ile or Ala (n = 1, each). Similarly, for the 26 peptides containing a WxY motif, Leu was predominantly encoded at the x position (n = 23) followed by Glu (n = 2) and Val (n = 1). The retrieved NCBI accession identifiers were subsequently subjected to RODEO analysis [28] to obtain the genome neighborhood of the identified putative precursor peptides. For all queries, a gene encoding a P450 enzyme was present in the BGC.

### Ortholog-guided mutation of *apyA*

To investigate the macrocyclization pattern of the naturally occurring ApyA orthologs containing WxY and YxW motifs, we mutated ApyA to introduce such motifs (ApyA-Y6W and -Y8W; for residue numbering, see Figure 2) and co-expressed the peptides with ApyO in *Escherichia coli*. ApyO successfully modified both the Y6W and Y8W variants as evident from the 2 Da mass loss observed by liquid chromatography coupled to mass spectrometric detection (LC-MS), characteristic of a crosslinking event (Figure 3A and B). Tandem MS analysis suggested a crosslink between the aromatic residues since b and y ions were not observed within the WLY and YLW motifs (Figures S2 and S3, respectively). Because of the relatively high occurrence of the WLY motif (n = 23) compared to the YLW motif (n = 2) among the putative precursors in orthologous BGCs (Figure 2C), we selected the ApyA-Y6W variant for further studies.

**Figure 3.**
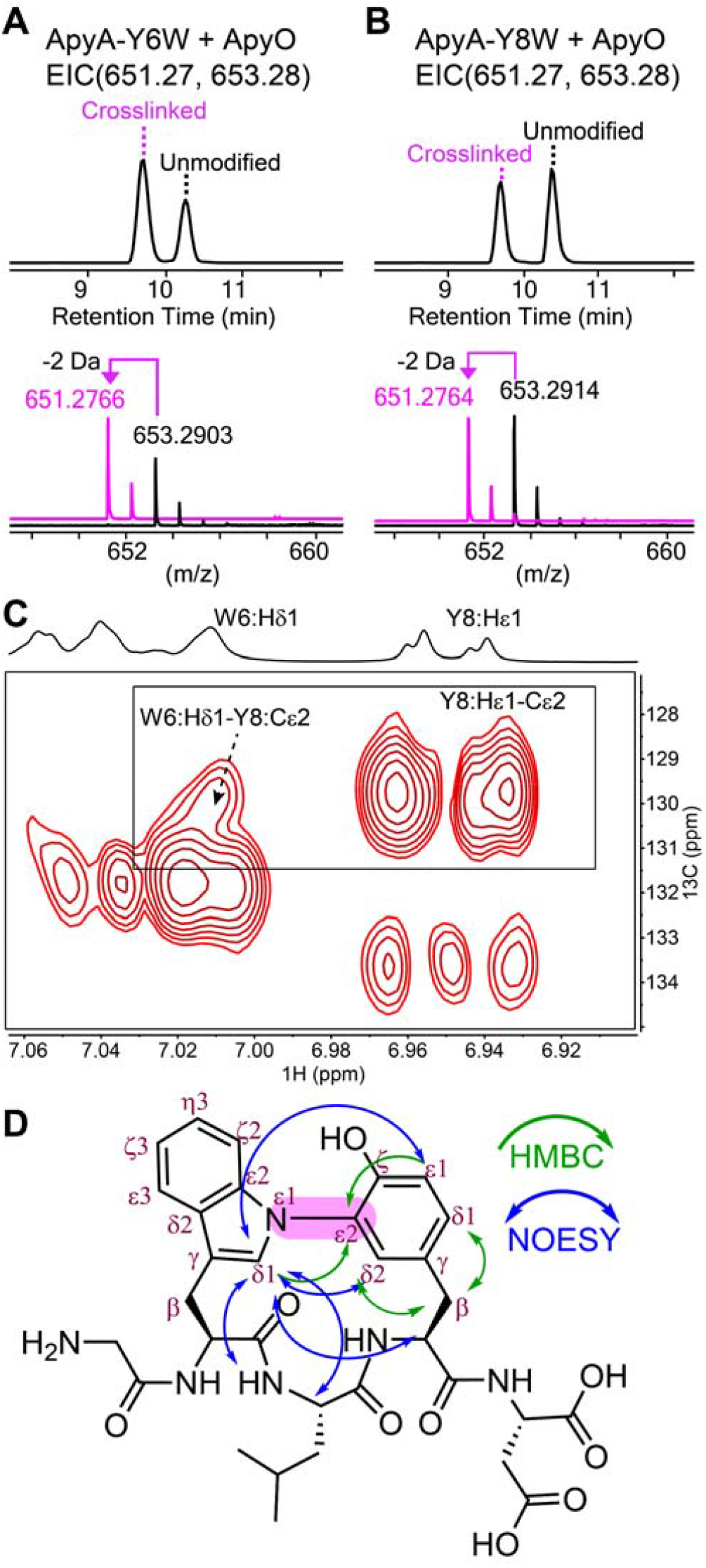
HRMS analysis and structural characterization of ApyA variants co-expressed with ApyO. (A) LC-MS analysis of ApyA-Y6W co-expressed with ApyO and digested with endoproteinase GluC. The extracted ion chromatogram (EIC) is shown with the calculated monoisotopic masses (top) and HRMS analysis (bottom) of the observed monoisotopic masses as [M+H]^+^ corresponding to the unmodified and macrocyclized compounds. (B) Same as panel A but for the ApyA-Y8W variant. (C) HMBC correlation between the Hδ1 of Trp6 and Cε2 of Tyr8 in d_6_-DMSO supplemented with 0.1% d-formic acid at 55 **°**C. HR-MS/MS analysis for panels A and B is shown in Figures S2-S3. (D) structure of the GluC-digested ApyA-Y6W peptide showing the macrocyclization between the Nε1 of Trp6 and Cε2 of Tyr8 as evidenced by the HMBC (green) and NOESY (blue) correlations (protons involved are not drawn).

### Structural characterization of ApyO-modified ApyA-Y6W

To substantiate the MS results, we purified the ApyO-modified N-terminally His_6_-tagged ApyA-Y6W from *E. coli* using immobilized metal-affinity chromatography. The purified peptide was digested with GluC endopeptidase to release a C-terminal pentapeptide (GWLYD), which contained the −2 Da modification. The pentapeptide fragment was purified by HPLC (Figure S4) and characterized by NMR spectroscopy using one-dimensional (1D) and two-dimensional (2D) NMR experiments including ^1^H-^1^H TOCSY, ^1^H-^1^H NOESY, ^1^H-^13^C-HSQC and ^1^H-^13^C HMBC. To improve resolution, experiments were carried out at 55 °C. The ^1^H and ^13^C assignments of the peptide in DMSO-d_6_ at 55 °C are given in Table S2. All expected aromatic protons of Trp6 were observed and were consistent with the expected pattern for a tryptophan residue (Figure S5A). In contrast, the aromatic protons of Tyr8 displayed splitting patterns that deviated from a typical tyrosine. Specifically, the 1D ^1^H NMR signals extracted from ^1^H-^1^H TOCSY data only showed three aromatic protons exhibiting two doublets and one broad singlet (Figure S5A). These data suggested a crosslink between the Trp6 indole nitrogen and either Cδ or Cε of Tyr8. More detailed analysis of the ^1^H-^13^C HSQC (Figure S5B) and HMBC (Figure S6A) spectra supported an N−C cross-link between the indole nitrogen of Trp6 and Cε of Tyr8. This crosslink was confirmed by a cross peak between the Hδ1 proton (7.01 ppm) of Trp6 and the Cε2 carbon (129.8 ppm) of Tyr8, and between the Hβ protons (2.97, 2.80 ppm) and Cδ1/Cδ2 of Tyr8 (131.9/131.3 ppm) (Figure 3C and D). NOESY cross peaks between protons on Trp6 and Tyr8, including interactions between their aromatic protons further support this crosslink (Figure 3D and S6B). The assignments are also consistent with the observed 2 Da mass loss detected by HR-MS (Figure 3A).

### AlphaFold-based prediction of substrate binding

ApyO accepts unmodified ApyA (66 amino acids) for crosslink formation without the need of the other PTMs introduced by the *apy* biosynthetic enzymes [26]. This feature is advantageous for exploring the substrate tolerance of the enzyme, which would be further facilitated if the enzyme were to show crosslinking activity with shorter peptides. An AlphaFold3 model of ApyO in complex with the C-terminal 10-mer peptide of ApyA (GRMGEGYLYD) placed both Tyr residues of the YLY motif near the heme and suggested an interaction between the Arg in the peptide (Arg1, Figure 1C) and a hydrophilic pocket formed by Glu20, Asp422 and Asp459 of ApyO (Figures 4A, B and S7). The sequence alignment of aminopyruvatide precursor peptides shows a prevalence of Arg in position 1 (74%), with a conservative Lys substitution in most other peptides (22%) (Figure S1). To test whether this 10-mer C-terminal (ct) peptide was a viable substrate, we prepared a version where the N-terminal Gly was substituted with formyl-Met using CFE (fMRMGEGYLYD; ApyA_ct_). Subsequently, purified ApyO and redox partners (ferredoxin, ferredoxin reductase and NADPH) were added to the CFE reaction. Analysis by Matrix-Assisted Laser Desorption/Ionization Time-of-Flight (MALDI-ToF) mass spectrometry and by LC-MS showed that ApyO successfully modified ApyA_ct_ (~70%, Figure 5) validating the 10-mer as a viable substrate. Shortening of the substrate further to an 8-mer (fMGEGYLYD) did not result in product formation (Figure S8A). We therefore conducted all experiments to test the AlphaFold model with the 10-mer peptide. We used a synthetic peptide (N-terminally acetylated-GRMGEGYLYD; Ac-ApyA_ct_) to determine a *k*_cat_ of 1.63 ± 0.04 min^−1^ and a *K*_m_ of 80 ± 7 µM (Figure S8B). Kinetic studies on RiPP-related P450 enzymes are scarce but the kinetic parameters of ApyO lie within the range of other P450s involved in natural product tailoring reactions [30–33].

**Figure 4.**
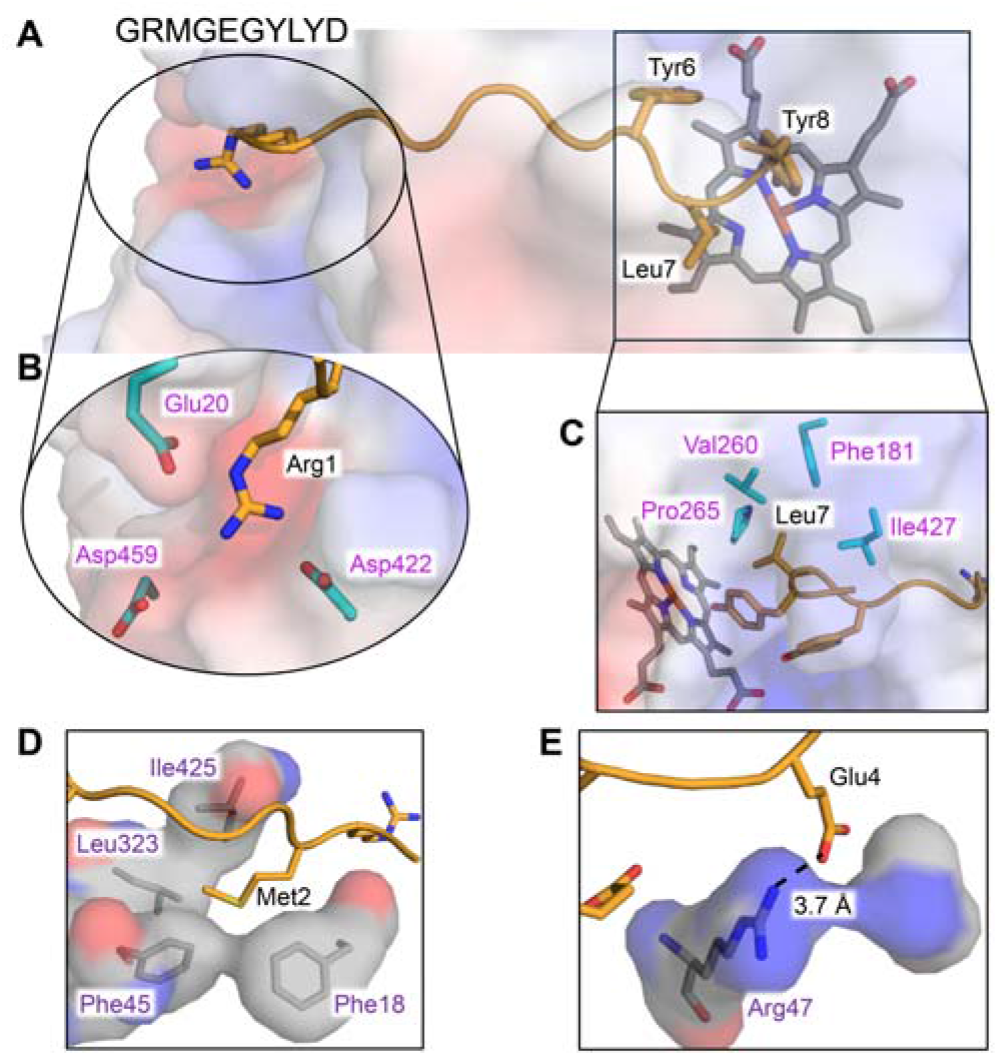
AlphaFold3 model of ApyO in complex with the C-terminal 10-mer of ApyA. (A) Model of the ApyA_ct_ precursor peptide bound to ApyO showing Tyr6 and Tyr8 near the active site heme. (B) Zoom-in image of the Arg1 residue embedded in a hydrophilic pocket of ApyO. (C) Zoom-in image of the ApyA_ct_ Leu7 residue occupying a hydrophobic pocket of ApyO. (D) Zoom-in image of Apy_ct_ Glu4 and Arg47 from ApyO illustrating a potential interaction. (E) Zoom-in image of the ApyA_ct_ Met2 occupying a hydrophobic pocket of ApyO. For the full structure of the 10-mer bound to ApyO, see Figure S7.

**Figure 5.**
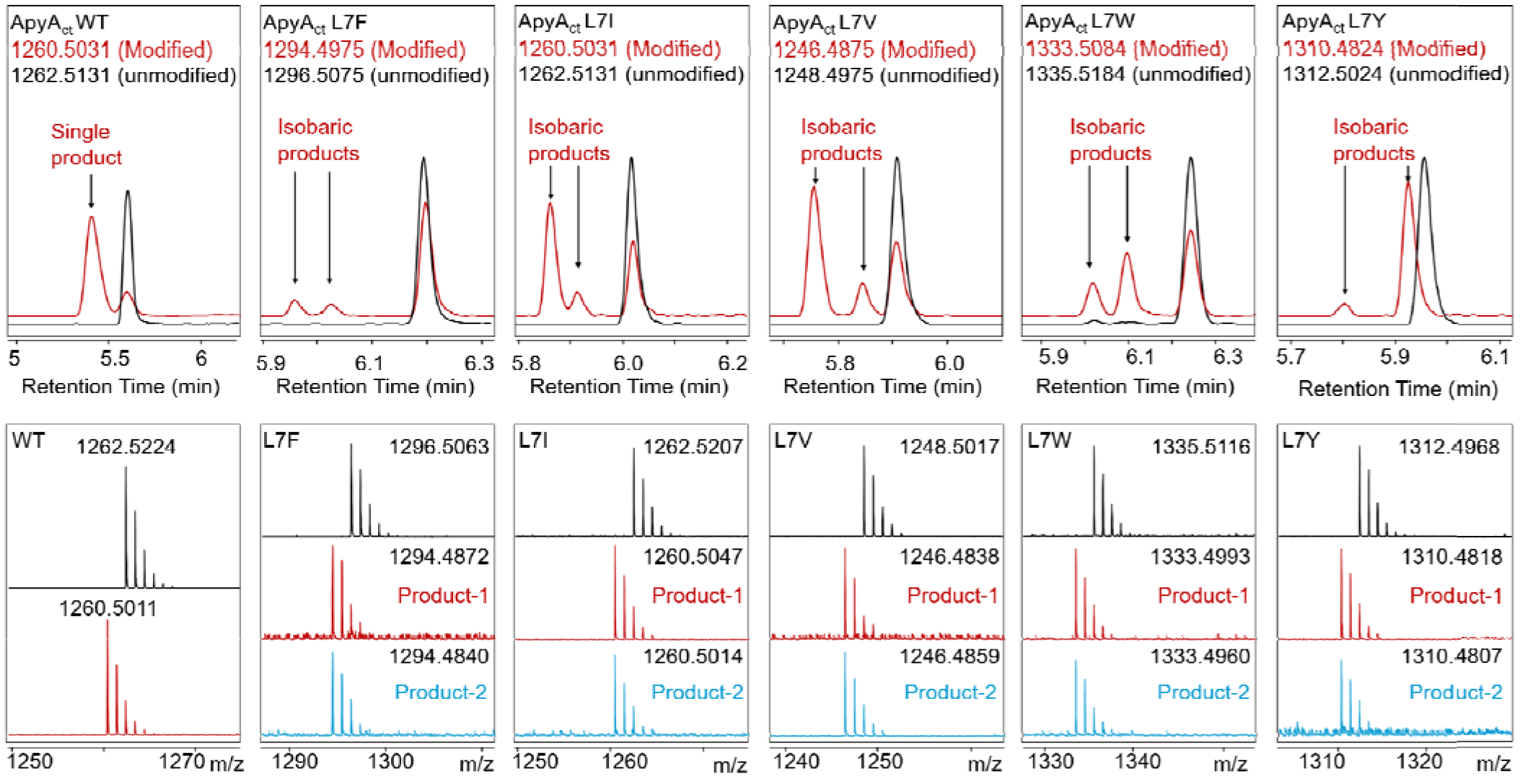
LC-HRMS analysis of variant ApyA_ct_ peptides reacted with ApyO. (Top) EICs of unmodified (black trace) and ApyO-modified (pink trace) ApyA_ct_ wildtype (WT) and Leu7 variant peptides. Calculated masses are shown in the top panel for *N*−formylated methionine-containing peptides (resulting from PURExpress in vitro translation) for both substrates and products. Two isobaric products were observed for L7F, L7I, L7V, L7W and L7Y variants. (Bottom) Corresponding HRMS spectra for the WT and each variant. The unreacted substrate (no enzyme control) is shown in black while the isobaric products (ApyO-modified leading to 2 Da mass loss, characteristics of macrocyclization) are shown in red and blue for products 1 and 2, respectively. For additional variants that underwent cyclization but did not form isobaric products or the mutants that were not modified, see the Supporting Information. Calculated monoisotopic masses [M+H]^+^ used for EICs are shown in the chromatogram (top) and the corresponding observed masses are annotated above the spectra (bottom).

In the AlphaFold model, the Leu in ApyA in between the two modified Tyr residues (Leu6) occupies a hydrophobic pocket formed by Phe181, Val260, Pro265 and Ile427 of ApyO (Figure 4C), and Met2 is embedded in a hydrophobic pocket formed by Phe18, Phe45, Leu323 and Ile425 (Figure 4D). In orthologous precursor peptides, position 2 is predominantly occupied by the hydrophobic residues Leu (45%), Met (21%), Ile (17%), Ala (7%), Val (5%) and Gly (4%). Additionally, the model predicts Glu4 of ApyA_ct_ to be in proximity to Arg47 of ApyO (Figure 4E). A negatively charged amino acid is less prevalent at position 4 than the positively charged Arg/Lys at position 1 (26% Glu, 42% Asp), but when the precursor peptide contains an Asp/Glu at this position, the corresponding ApyO homologs almost always contain an Arg residue at the equivalent position of Arg47 (Figure S9). Hence, this predicted substrate-enzyme interaction may also be important for activity.

### Site directed mutagenesis of ApyA

To experimentally test the AlphaFold model, we investigated the substrate tolerance of ApyO with respect to several of the residues in ApyA_ct_ using the CFE assay. The wild type (WT) ApyA_ct_ sequence underwent about 70% conversion based on the area under the curve (AUC) of the corresponding EIC derived from HR LC-MS analysis (Figure 5). Replacement of Arg1 with any of the 19 proteinogenic amino acids proved deleterious to ApyO catalysis (Figure S10). Low activity was observed for the R1K variant (~8% conversion). Replacement of Leu7 was less detrimental with several amino acids showing good activity (Figure S11). Conversion was highest for the L7Y variant (near quantitative, Figure 5), with lower conversions for L7I (~60%), L7V (~57%), L7W (~39%), L7F (~8%), L7T (~3%) and L7G (~1%) as determined by LCMS (Figure 5 and S11, S12). Data for the variants that did not undergo modification are shown in Figures S13-S15. Replacing Glu4 with an Ala in ApyA diminished product formation by ApyO (~56%) whereas a Lys substitution proved deleterious (~23% conversion) (Figure S16A,B). Inspired by the biarylitides where crosslinks have been reported between Tyr and His residues [18, 34], we also utilized the CFE setup to test conversion of Y6H, Y8H and Y6/8H variants of ApyA_ct_ (Figure S16C-E). Only the Y8H variant resulted in a 2 Da mass loss upon reaction with ApyO suggesting a crosslinked product. Since this would not be a natural crosslink based on the substrate sequences in Figure S1, we did not determine its structure.

### Identification of two isobaric products from select mutant substrates

The LC-MS analysis for certain mutants (L7Y, L7W, L7F, L7I and L7V) showed two isobaric products eluting at different retention times (Figure 5), whereas other mutants such as L7T and L7G resulted in only one product peak (Figure S12). This observation suggested that different isomers were formed by ApyO upon substituting the residue in between the Tyr residues with specific amino acids. These could be stereoisomers or constitutional isomers. We selected the L7Y and L7W variants because of the relatively high amounts of the putative isomers that were produced in the CFE experiments. Full length His-tagged ApyA-L7Y and -L7W precursors were co-expressed with ApyO in *E. coli* BL21 (DE3) TUNER and purified by metal affinity chromatography. Following GluC digestion, the relevant macrocyclic fragments were analyzed by LC-MS and tandem MS. The lack of fragmentation in the YYY- and YWY-motifs (Figure S17) in HR-MS/MS analysis suggested that the crosslinks were still present between the Tyr6 and Tyr8 residues. Both peaks of the L7Y and L7W products were purified by HPLC (Figure S18). For the L7W variant, the ratio of the two isobaric products was similar for the co-expression and the CFE experiments, but for the L7Y variant, the CFE experiment favored one of the isomers more than the co-expression conditions (compare Figures 5 and S18). These observations are not surprising since the full-length substrates are imperfectly mimicked by a truncated N-terminal formyl-Met containing peptide in the CFE experiments. Importantly, however, the key points gained from the CFE experiments were recapitulated in the co-expressions with full length substrates. These observations also support the hypothesis that the C-terminal 10 amino acids are the main determinant of the outcome of ApyO catalysis.

### Structural characterization of L7Y and L7W product isomers

The two isomers (denoted 1 or 2 based on LC elution order) of the pentapeptides originating from the ApyO-modified and GluC-digested ApyA-L7Y and ApyA-L7W peptides were next characterized by NMR spectroscopy (^1^H−^1^H TOCSY, ^1^H−^1^H NOESY, ^1^H−^13^CHSQC and ^1^H−^13^C HMBC). For the ApyA-L7Y isomer−1 and isomer−2 peptides, the spectroscopic measurements were conducted in 90% H_2_O, 10% D_2_O and 0.2% deuterated formic acid (dFA) at 45 ^°^C and 25 ^°^C, respectively (^1^H and ^13^C assignments are shown in Tables S4 and S5). For the ApyA-L7Y minor isomer−1, the aromatic protons of Tyr6 and Tyr8 exhibited different splitting patterns from a typical Tyr residue, integrating to three protons each and appearing as two doublet peaks and one singlet in the 1D ^1^H−^1^H TOCSY spectra (Figure S19). More detailed analysis of ^1^H−^13^C HSQC and HMBC (Figures S20 and S21, respectively) data revealed a C−C crosslink between the two aromatic rings of Tyr6 and Tyr8 at the Cε2-positions. This assignment was confirmed by the observation of a cross peak between the Hδ2 proton at 6.76 ppm of Tyr6 and the Cε2 carbon at 125.8 ppm of Tyr8. Thus, the crosslink in minor isomer 1 connects the same Tyr carbons as in the WT product. In contrast, the 1D TOCSY spectra of ApyA-L7Y major isomer−2 displayed three aromatic protons (two doublets and one singlet) for Tyr6, whereas four aromatic protons were observed for Tyr8 as doublets of doublets characteristic of a typical Tyr residue (Figure S22). The ^1^H and ^13^C chemical shifts (Figure S23), the cross peaks in the ^1^H−^13^C HMBC spectrum (Figure S24), and the observed NOE cross peaks (Figure S25), suggested a C−O crosslink between the Cε2 carbon of Tyr6 and the phenolic oxygen of Tyr8 (Figure 6 and Figure S24). Both structures of the constitutional isomers of the ApyA-L7Y product are consistent with the HR-MS and HR-MS/MS analysis shown in Figures 5 and S17, respectively.

**Figure 6.**
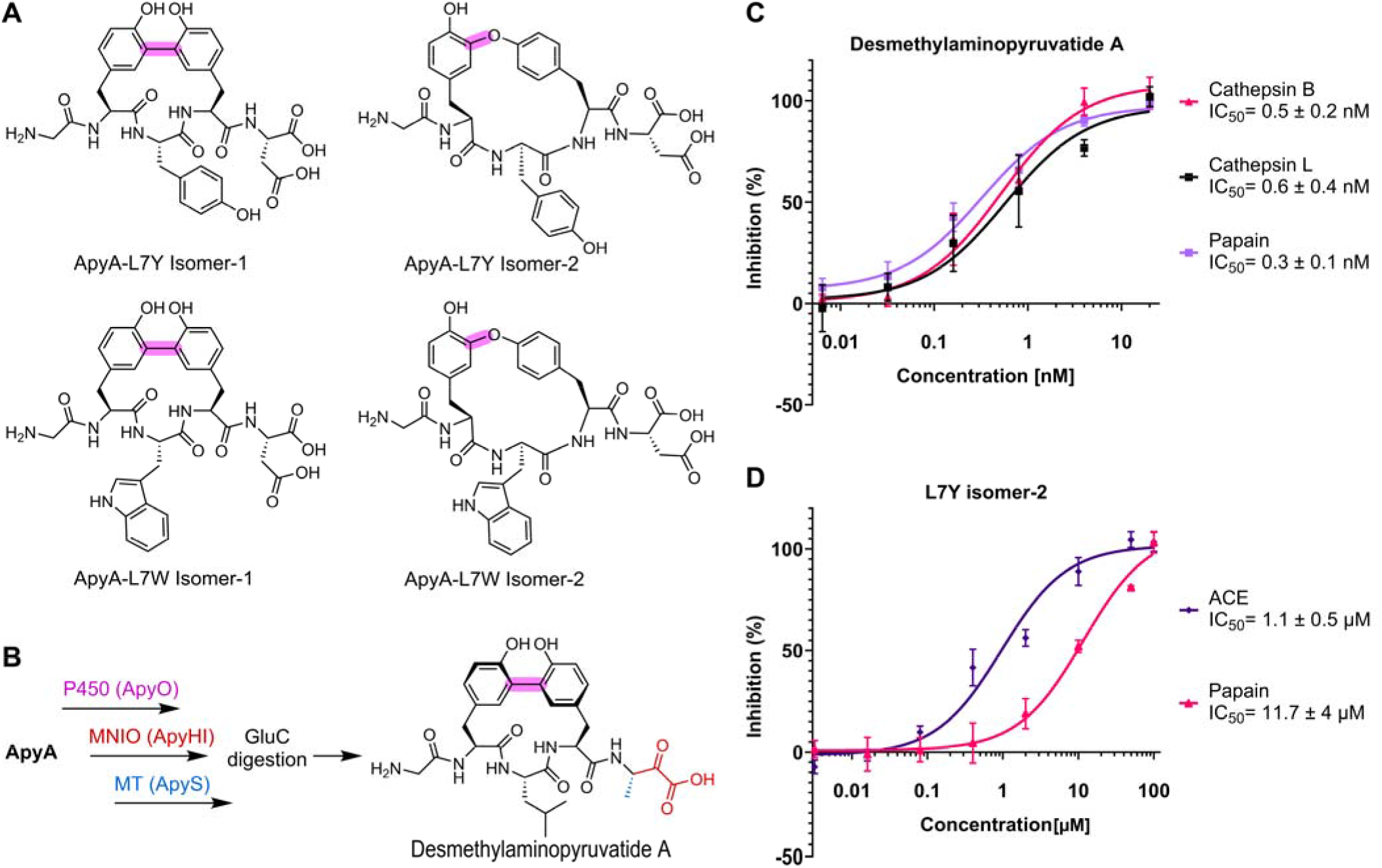
Chemical structures of the ApyO-modified L7Y and L7W mutants and protease inhibition. (A) Structures of ApyO-modified and GluC-digested ApyA-L7Y isomers 1 and 2 (top), and ApyA-L7W isomers 1 and 2 (bottom). (B) The structure of the desmethylaminopyruvatide A generated by modification of ApyA by the P450, MNIO and methyltransferase (MT) enzymes encoded in the *apy* BGC. The modified pentamer product was obtained by endoproteinase GluC digestion. (C) Non-linear regression plot showing the inhibition of cathepsin B, cathepsin L and papain by the desmethylaminopyruvatide A. (D) Non-linear regression plot showing the inhibition of ACE and papain by L7Y isomer-2.

Similarly, the two isomers of ApyO-modified ApyA-L7W (isomers −1 and −2) were analyzed. Their ^1^H and ^13^C chemical shift assignments are provided in Tables S6 and S7. Comparable to the ApyA-L7Y isomers, a C−C crosslink between the aromatic Cε2 positions of the two Tyr residues was observed for the ApyA-L7W isomer−1, whereas a C−O crosslink was identified for the L7W isomer−2 peptide (Figure 6A, and Figures S26-S32). The structures of both peptides are consistent with the HR-MS/MS results (Figure S17).

### Bioactivity

The bioactivity of the natural product produced by the *apy* BGC has not yet been reported. The final product contains an α-keto-β-amino acid moiety (Figure 2B), a structure (or the corresponding amides) that is present in known protease inhibitors [35]. Therefore, we tested wild type ApyA that was modified by the MNIO (ApyHI), P450 (ApyO) and methyltransferase (ApyS) enzymes (Figure 6B). The product lacks the β-methyl group that the cobalamin-dependent radical SAM enzyme ApyD installs on the β-carbon of Tyr8 [25], and is therefore designated desmethylaminopyruvatide A. This congener was selected because it shares the desmethyl scaffold of the ApyO-derived variant peptides evaluated below, allowing the contributions of the α-keto-β-amino acid group and the macrocycle to be compared directly across compounds. Whether the β-methyl group modulates potency was not determined here. The purified peptide was digested by endoproteinase GluC to afford the pentameric desmethylaminopyruvatide A (Figure 6B). The compound inhibited cathepsin L, cathepsin B and papain with IC_50_ values of 0.6 nM, 0.5 nM and 0.3 nM, respectively (Figure 6C). No inhibition of trypsin, chymotrypsin and M^pro^ was observed, suggesting specificity towards the papain family of Cys proteases.

Although the α-keto-β-amino acid moiety in desmethylaminopyruvatide A is the likely pharmacophore for the observed inhibition, compounds containing only ether-linked biaryl scaffolds such as OF4949 and K-13 (Figure 7) are also known protease inhibitors [36, 37]. Therefore, the GluC-digested pentameric peptides originating from ApyO catalysis on the ApyA variants Y6W, Y8W, L7Y (both isomers) and L7W (both isomers) were also tested at 25 µM against an array of proteases including Ser proteases like trypsin and chymotrypsin, Cys proteases like papain, cathepsin B and cathepsin L, the zinc-dependent metalloprotease angiotensin I-converting enzyme (ACE) and the main protease from SARS-CoV-2 virus (M^pro^). No inhibition was observed with the exception of L7Y isomer-2, which inhibited ACE and papain with an IC_50_ of 1.1 µM and 11.7 µM, respectively (Figure 6D). L7W isomer-2 exhibited weak inhibition of ACE (~40 %) at 25 µM. Therefore, the IC_50_ value of L7W isomer-2 was not determined. Compared to desmethylaminopyruvatide A, these activities were modest suggesting that the α-keto-β-amino acid moiety is critical for its observed activity.

**Figure 7.**
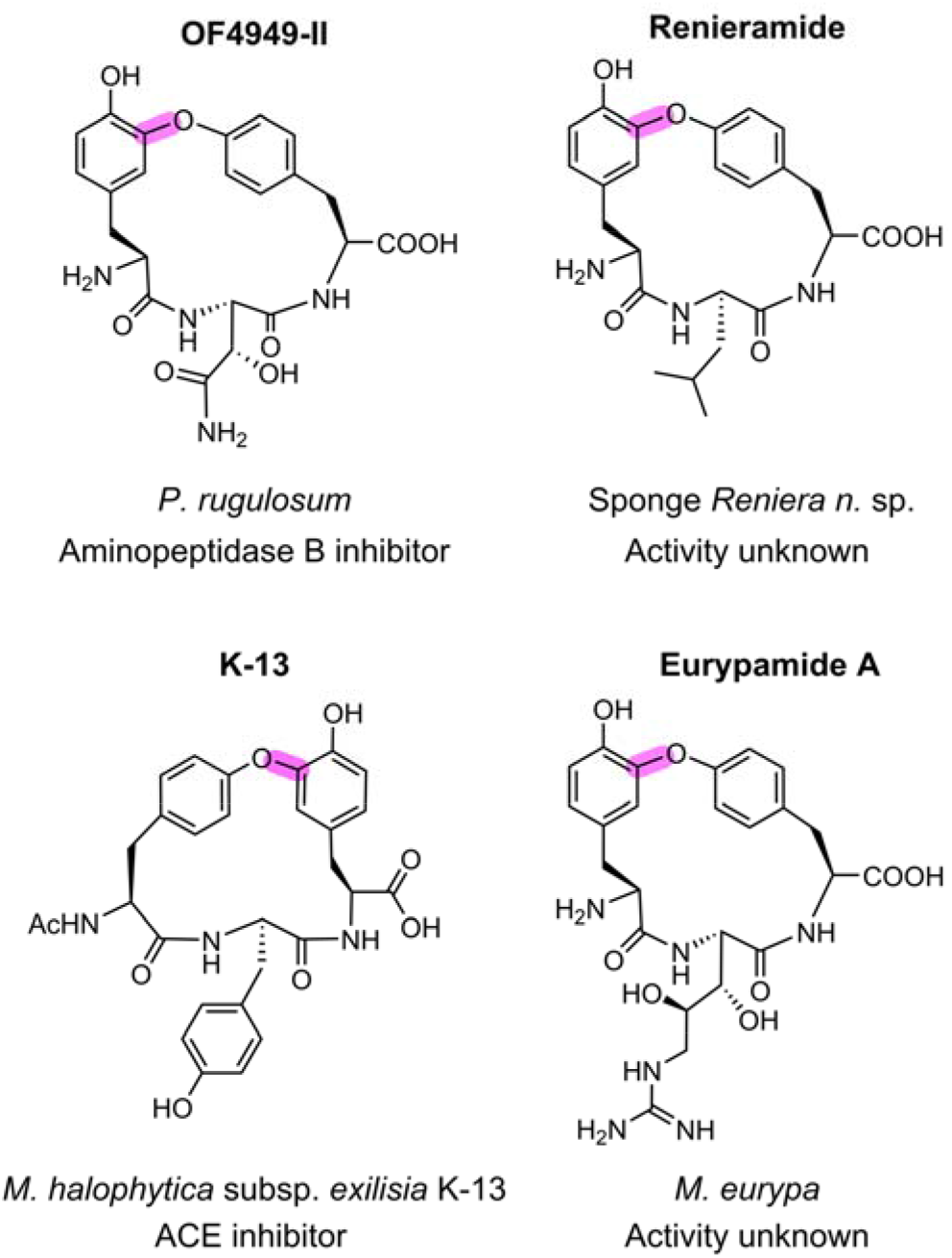
Examples of ether-linked tyrosines in tripeptide scaffolds. The aminopeptidase B inhibitor OF4949 and the ACE inhibitor K-13 contain a C−O linkage. Similar macrocyclic scaffolds are present in renieramide and eurypamide, however their bioactivities are unknown.

We therefore also evaluated whether the L7Y variant of ApyA would be a substrate for subsequent modifications by the other tailoring enzymes encoded in the *apy* BGC in the hope to convert the C−O crosslinked peptide into a product with an α-keto-β-amino acid. Co-expression of ApyA-L7Y with ApyOHIDS in *E. coli* resulted in inefficient conversion producing many incompletely modified intermediates and with α-keto-β-amino acid containing peptides as minor products (Figure S33). Co-expression of the L7Y variant and ApyOHIDS in *Burkholderia* sp. FERM BP-3421 that was used for the generation of aminopyruvatides in our previous study [26] also resulted in multiple products with keto acid-containing peptides as minor components (Figures S34-S35). Because of the low yield and mixture of compounds obtained, we were not able to test whether an α-keto-β-amino acid containing constitutional isomer with a non-native crosslink inhibited proteases.

### Comparison with Other RiPP Systems

In RiPP biosynthesis, the leader peptide often plays a significant role in substrate recognition by the class-defining maturases, often requiring a RiPP recognition element as a fused domain or a separate protein [29, 38–40]. P450 enzymes in RiPP pathways generally seem to be less dependent on remote binding interactions. For instance, P450_Blt_ and its engineered variants require only a short peptide for crosslinking [20, 41–44]. Genome mining studies have revealed the widespread distribution of over 400 short ORFs that encode only five residues co-occurring with P450s throughout the bacterial kingdom [19]. In addition, a P450 was recently reported from the typertide BGC that acts in a leader-independent manner to form C−C, C−O and C−N linkages in tetrapeptide substrates [45]. Consistent with these previous reports, in this study, we utilized AlphaFold 3 modelling of ApyO with ApyA along with CFE to identify a short 10-mer substrate peptide. Analysis of ApyO activity on variants of this 10-mer peptide showed that an Arg near the N-terminus of this peptide is required for crosslinking of the C-terminal Tyr-Leu-Tyr motif. In contrast, substitution of the Leu with select residues was tolerated, with hydrophobic side chains resulting in promising levels of crosslinked products. Surprisingly, several substrates that were successfully crosslinked produced two isobaric products. NMR characterization showed that these different products were not stereoisomers nor peptides that were crosslinked at different residues, but rather constitutional isomers in which the Tyr-Tyr crosslinks involved C−O rather than C−C bond formation. Our findings reinforce the importance of full structural elucidation of the products of such variants to evaluate whether the native crosslink is still generated [18, 43], even when the crosslinked residues themselves were not substituted. At present, the origin of the different crosslinking processes is unclear, but probably involves subtle differences in the spatial arrangement of the two Tyr residues when the Leu in the YLY motif is substituted.

Based on the AlphaFold predictions and CFE experiments, additional side chain interactions upstream of the YLY motif may play a role in substrate recognition with several hydrophobic side chains of residues on ApyA occupying hydrophobic pockets on ApyO, and Glu4 making an electrostatic contact with an Arg in the substrate binding site of ApyO. Mechanistically, P450-mediated biaryl crosslinking involving Tyr residues has been proposed previously to proceed through a high-valent Fe(IV)-oxo-porphyrin radical cation (Compound I) intermediate that abstracts a hydrogen atom forming a tyrosyl radical and Compound II [46–48]. This Tyr radical could crosslink with the second Tyr followed by electron transfer and tautomerization to provide the product (Figure S36). Alternatively, it has been suggested that Compound II formed from the first step could abstract another hydrogen atom from the second Tyr followed by radical-radical coupling [46–48] (Figure S37). Mechanistic studies will be required to provide additional insights.

Discovery of macrocyclic tripeptide molecules with crosslinked aromatic residues can be traced back to the 1980s before the genomic era. Recently, we deorphanized the biosynthesis of the biphenomycin class of biaryl crosslinked antibiotics [39]. The biosynthetic pathway of the aminopeptidase B inhibitor OF4949 from the fungus *P. rugolosum* OF4949 (Figure 7) was also recently reported [36, 49]. Like isomer-2 of the ApyO-product generated from the Leu7Tyr variant of the C-terminal peptide of ApyA, OF4949 contains a C−O crosslink between two Tyr residues in a tripeptide sequence. Structures with related crosslinks for which the biosynthesis has not yet been resolved include renieramide isolated from the sponge *Reniera* sp. [50], eurypamide from the sponge *M. eurypa* [51] and K-13 from *M. halophytica* subsp. *exilisia* K-13 (an ACE inhibitor) (Figure 7) [37]. A subset of these compounds inhibit proteases, but the ApyO-crosslinked peptides generated in this work did not display inhibition against proteases we tested except modest activity of the L7Y isomer-2 against ACE and papain. Isomer-2 of the L7Y variant shares structural resemblance with the ACE specific inhibitor K-13. However, unlike isomer-2, the oxygen atom in the C−O linkage of K-13 is derived from the N-terminal Tyr. The genome of the K-13 producer organism *M. halophytica* subsp. *exilisia* K-13 is not available, and therefore, its biosynthesis remains unexplored. Previous approaches for production of the 17-membered macrocycle in K-13 employed chemical synthesis using various strategies such as arene-ruthenium chemistry or macrocyclization by cycloetherification [52–54]. This study shows the possibility to access a K-13-like scaffold using an enzymatic approach.

The desmethylaminopyruvatide A containing the macrocyclic scaffold in addition to the C-terminal α-keto acid showed nanomolar activity against Cys-proteases demonstrating the potency imparted by the α-keto acid group. Cathepsins L and B are lysosomal Cys proteases that hold clinical significance. Their upregulation has been implicated in cancer metastasis, and their suppression reduces the invasiveness of aggressive tumor cells [55, 56]. Cathepsins are also implicated in viral entry of clinical pathogens including SARS-CoV-2 and their inhibition has demonstrated reduced viral entry and decreased gene copy numbers in host cells [57–59]. α-Keto acid/amide moieties are privileged motifs that have been employed in targeting Ser- and Cys-proteases as well as in antiviral drug discovery [35]. The presence of two privileged moieties in the aminopyruvatide class of compounds as well as the promiscuity of the crosslinking maturase ApyO provides a potential route for compound diversification for protease inhibition.

## Conclusion

The P450 ApyO encoded in the *apy* BGC from *B. thailandensis* E264 accepts a C-terminal 10-mer of the precursor ApyA. Arg1 in this peptide is required for ApyO-mediated crosslinking, and substitution of Tyr6 with a Trp yielded a N−C linkage between the indole Nε1 of Trp and Cε2 of Tyr8 instead of the Cε2−Cε2 linkage between Tyr6 and Tyr8 formed in the native substrate. Replacement of Leu7 with Tyr and Trp afforded two constitutional isomers for each variant that contained either the wild type Cε2−Cε2 linkage or a Cε2−O crosslink between Tyr6 and Tyr8. The desmethylpyruvatide A product from the *apy* biosynthetic pathway displayed subnanomolar inhibitory activity against clinically relevant Cys proteases through the combination of a 15-17 membered macrocyclic scaffold and an α-keto acid warhead.

## ACKNOWLEDGMENTS

We thank Prof. Angad P. Mehta for providing access to Agilent Synergy H1 plate reader and Andrea Dayal (laboratory of Prof. Satish Nair) for assistance with using a pipetting robot. This manuscript is the result of funding in part by the National Institutes of Health (R35 GM158411 to D.A.M.) and therefore it is subject to the NIH Public Access Policy. Through acceptance of this federal funding, NIH has been given a right to make this manuscript publicly available in PubMed Central upon the Official Date of Publication, as defined by NIH. A Bruker UltrafleXtreme mass spectrometer used was purchased with support from the Roy J. Carver Charitable Trust (Grant No. 22-5622). W.A.V. is an Investigator of the Howard Hughes Medical Institute and a Partner Investigator of the Australian Centre of Excellence for Innovations in Peptide and Protein Science.

## Data Availability Statement

The data that support the findings of this study are available at Padhi, Chandrashekhar; Zhu, L; van der Donk, Wilfred (2026), “Data associated with “Substrate-dependent crosslinking by the cytochrome P450 from aminopyruvatide biosynthesis”“, Mendeley Data, V1, doi: 10.17632/cr798k4bhw.1.

## Supporting Information

Additional supporting information can be found in the Supporting Information section.

**Supporting File:** Experimental procedures, Figures S1-S37 showing spectroscopic data, and Tables S1-S7 with gene sequences, primers and NMR annotations.

