## Supporting Information for "Substrate-Dependent Crosslinking by the Cytochrome P450 from Aminopyruvatide Biosynthesis"

Figure S29. 1D $^1H$-$^1H$ TOCSY spectra showing spin systems of the aromatic region of Tyr6, Trp7 and Tyr8 of the GluC-digested ApyO-modified ApyA-L7W isomer-2 in 90% $H_2O$, 10% $D_2O$, and 0.2% dFA. 30
Figure S30. 2D $^1H$-$^{13}C$ HSQC data showing the aromatic region of the GluC-digested ApyO-modified ApyA-L7W isomer-2 in 90% $H_2O$, 10% $D_2O$, and 0.2% dFA. 31
Figure S31. 2D $^1H$-$^{13}C$ HMBC data showing the aromatic region of the GluC-digested ApyO-modified ApyA-L7W isomer-2 in 90% $H_2O$, 10% $D_2O$, and 0.2% dFA. 32
Figure S32. 2D $^1H$-$^1H$ NOESY data showing the aromatic region of the GluC-digested ApyO-modified ApyA-L7W isomer-2 in 90% $H_2O$, 10 % $D_2O$, and 0.2% dFA. 33
Figure S33. LC-HRMS analysis of GluC-digested Apy-L7Y co-expressed with ApyOHIDS in *E. coli*. 34
Figure S34. LC-HRMS analysis of LysC-digested Apy-L7Y variant co-expressed with ApyOHIDS in *E. coli* and *Burkholderia* sp. 35
Figure S35. Tandem MS analysis of LysC-digested Apy-L7Y variant co-expressed with ApyOHIDS. 36
Figure S36. Possible mechanism for the two observed constitutional isomers formed from ApyO catalysis. 37
Figure S37. Alternative mechanisms proposed in the literature for Tyr-Tyr crosslink formation that involves radical-radical couplings 38

#### Supplementary Tables

Table S1. Plasmid constructs used in this study. 39
Table S2. NMR assignments of ApyO-modified, GluC-digested ApyA-Y6W 43
Table S3. Primers used for KLENOW fragment extension. 44
Table S4. NMR assignments of ApyO-modified, GluC-digested ApyA-L7Y Isomer–1 45
Table S5. NMR assignments of ApyO-modified, GluC-digested ApyA-L7Y Isomer–2 45
Table S6. NMR assignments of ApyO-modified, GluC-digested ApyA-L7W Isomer–1 46
Table S7. NMR assignments of ApyO-modified, GluC-digested ApyA-L7W Isomer–2 46

#### Materials and Methods

Plasmid constructs 47
Heterologous expression and purification of peptides 47
Isolation of matured C-terminal core peptide 48
HPLC purification of modified peptide core fragments 48
High-resolution tandem mass spectrometry 49
Expression and purification of the cytochrome P450 ApyO 49
Klenow fragment extension 50
*In vitro* assays 50
Kinetic experiments 51
NMR data acquisition and analysis 51
Protease inhibition assays 52

#### References

References 54

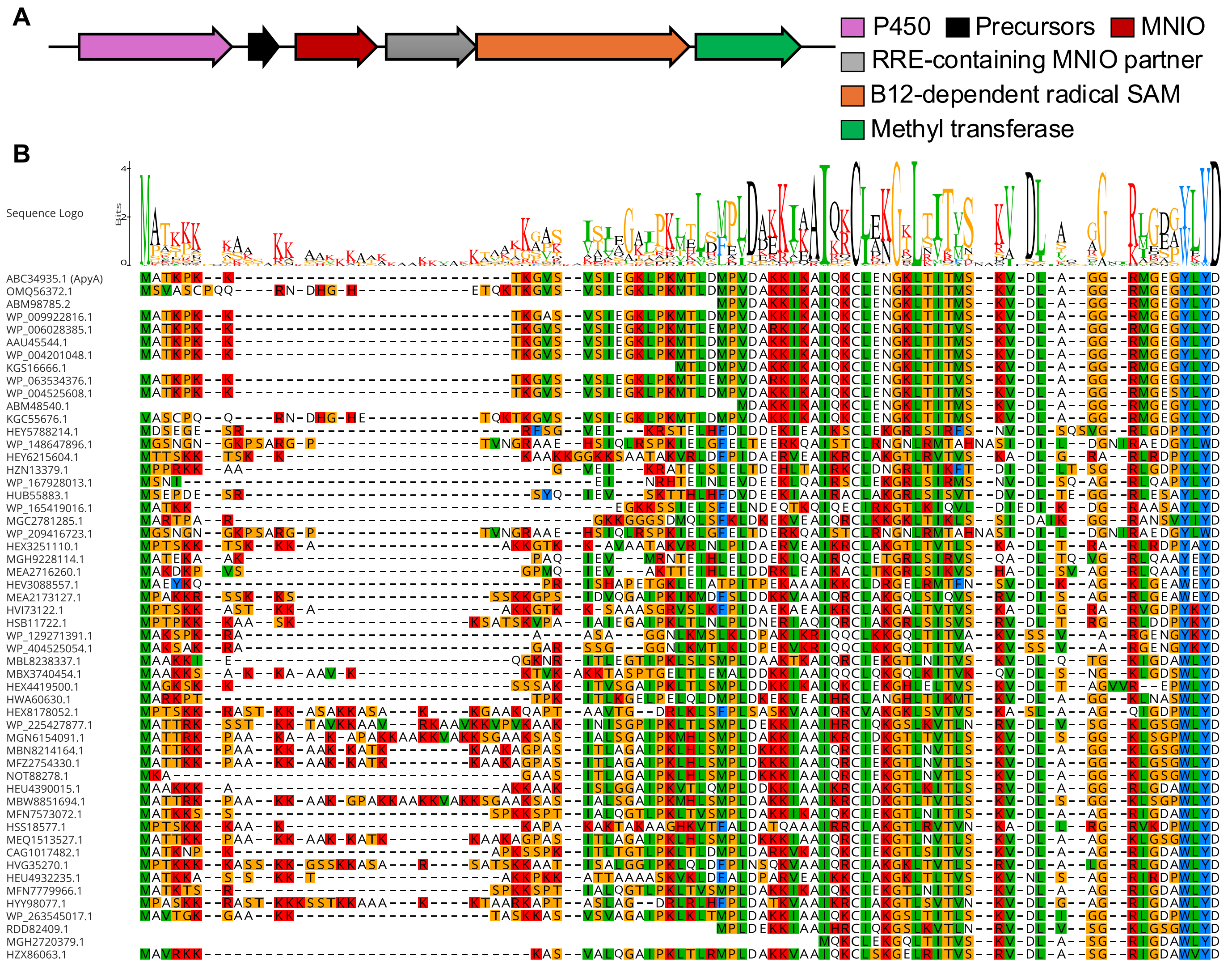

**Figure S1. BGC architecture and precursor sequence logo of orthologous clusters containing ApyO homologues.** (A) The core architecture of BGCs orthologous to the *apy* BGC. (B) Multiple sequence alignment of the precursor peptides (and their corresponding NCBI accession IDs) containing aromatic residues at the C-terminus with the resulting sequence logo. The BGCs encoding precursor peptides with the C-terminal aromatic residues predicted to participate in crosslink formation always encode a P450 enzyme homologous to ApyO. The accession number for the ApyA peptide that is the subject of this investigation is ABC34935.1.

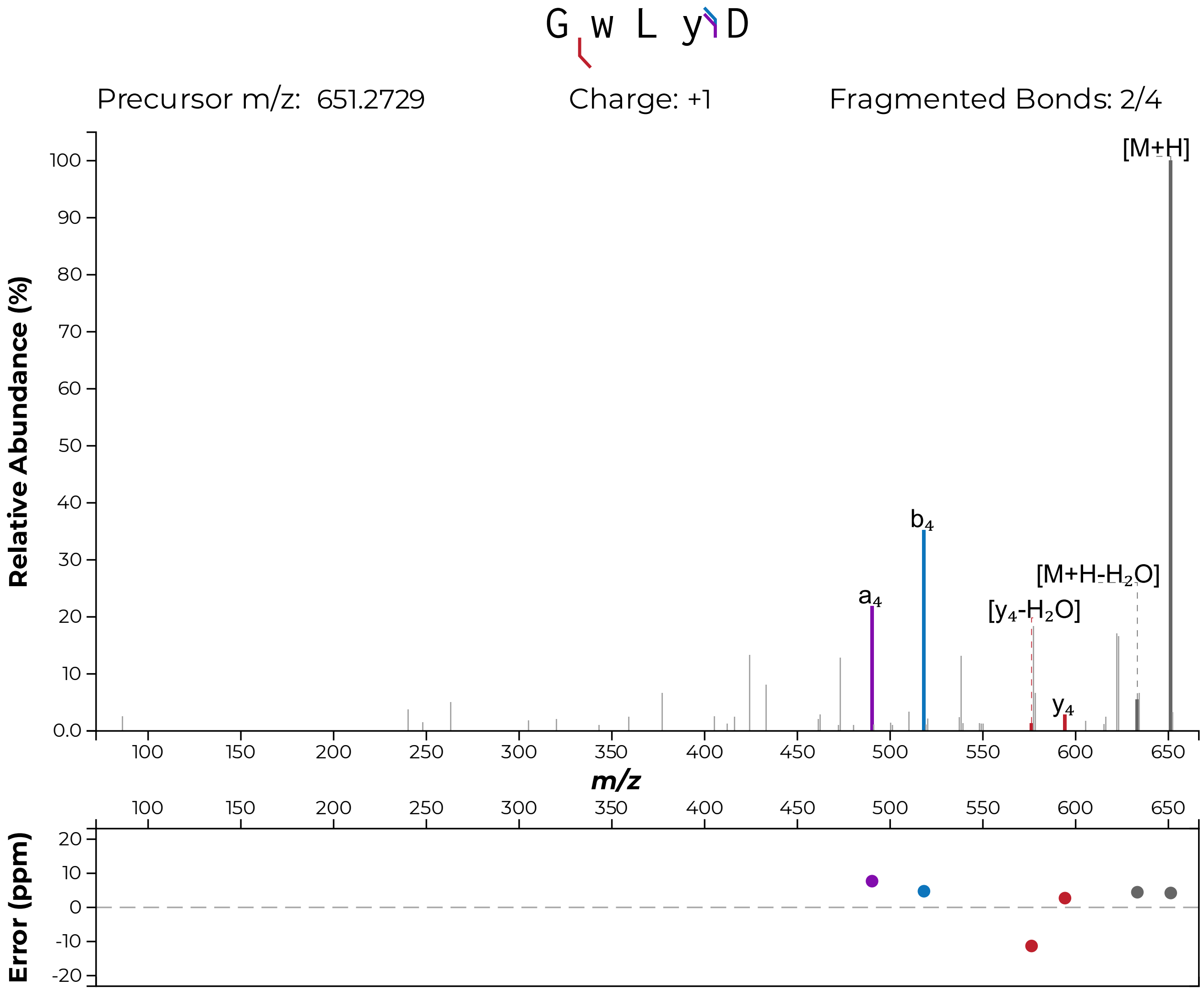

**Figure S2. Tandem MS analysis of the GluC-digested ApyA-Y6W mutant modified by ApyO.** HR-MS/MS analysis shows no fragmentation in the WLY motif suggesting a crosslink formed between Trp6 and Tyr8. For residue numbering, see Fig. 2. Fragment ion annotation was performed using the interactive peptide spectral annotator[1] with residues indicated in lower case w and y entered as dehydrogenated through a crosslink (M - 2 Da).

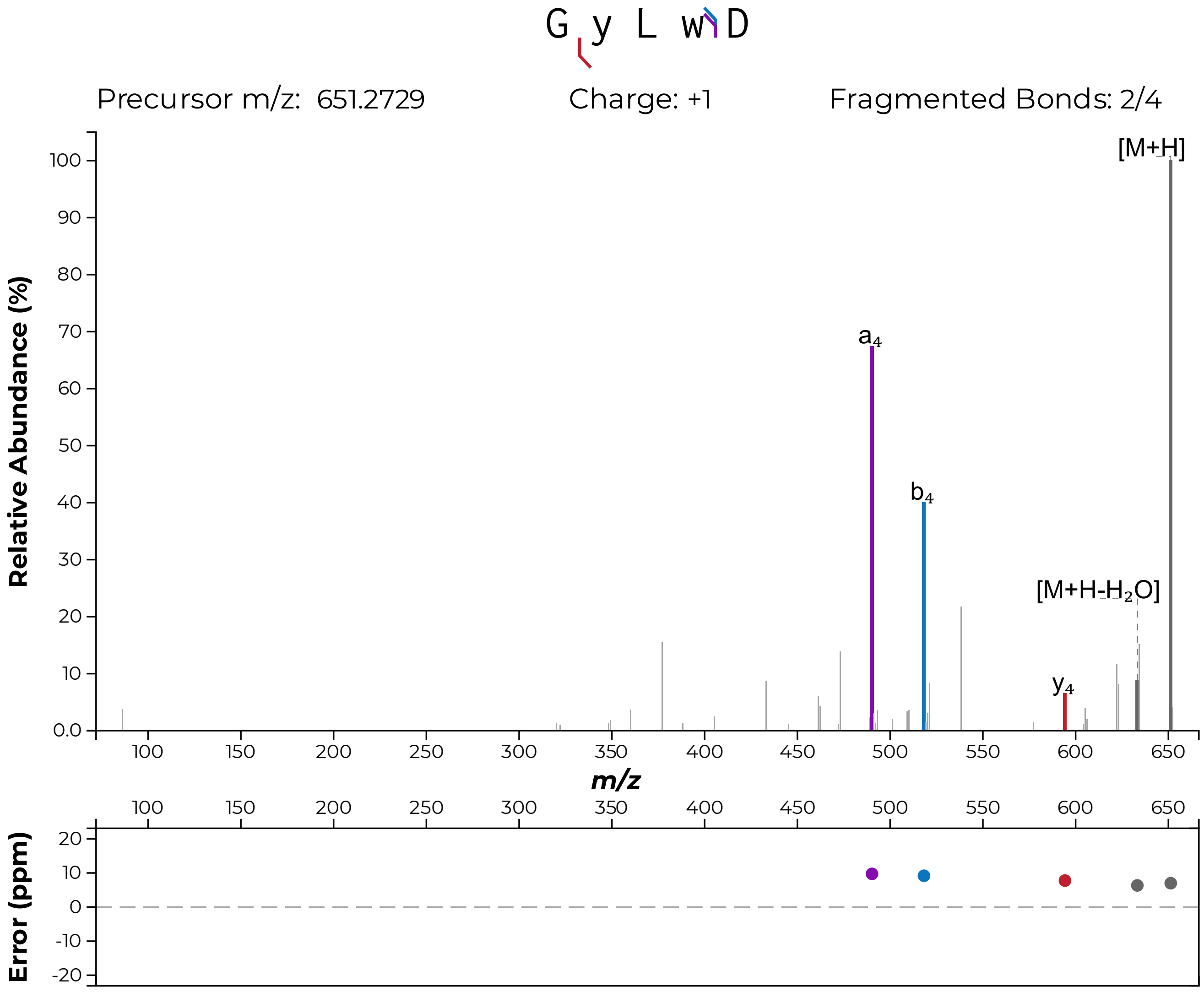

**Figure S3. Tandem MS analysis of the GluC-digested ApyA-Y8W mutant modified by ApyO.** HR-MS/MS analysis shows no fragmentation in the YLW motif suggesting a crosslink formed between Tyr6 and Trp8. For residue numbering, see Fig. 2. Fragment ion annotation was performed using the interactive peptide spectral annotator[1] with residues indicated in lower case w and y entered as dehydrogenated through a crosslink (M - 2 Da).

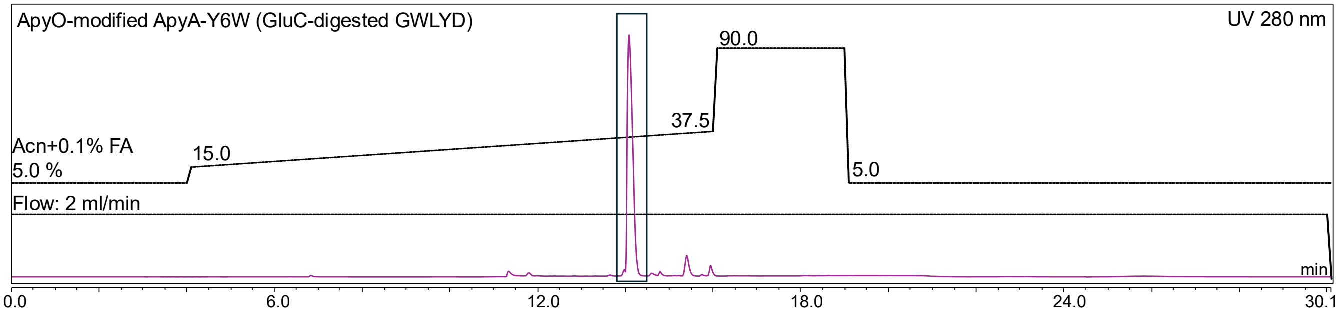

**Figure S4. HPLC purification of ApyO-modified GluC-digested ApyA-Y6W.** The C-terminal fragment was purified and is shown.

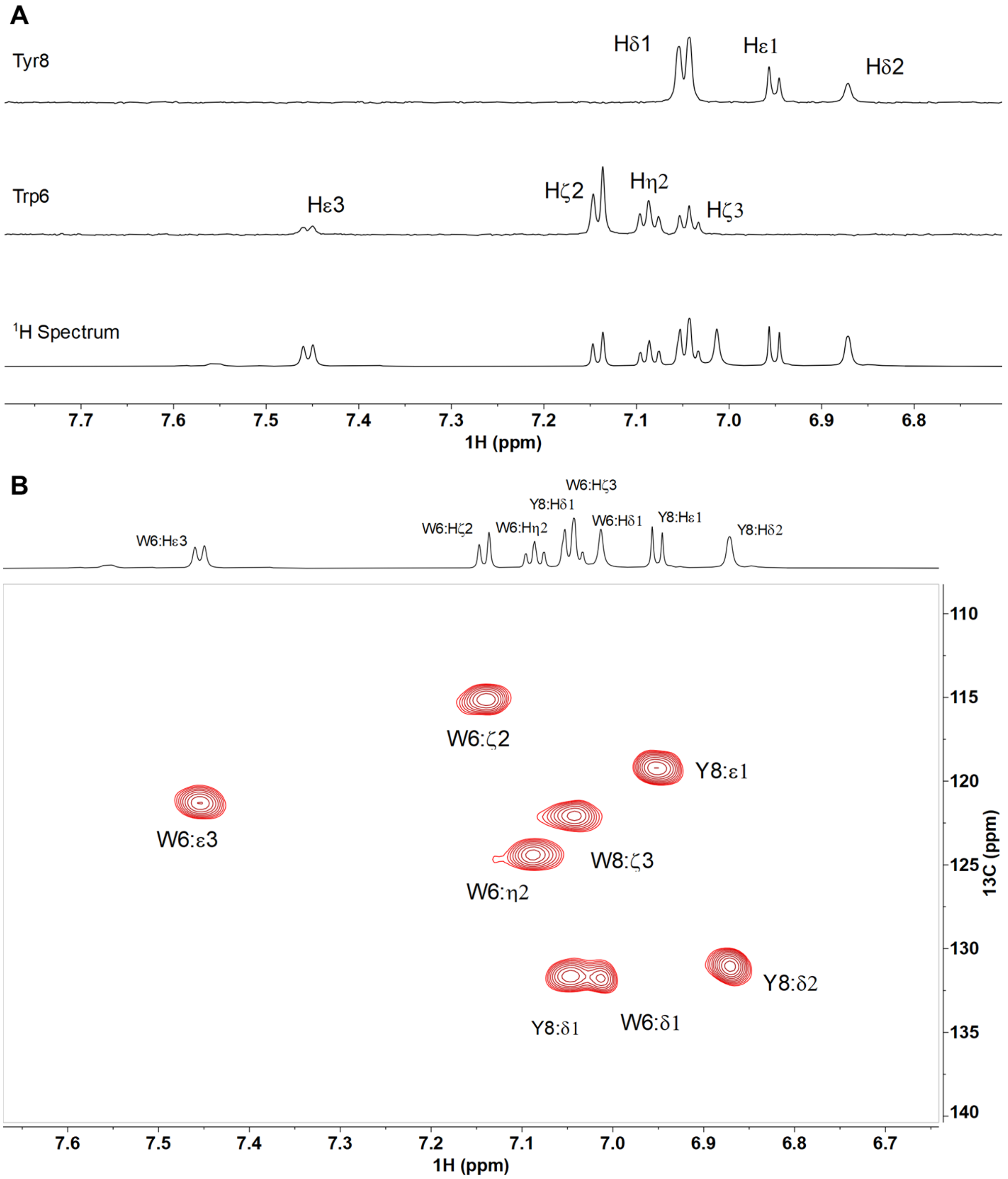

**Figure S5. 1D $^{1}H$-$^{1}H$ TOCSY and 2D $^{1}H$-$^{13}C$ HSQC analysis of the aromatic region of GluC-digested, ApyO-modified ApyA-Y6W in DMSO-$d_6$ containing 0.1% d-formic acid at 55 ºC.** (A) 1D $^{1}H$-$^{1}H$ TOCSY spectra showing the aromatic side chain protons of Trp6 and three aromatic side chain protons of Tyr8. The Hδ1 proton of Trp6 is visible in the $^{1}H$ NMR spectrum (bottom) but does not couple to other protons and therefore does not show up in the 1D TOCSY data (middle). (B) 2D $^{1}H$-$^{13}C$ HSQC data showing five and three aromatic proton-carbon cross peaks for Trp6 and Tyr8 residues are shown, respectively.

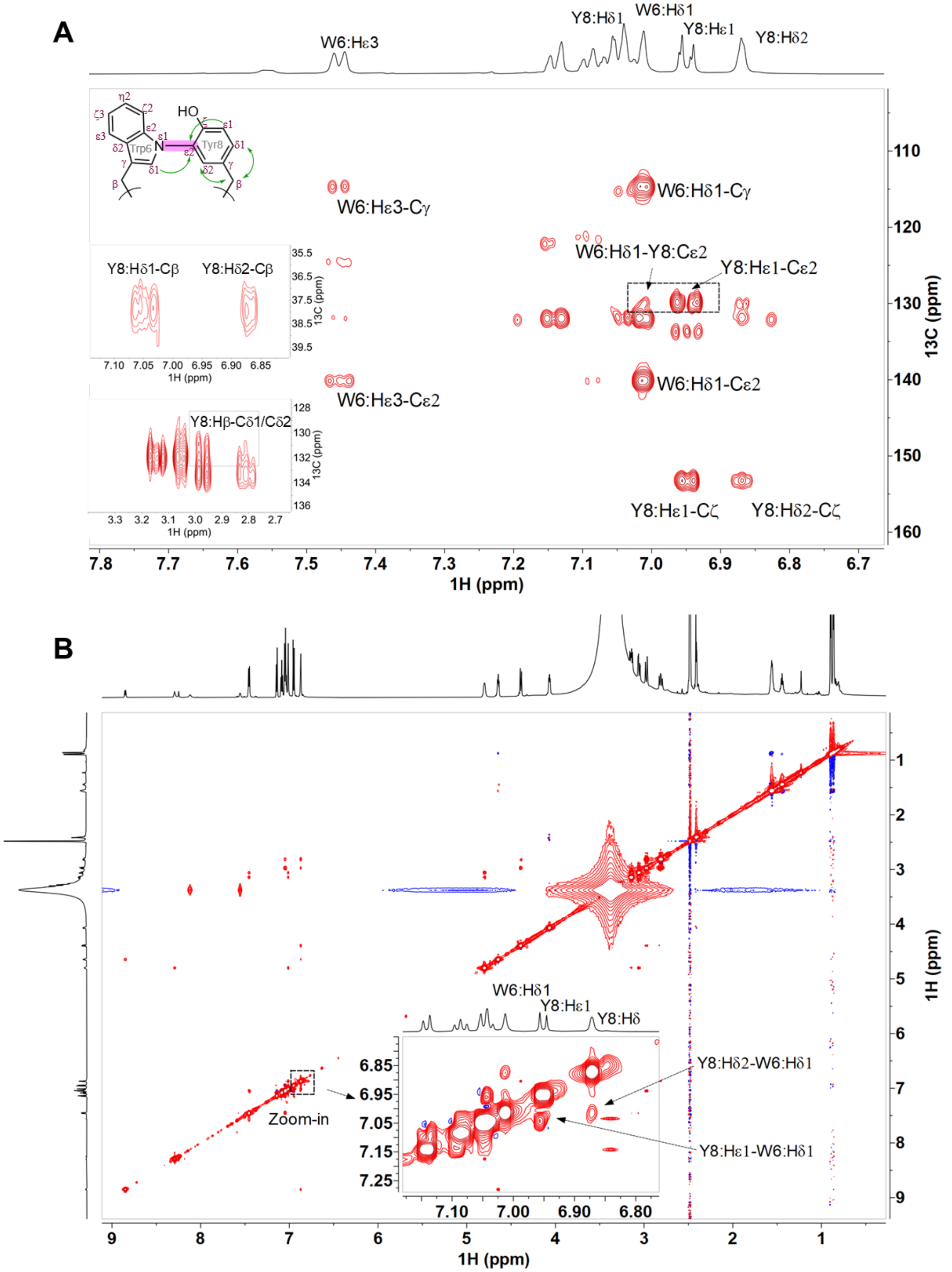

**Figure S6. 2D $^{1}$H-$^{13}$C HMBC of the aromatic region and 2D $^{1}$H-$^{1}$H NOESY (full spectrum) of GluC-digested, ApyO-modified ApyA-Y6W in DMSO-$d_6$ containing 0.1% d-formic acid at 55 ºC.** (A) Several key HMBC cross peaks are shown, notably the one between the two aromatic rings of Trp6 and Tyr8 residues. Insets are shown for the cross peaks observed for β and δ atoms. A $^{1}$H-$^{13}$C multiband J coupling constant ($J_{nxh}$) of 5 Hz was used in the HMBC experiment. (B) $^{1}$H-$^{1}$H NOESY spectrum showing a zoom-in of the aromatic region as an inset.

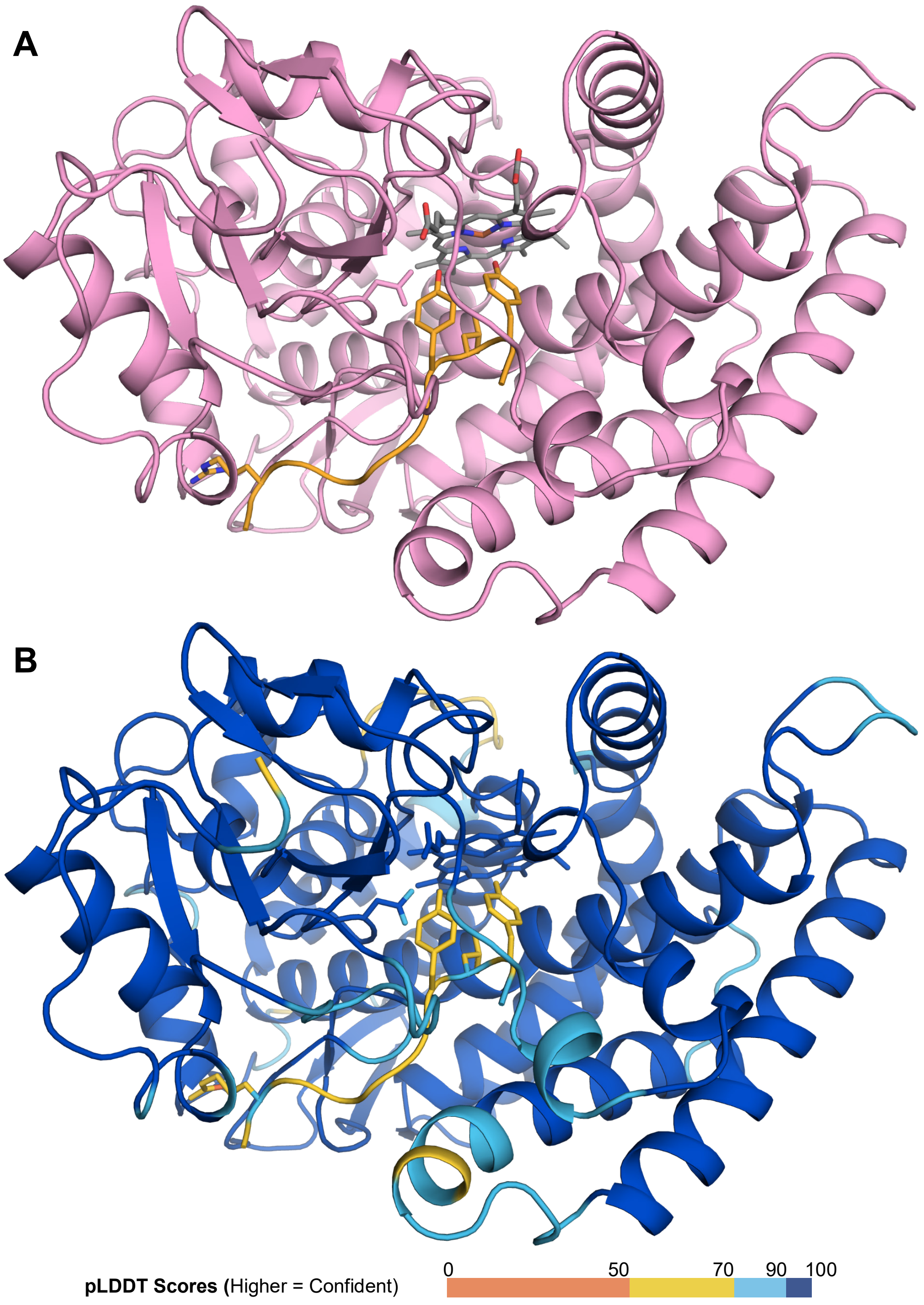

**Figure S7. AlphaFold3 model of ApyO in complex with ApyA$_{ct}$.** (A) AlphaFold-predicted complex of ApyO (pink) and the 10-mer ApyA$_{ct}$ (orange) showing the Y6 and Y8 residues modelled close to the active site heme. (B) Confidence metrics of the AlphaFold 3-predicted structures. pLDDT scores are shown as colored outputs (higher value means higher confidence as shown in the legend).

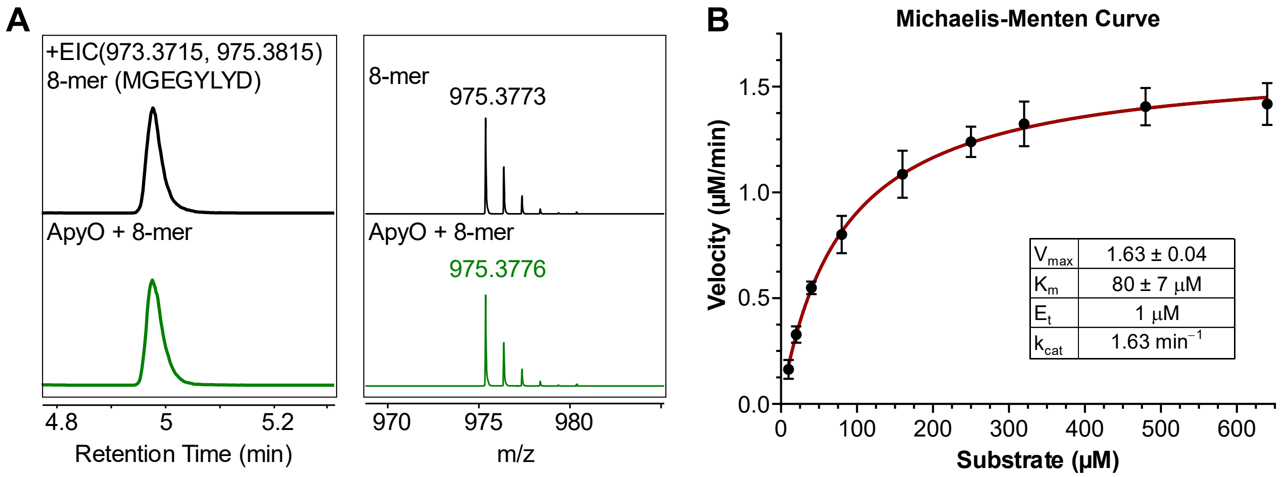

**Figure S8. LC-HRMS analysis of ApyA 8-mer and Michaelis-Menten curve for ApyO with the 10-mer peptide.** (A) Extracted ion chromatograms (EIC; left) of the in vitro reaction of a ApyA$_{ct}$ variant encompassing the C-terminal 8-mer peptide in the absence (black chromatogram) or presence (green chromatogram) of ApyO. The corresponding HRMS isotopic distribution (right) in the absence (black spectrum) or presence (green spectrum) of ApyO is displayed for the $[M+H]^+$ ion. (B) Michaelis-Menten curve showing the initial reaction velocity of ApyO in response to the substrate (*N*-acetylated-GRMGEGYLYD) concentration.

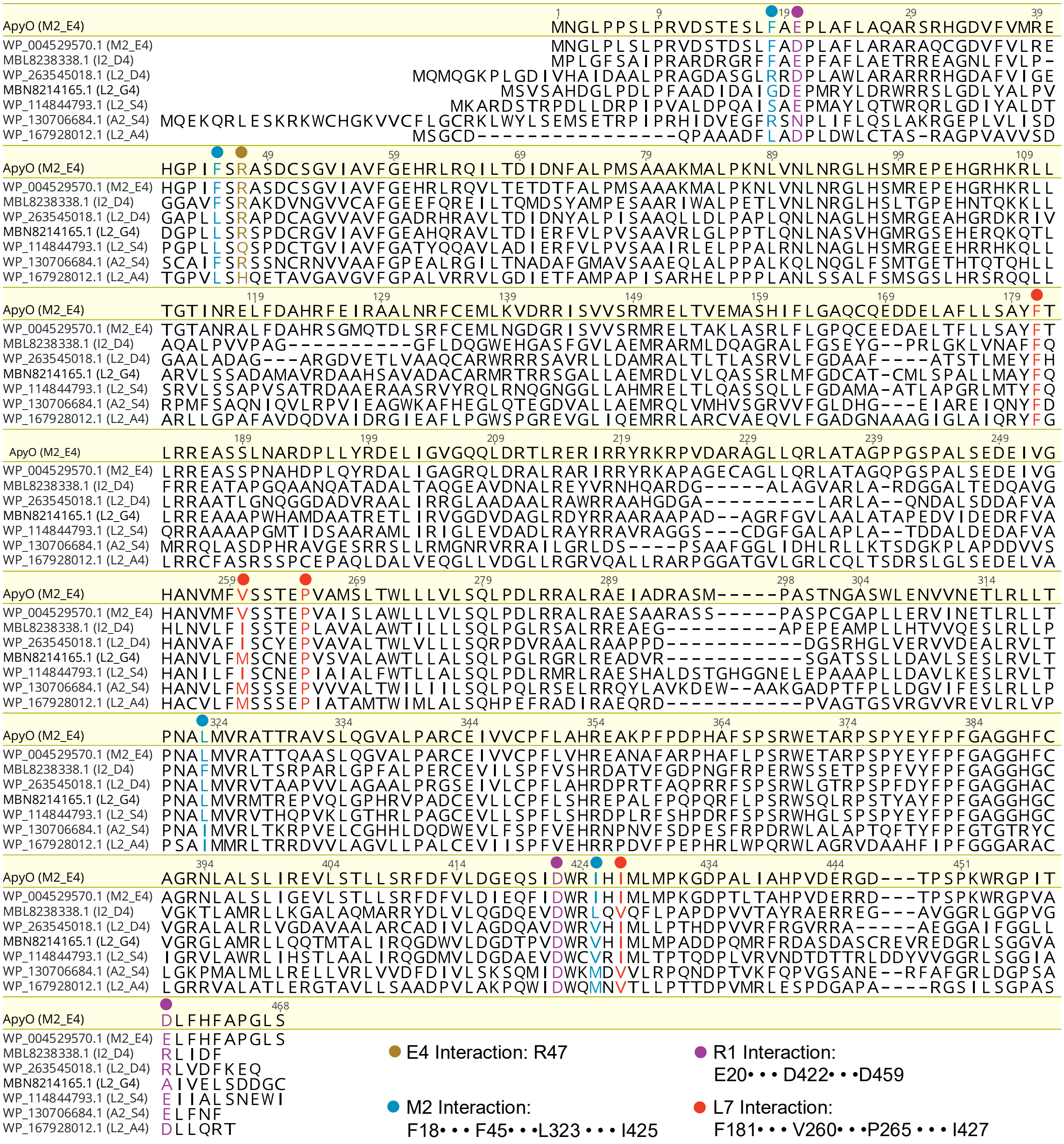

**Figure S9. Sequence alignment of ApyO homologues.** (A) Multiple sequence alignment of select ApyO homologs (NCBI accession IDs) from orthologous BGCs encoding ApyA variants with alternate residues at the second and fourth positions (in brackets). ApyO residues interacting with the R1, M2, E4 and L7 residues of ApyA$_{ct}$ predicted by AlphaFold are color-coded.

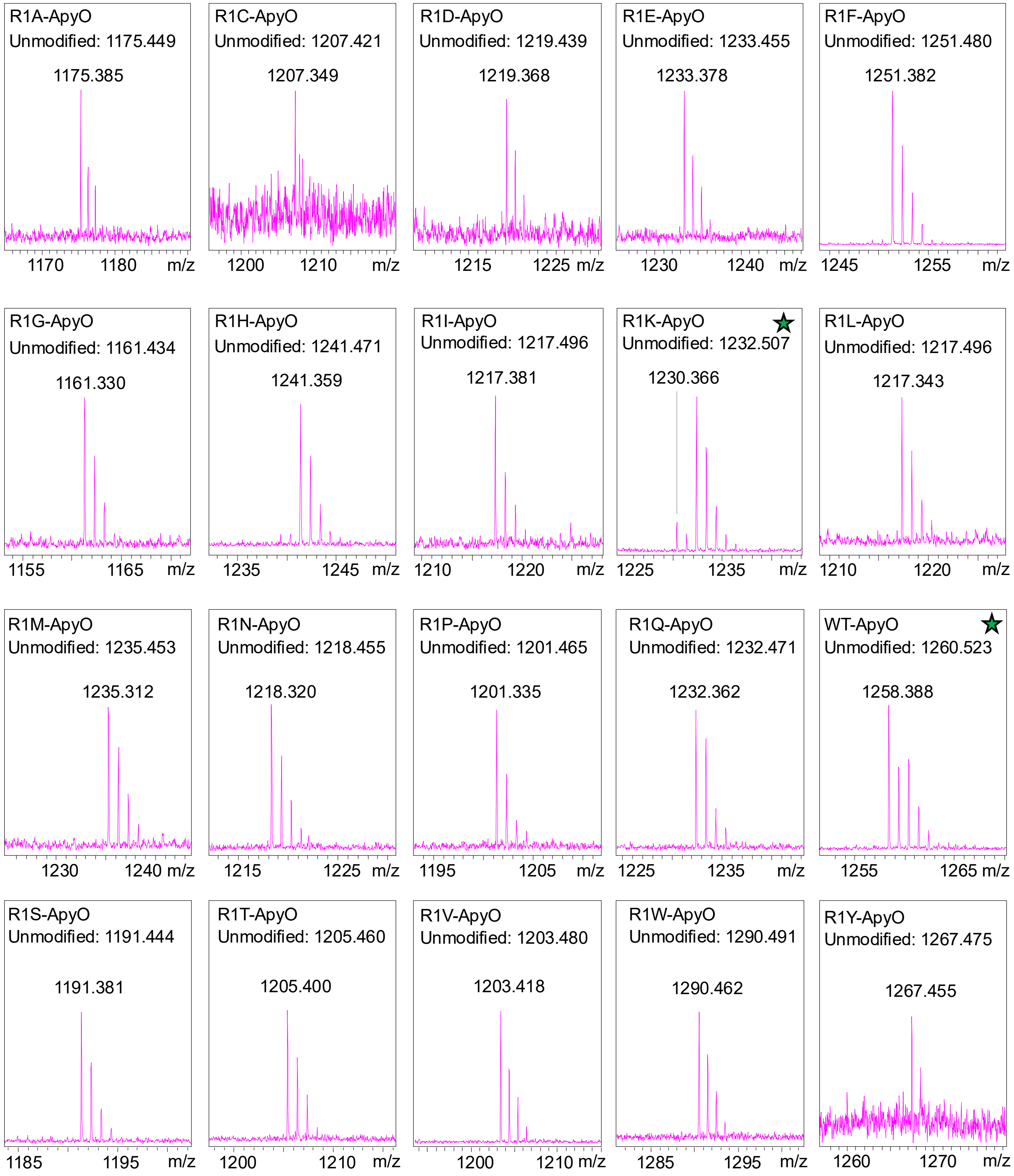

**Figure S10. MALDI-ToF mass spectra of ApyA$_{ct}$-Arg1 variants.** Zoomed-in mass spectra of the ApyA$_{ct}$ mutants where Arg1 was replaced and reacted with ApyO in vitro. Calculated masses for the unmodified substrates are provided for each spectra as $[M–H]^{-}$ ions. The monoisotopic $[M–H]^{-}$ masses are annotated above the observed spectra. Star indicates the only variant that underwent some level of modification.

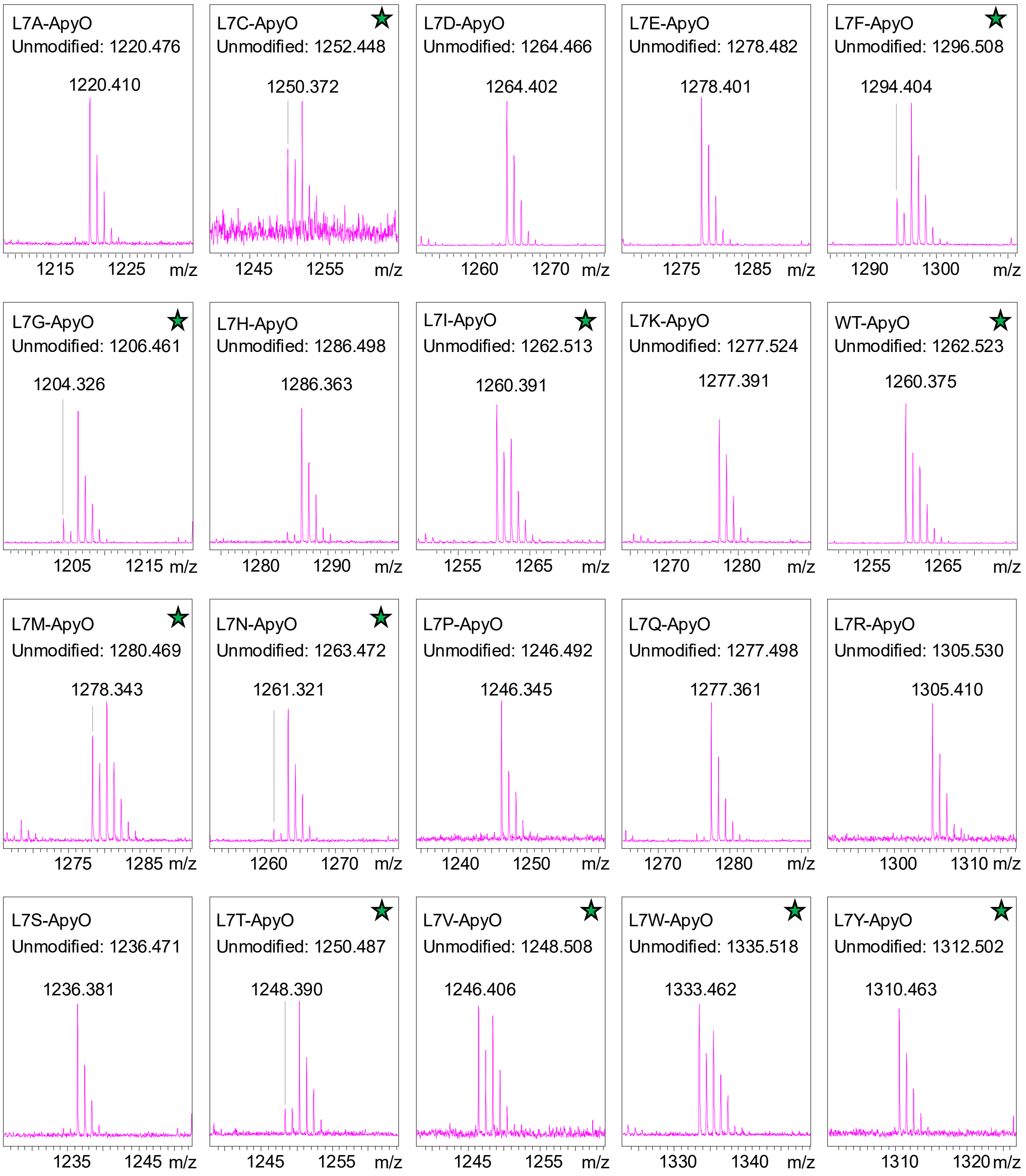

**Figure S11. MALDI-ToF mass spectra of ApyA$_{ct}$-Leu7 variants.** Zoomed-in mass spectra of the ApyA$_{ct}$ mutants where Leu7 was replaced and reacted in vitro with ApyO. Calculated masses for the unmodified substrates are provided for each spectra as $[M+H]^+$ ions. The monoisotopic $[M+H]^+$ masses are annotated above the observed spectra. Star represents variants or the wildtype (control) that underwent some level of modification.

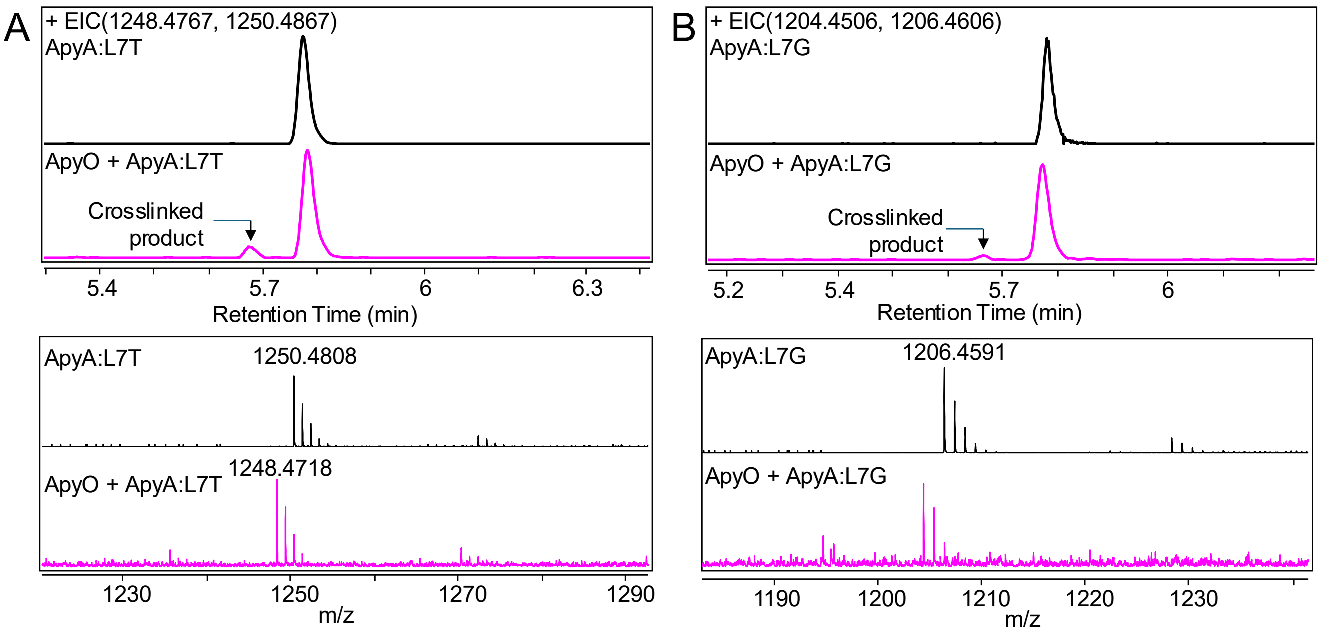

**Figure S12. LC-HRMS analysis of Leu7 variants of ApyA$_{ct}$.** Extracted ion chromatogram (EIC; top) of the in vitro reaction of ApyA variants where Leu7 was replaced with (A) Thr, and (B) Gly in the absence (black chromatogram) or presence (pink chromatogram) of ApyO. The corresponding HRMS isotopic distribution in the absence (black spectrum) or presence (pink spectrum) of ApyO is displayed for the $[M+H]^+$ ion. These residues were modified by ApyO (in addition to the ones shown in main text Figure 5) as evident from the 2 Da loss in mass.

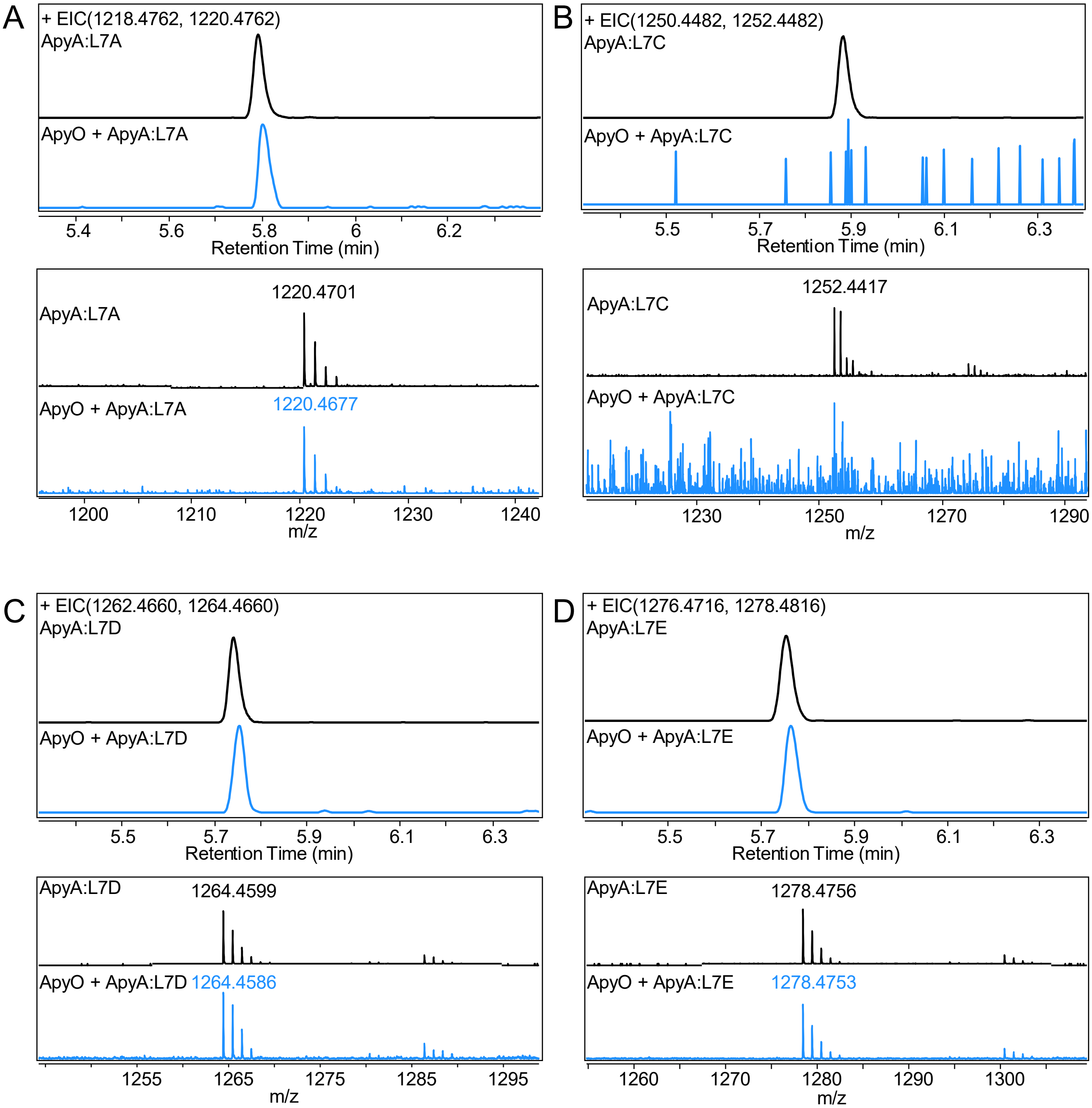

**Figure S13. LC-HRMS analysis of Leu7 variants of ApyA$_{ct}$.** Extracted ion chromatogram (EIC; top) of the in vitro reaction of ApyA variants where Leu7 was replaced with (A) Ala, (B) Cys, (C) Asp and (D) Glu in the absence (black chromatogram) or presence (blue chromatogram) of ApyO. The corresponding HRMS isotopic distribution in the absence (black spectrum) or presence (blue spectrum) of ApyO is displayed for the $[M+H]^+$ ion. For the L7C mutant, neither the substrate nor the product was observed after treatment with ApyO.

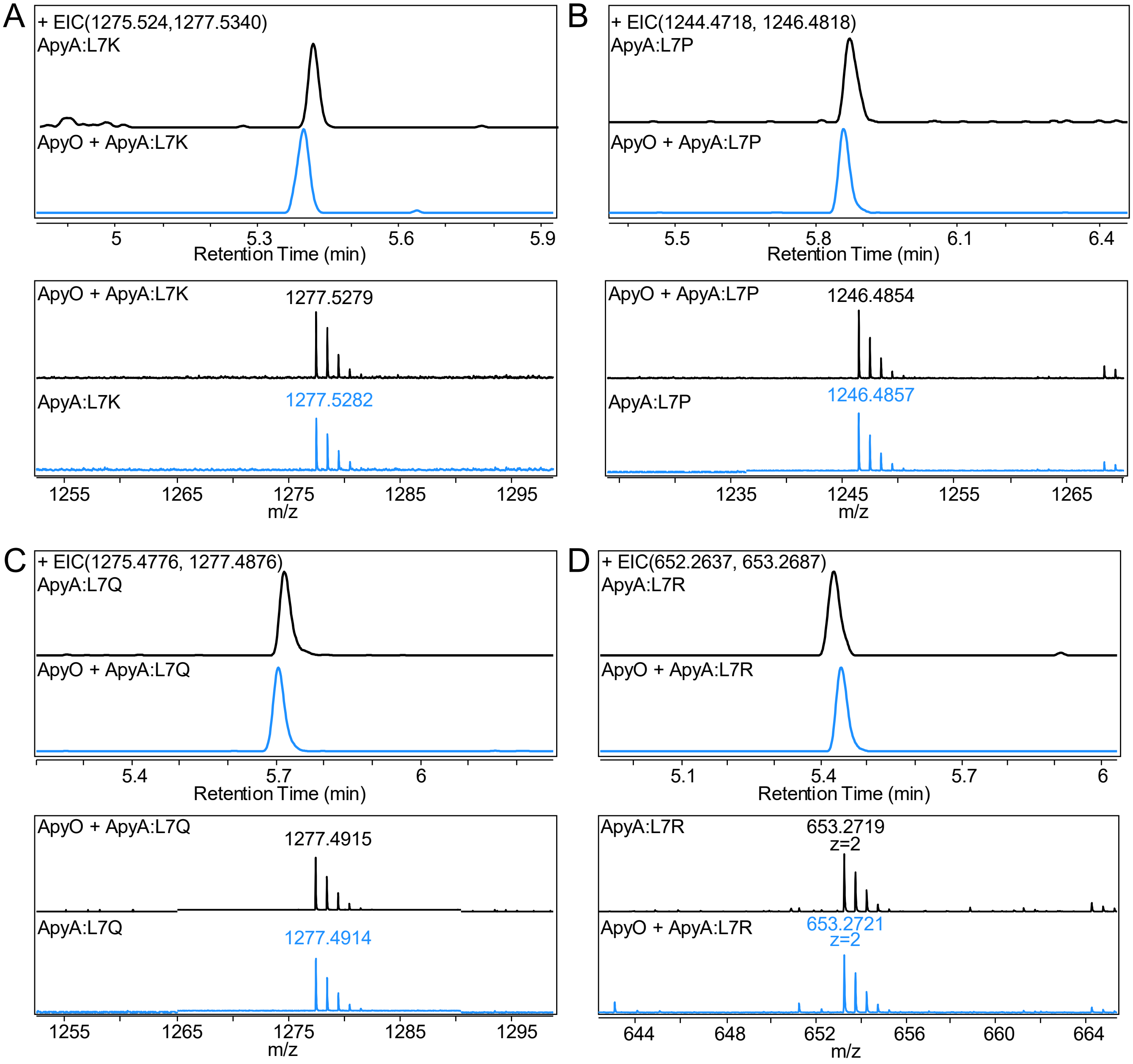

**Figure S14. LC-HRMS analysis of Leu7 variants of ApyA$_{ct}$.** Extracted ion chromatogram (EIC; top) of the in vitro reaction of ApyA variants where Leu7 was replaced with (A) Lys, (B) Pro, (C) Gln and (D) Arg in the absence (black chromatogram) or presence (blue chromatogram) of ApyO. The corresponding HRMS isotopic distribution in the absence (black spectrum) or presence (blue spectrum) of ApyO is displayed for the $[M+H]^+$ ion except for the L7R mutant where the $[M+2H]^{2+}$ ion was observed.

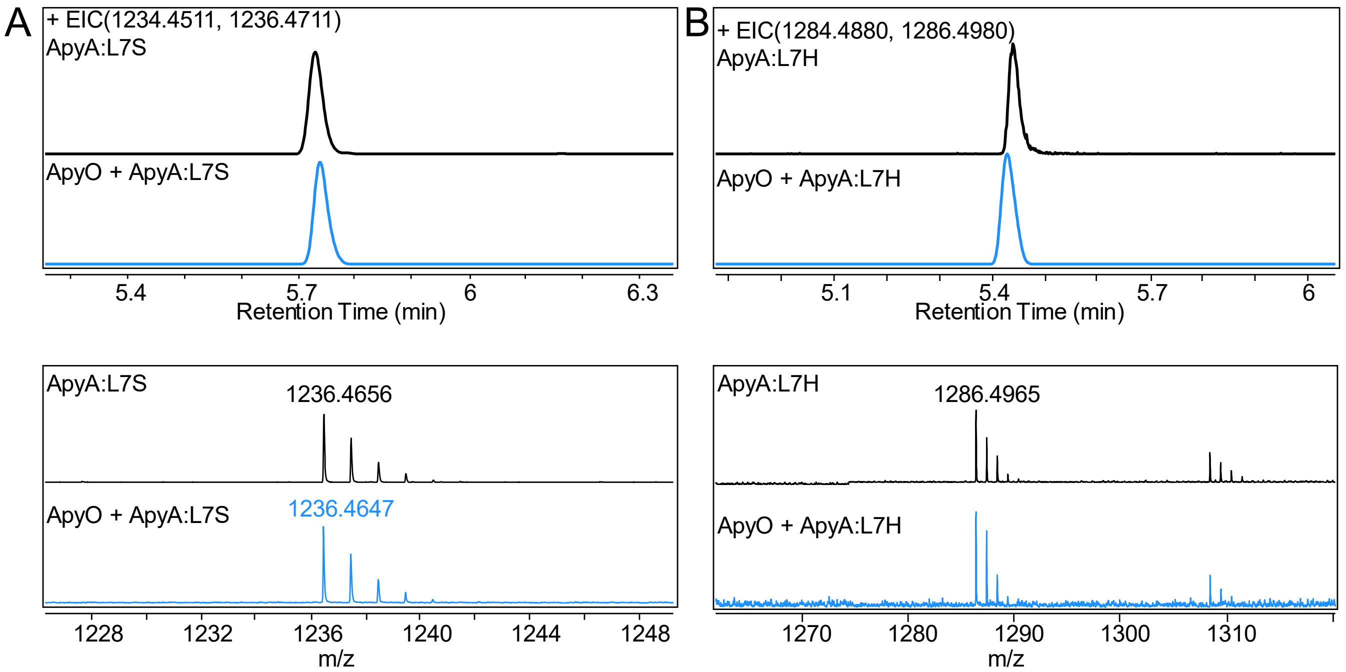

**Figure S15. LC-HRMS analysis of Leu7 variants of ApyA$_{ct}$.** Extracted ion chromatogram (EIC; top) of the in vitro reaction of ApyA variants where Leu7 was replaced with (A) Ser, and (B) His in the absence (black chromatogram) or presence (blue chromatogram) of ApyO. The corresponding HRMS isotopic distribution in the absence (black spectrum) or presence (blue spectrum) of ApyO is displayed for the $[M+H]^+$ ion.

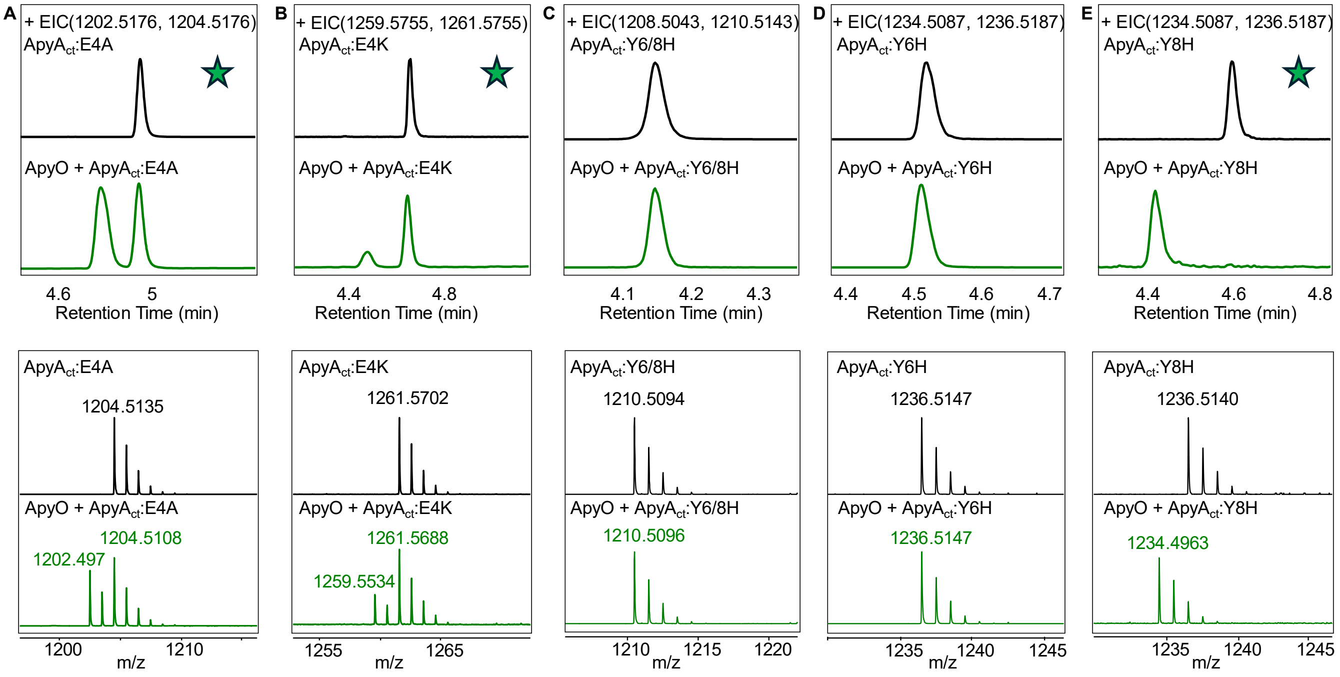

**Figure S16. LC-HRMS analysis of variants of ApyA$_{ct}$.** Extracted ion chromatograms (EIC; top) of the in vitro reaction of ApyA$_{ct}$ variants with ApyO. (A) ApyA$_{ct}$-E4A, (B) ApyA$_{ct}$-E4K (C) ApyA$_{ct}$-Y6H/Y8H, (D) ApyA$_{ct}$-Y6H, and (E) ApyA$_{ct}$-Y8H in the absence (black chromatogram) or presence (green chromatogram) of ApyO. The corresponding HRMS isotopic distribution (bottom) in the absence (black spectrum) or presence (green spectrum) of ApyO is displayed for the $[M+H]^+$ ion. Star indicates the variants that underwent some level of modification.

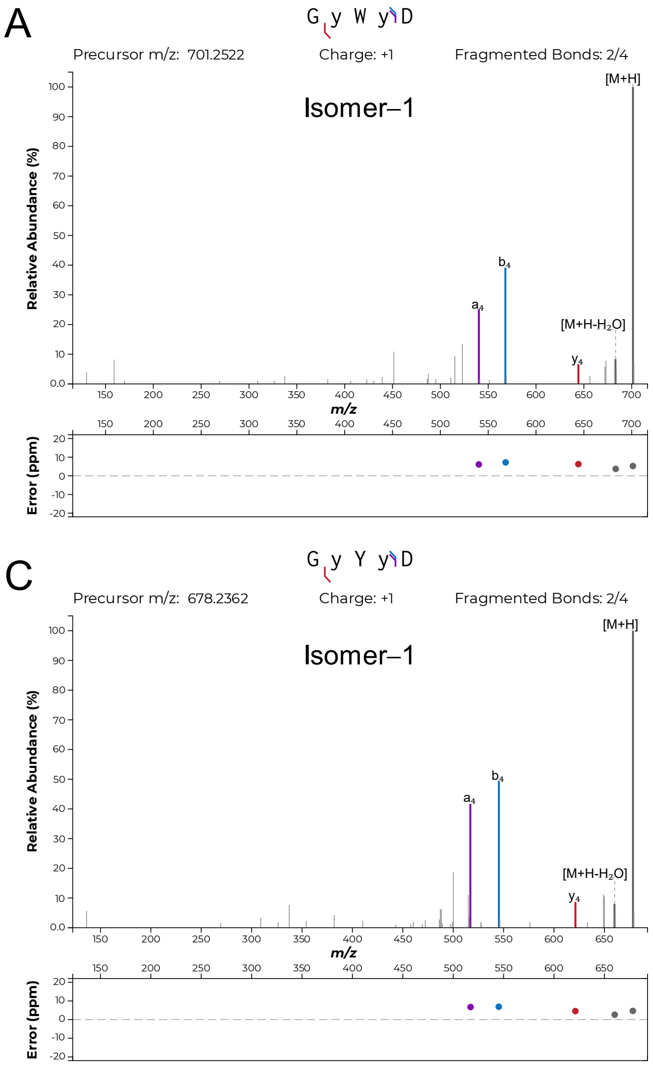

**Figure S17. Tandem MS analysis of ApyA-L7W and -L7Y isomers.** MS/MS fragmentation of (A) ApyA-L7W Isomer–1, (B) ApyA-L7W Isomer–2, (C) ApyA-L7Y Isomer–1, and (D) ApyA-L7Y Isomer–2. No b and y ion fragments were observed between the aromatic residue-containing tripeptide motifs. For residue numbering, see Fig. 2. Fragment ion annotation was performed using the interactive peptide spectral annotator[1] with residues indicated in lower case y entered as dehydrogenated through a crosslink (M - 2 Da).

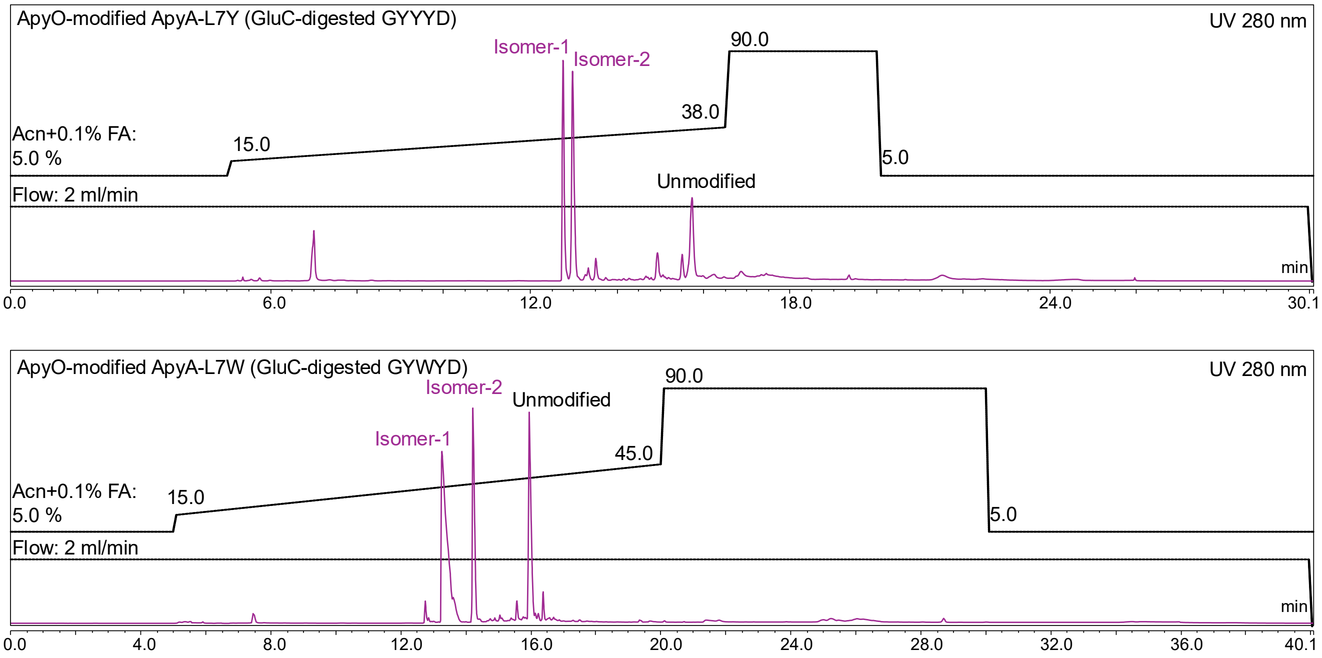

**Figure S18. HPLC purification of ApyO-modified and GluC-digested ApyA-L7Y and -L7W isomers obtained from co-expression in *E. coli*.**

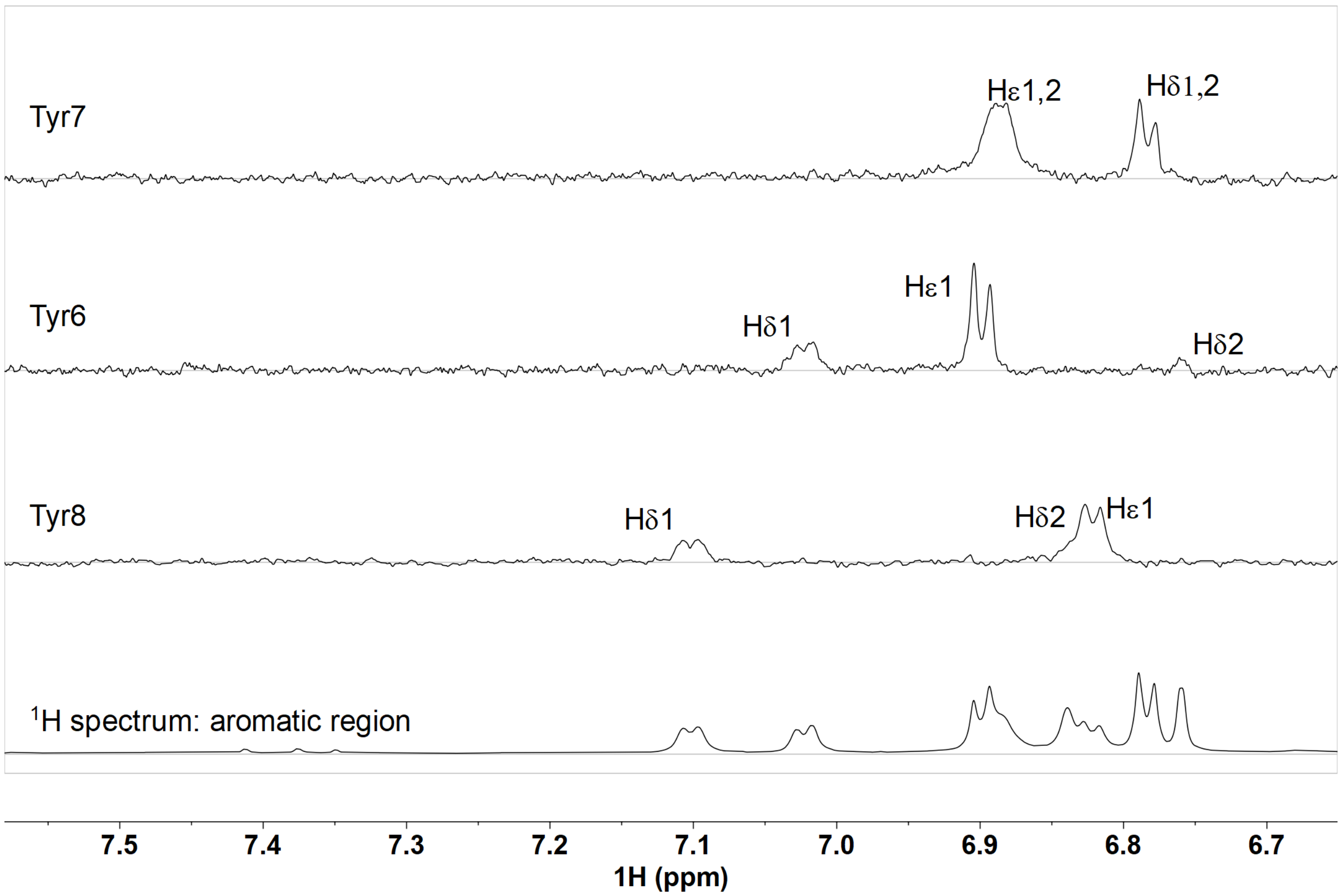

**Figure S19. 1D $^{1}H$-$^{1}H$ TOCSY showing the spin system of the aromatic region of residues 6, 7 and 8 of ApyA-L7Y isomer-1 in 90% $H_2O$, 10% $D_2O$, and 0.2% deuterated formic acid (dFA), collected at 50 ºC.** Three protons were observed for Tyr6 and Tyr8 on their aromatic rings. For Tyr7, the symmetrical two pairs of protons were observed as expected for an unmodified Tyr residue.

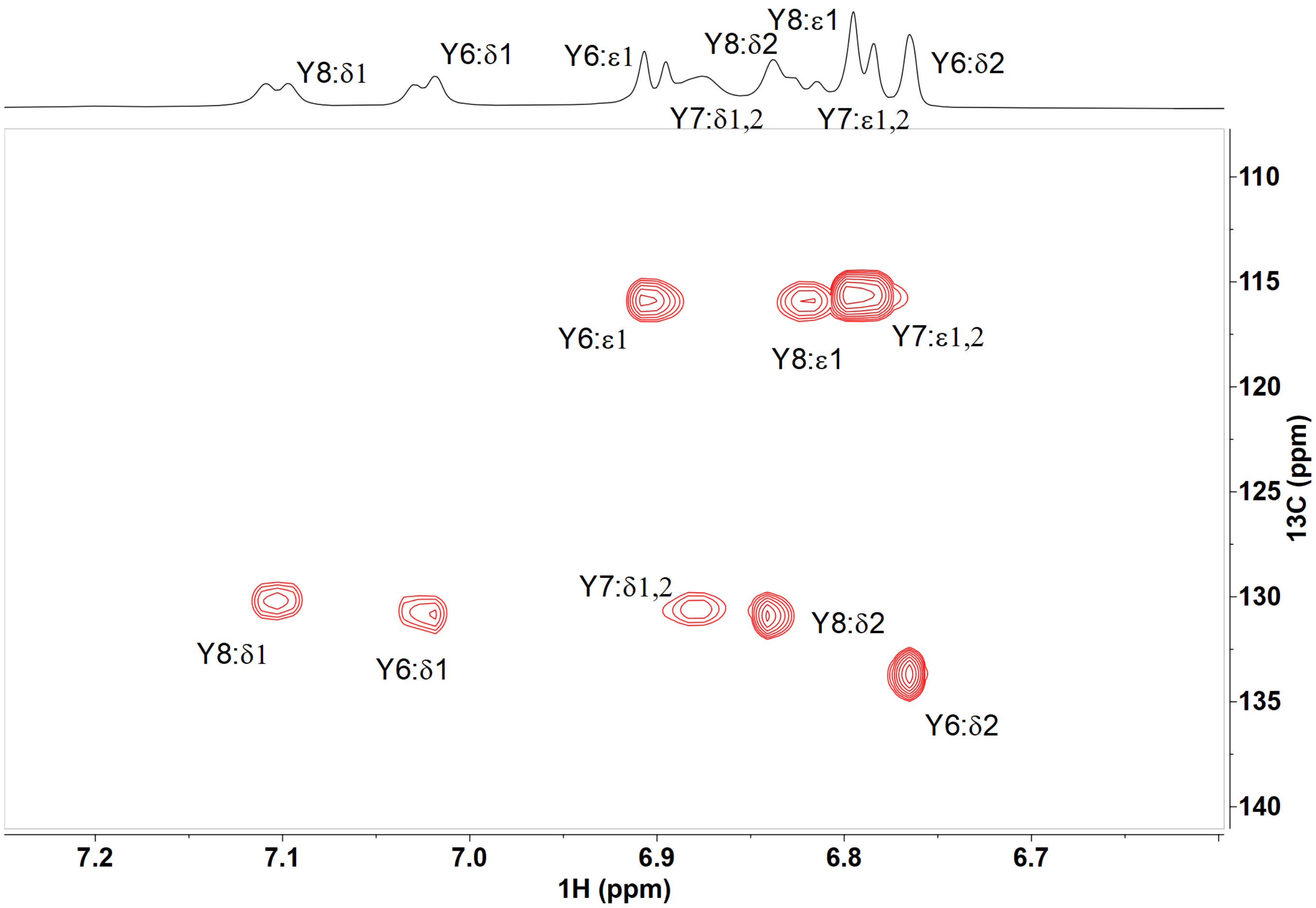

**Figure S20. 2D $^{1}H$-$^{13}C$ HSQC showing the aromatic region of ApyA-L7Y isomer-1 in 90% $H_2O$, 10% $D_2O$, and 0.2% dFA, collected at 45 ºC.** Three aromatic proton-carbon cross peaks were observed for both Tyr6 and Tyr8. For Tyr7, the symmetrical two pairs of protons were observed as expected for an unmodified Tyr residue.

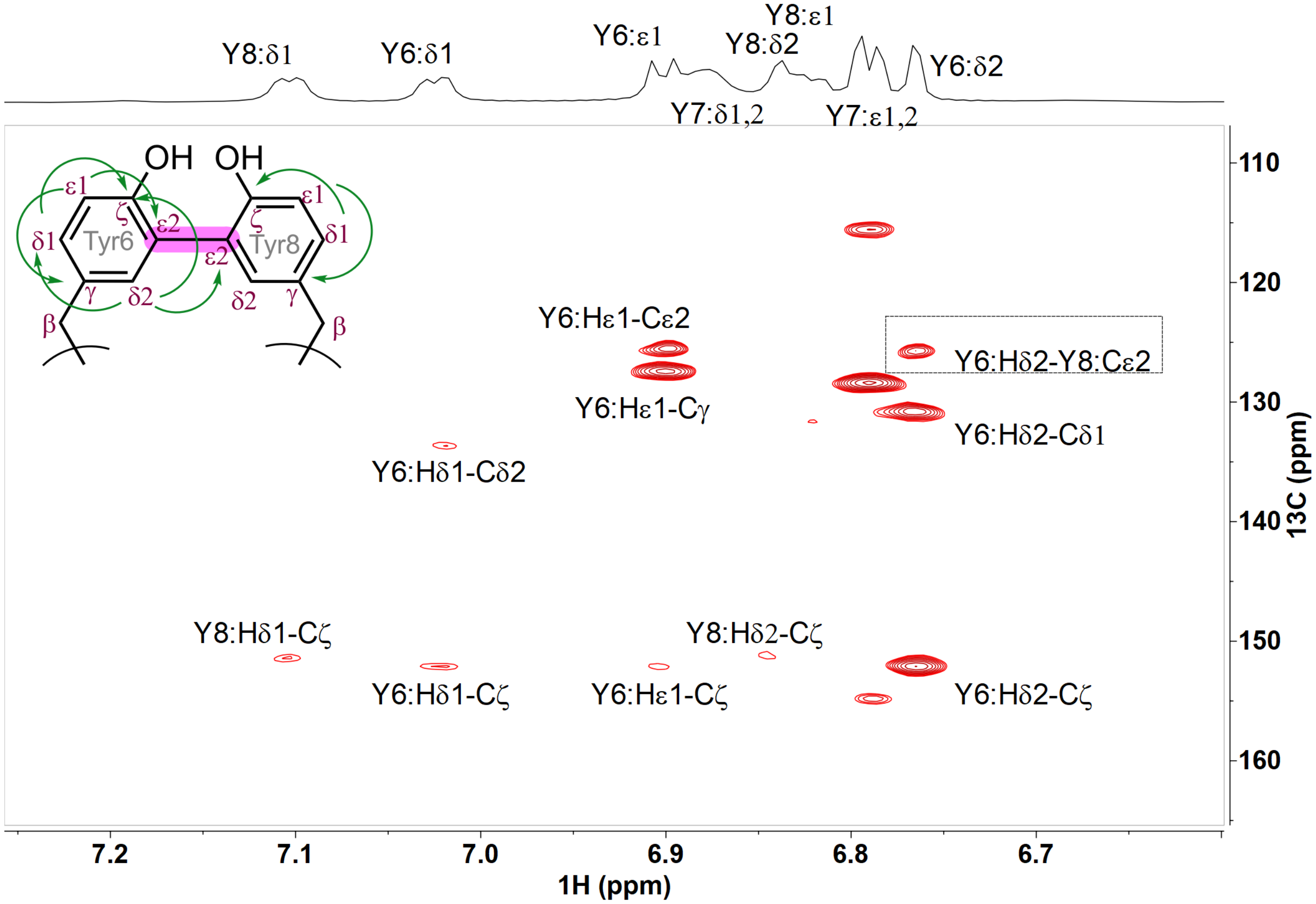

**Figure S21. The aromatic region of 2D $^{1}H$-$^{13}C$ HMBC of ApyA-L7Y isomer-1 in 90% $H_2O$, 10% $D_2O$, and 0.2% dFA, collected at 45 ºC.** Critical cross peaks involving the Tyr6 and Tyr8 residues, highlighting the ring patterns and the linkage between the two Tyr rings via a C-C bond, are shown.

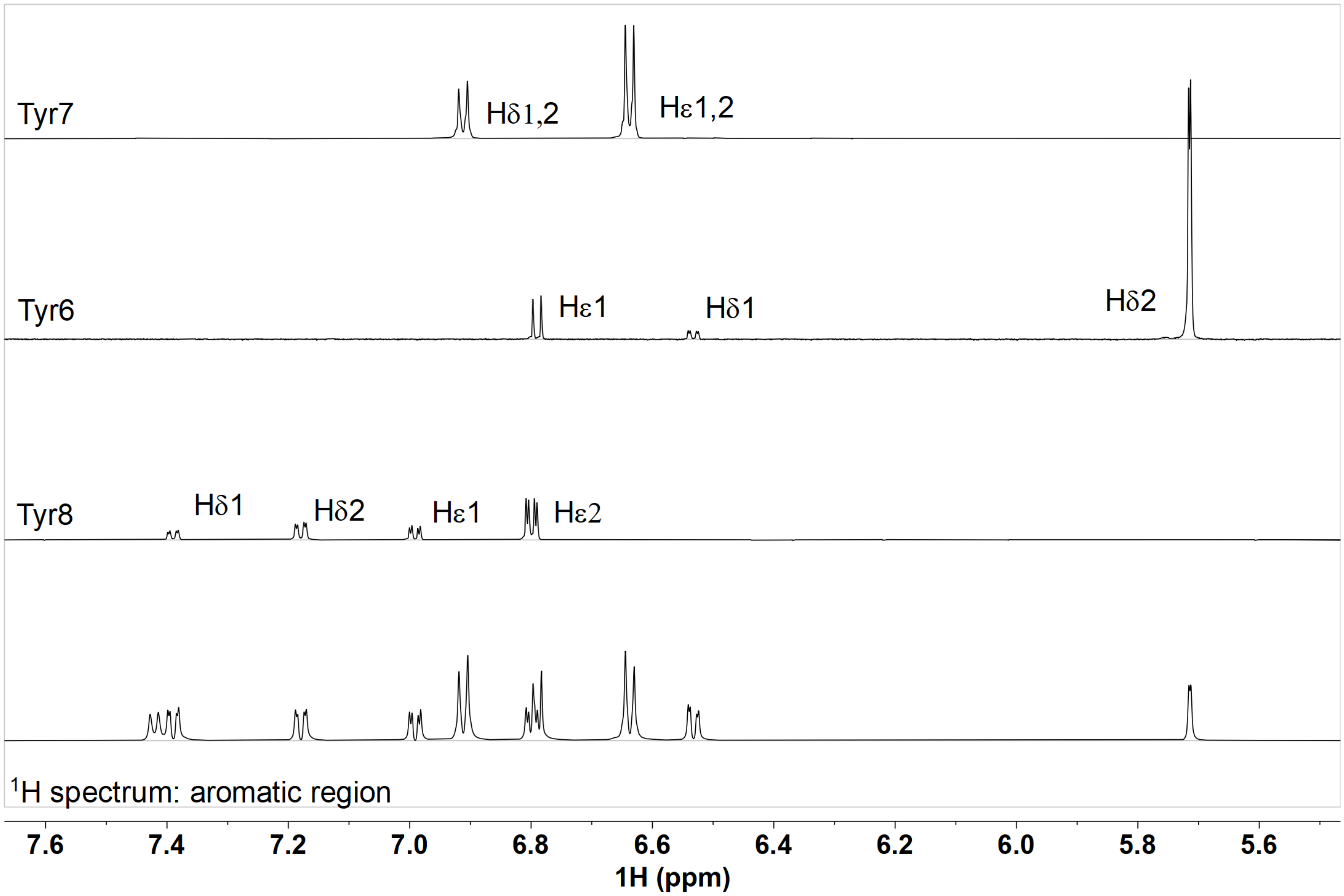

**Figure S22. 1D $^1H$-$^1H$ TOCSY data showing the spin system of the aromatic region of ApyA-L7Y isomer-2 in 90% $H_2O$, 10% $D_2O$, and 0.2% dFA.** Three aromatic protons for Tyr6 and four aromatic protons for Tyr8 were observed. For Tyr7, the unmodified symmetrical 2 pairs of protons were observed as well.

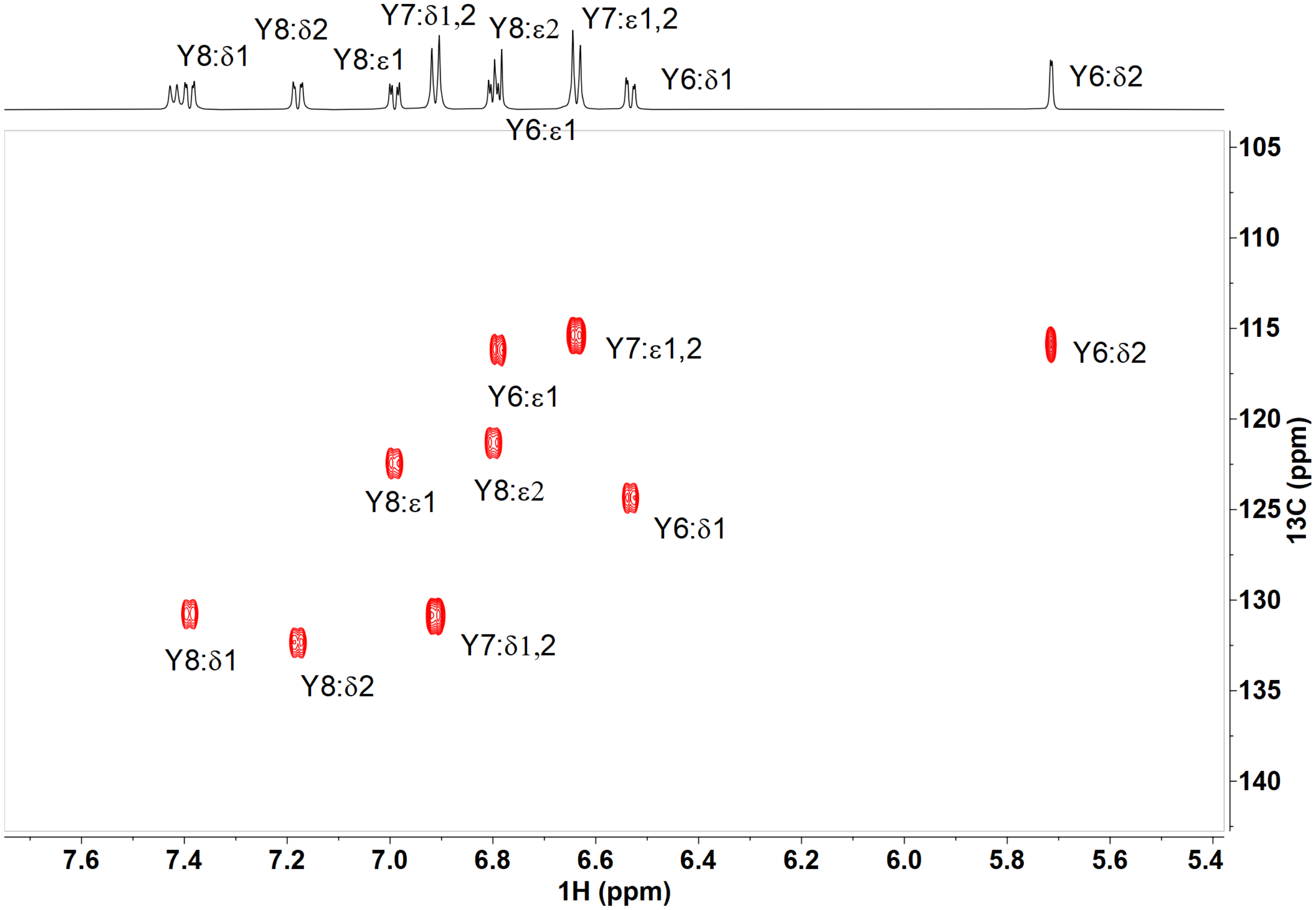

**Figure S23. 2D [1]H-[13]C HSQC showing the aromatic region of ApyO-modified ApyA-L7Y isomer-2 in 90% $H_2O$, 10% $D_2O$, and 0.2% dFA.** Three and four aromatic proton-carbon cross peaks were observed for Tyr6 and Tyr8, respectively.

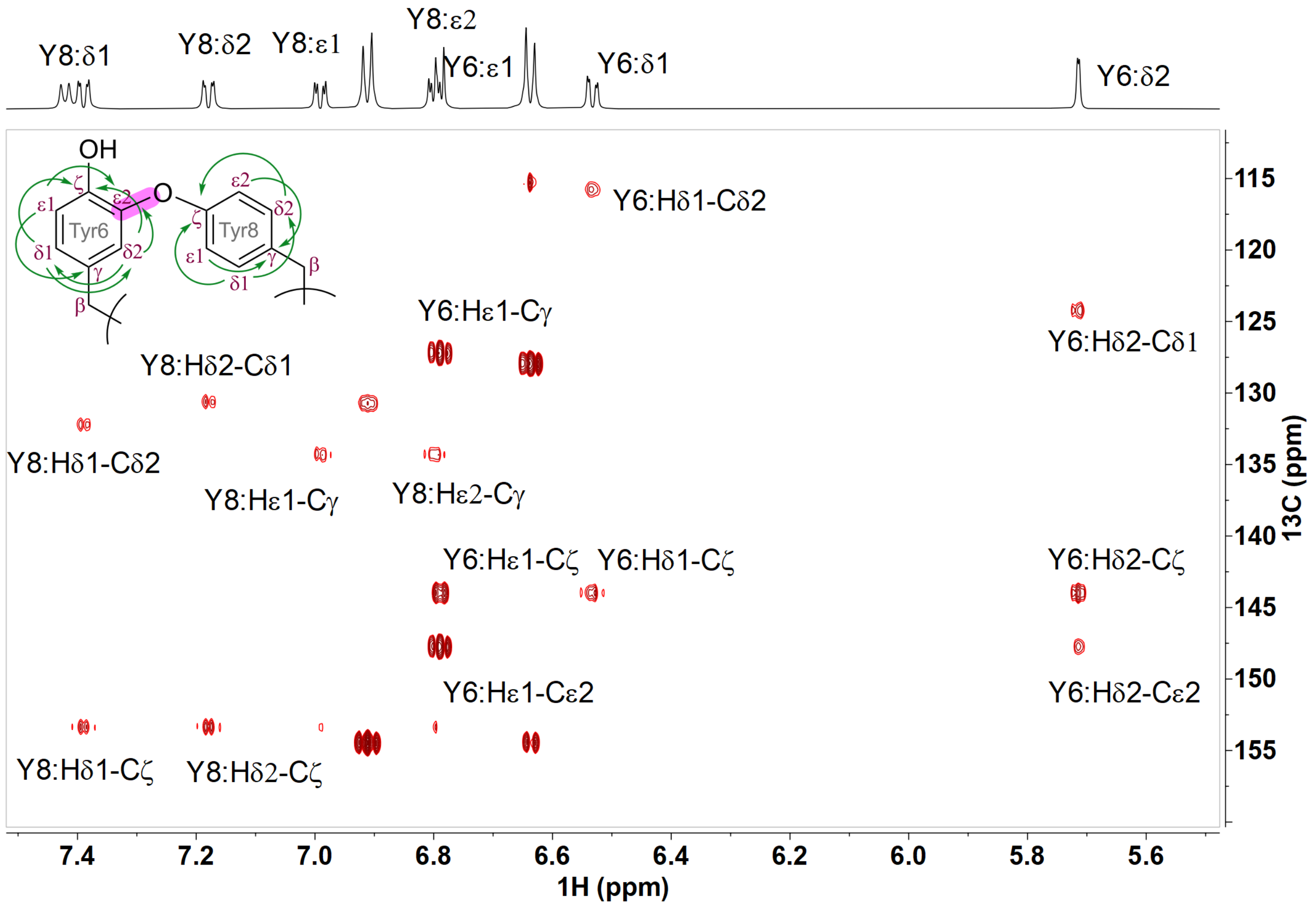

**Figure S24. 2D $^1$H-$^{13}$C HMBC data showing the aromatic region of ApyO-modified ApyA-L7Y isomer-2 in 90% $H_2O$, 10% $D_2O$, and 0.2% dFA.** Select $^1$H-$^{13}$C two and three-bond cross peaks within each aromatic side chain of Tyr6 and Tyr8 indicated in the figure confirmed the spin systems of the two aromatic rings, agreeing with a C-O linkage between the two residues.

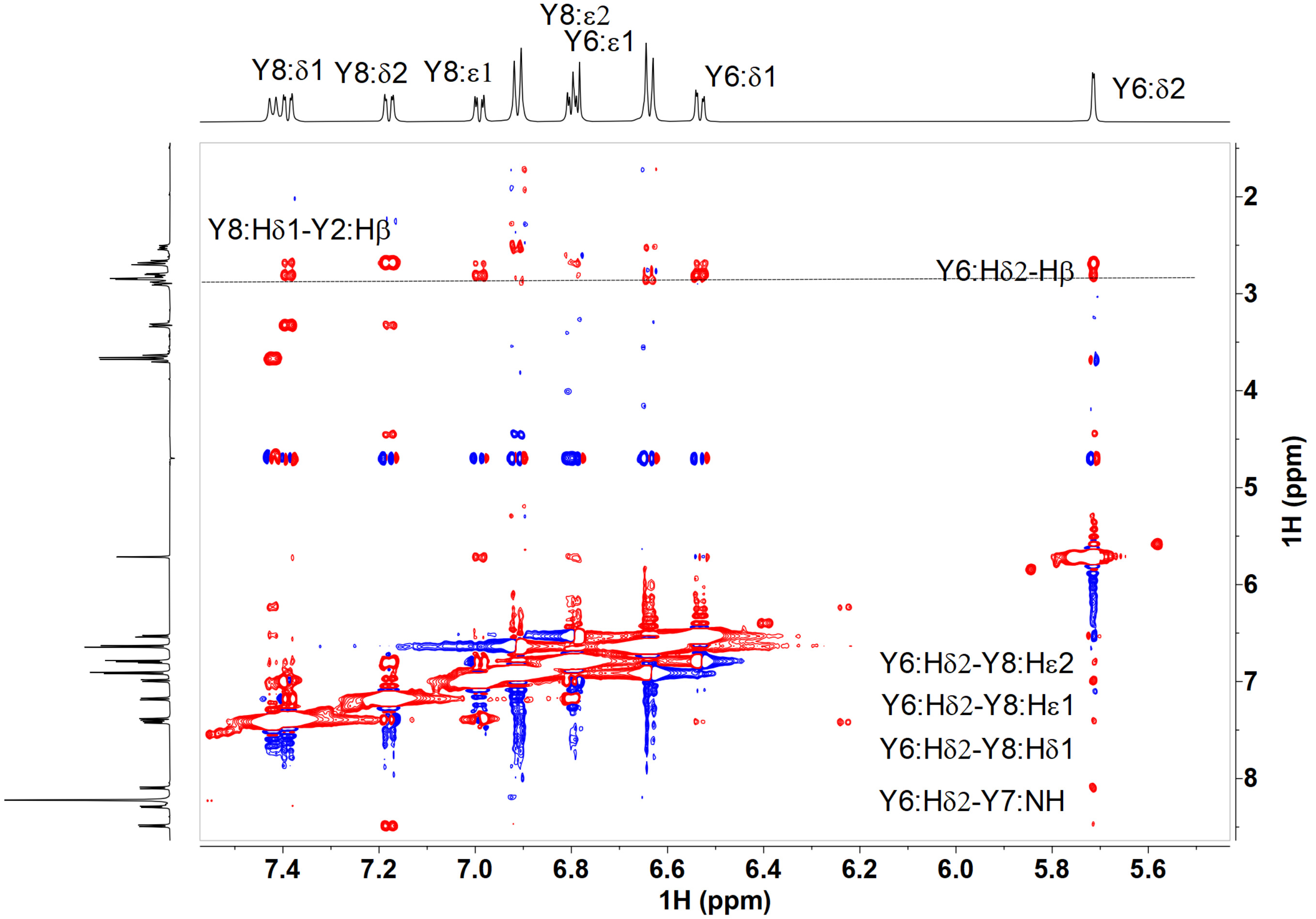

**Figure S25. 2D $^{1}$H-$^{1}$H NOESY data showing the aromatic region of ApyA-L7Y isomer-2 in 90% $H_2O$, 10% $D_2O$, and 0.2% dFA.** Select NOESY cross peaks between Hδ2 of Tyr6 and the aromatic protons of Tyr8 are consistent with the C–O linkage between the two residues.

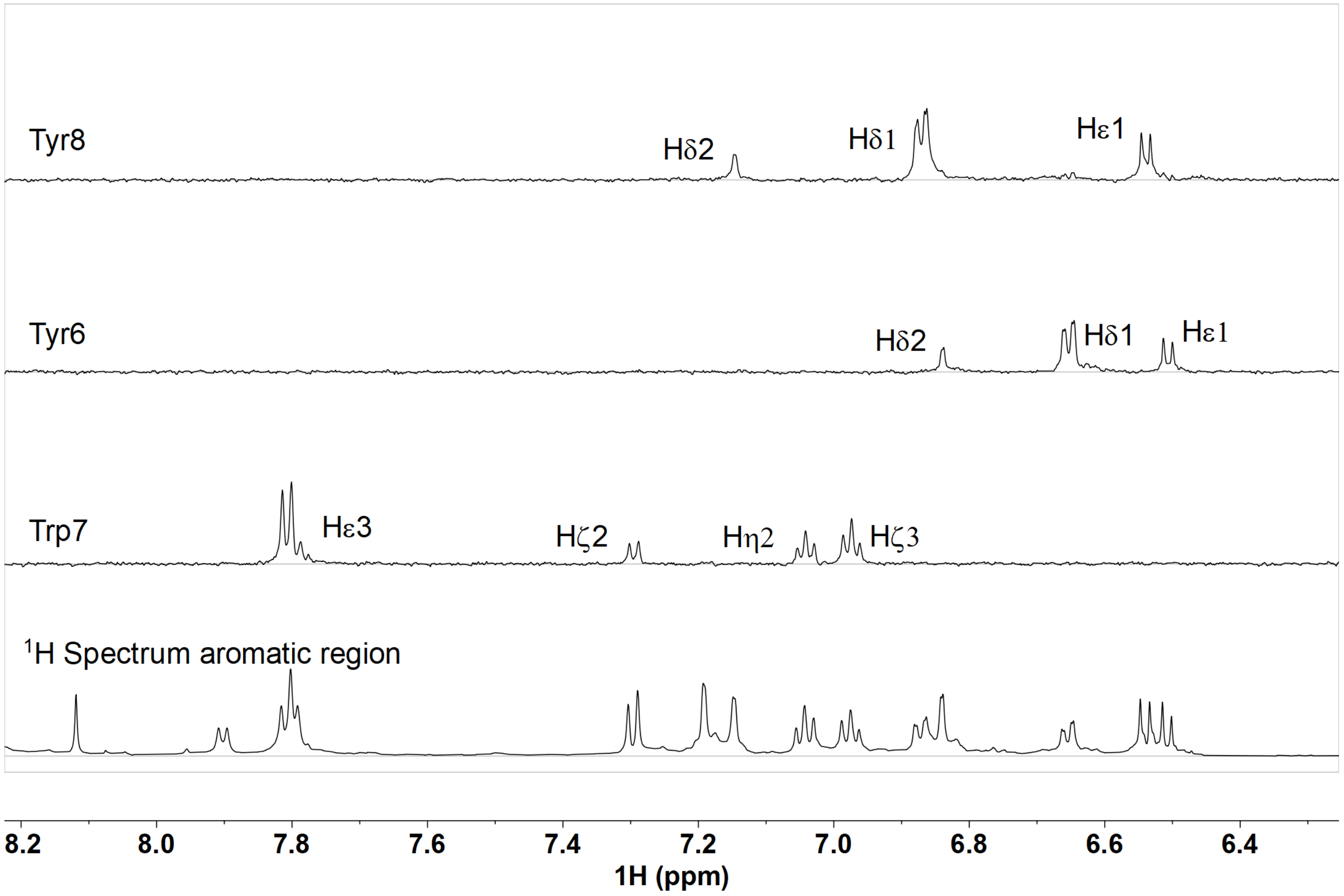

**Figure S26. 1D $^{1}H$-$^{1}H$ TOCSY spectra showing spin systems of the aromatic region of the GluC-digested, ApyO-modified ApyA-L7W isomer-1 in DMSO-$d_6$ and 0.2% dFA.** The spectra show a spin system of only three protons each in the aromatic rings of Tyr6 and Tyr8. The four aromatic protons of the phenyl ring in the indole of Trp7 are also shown.

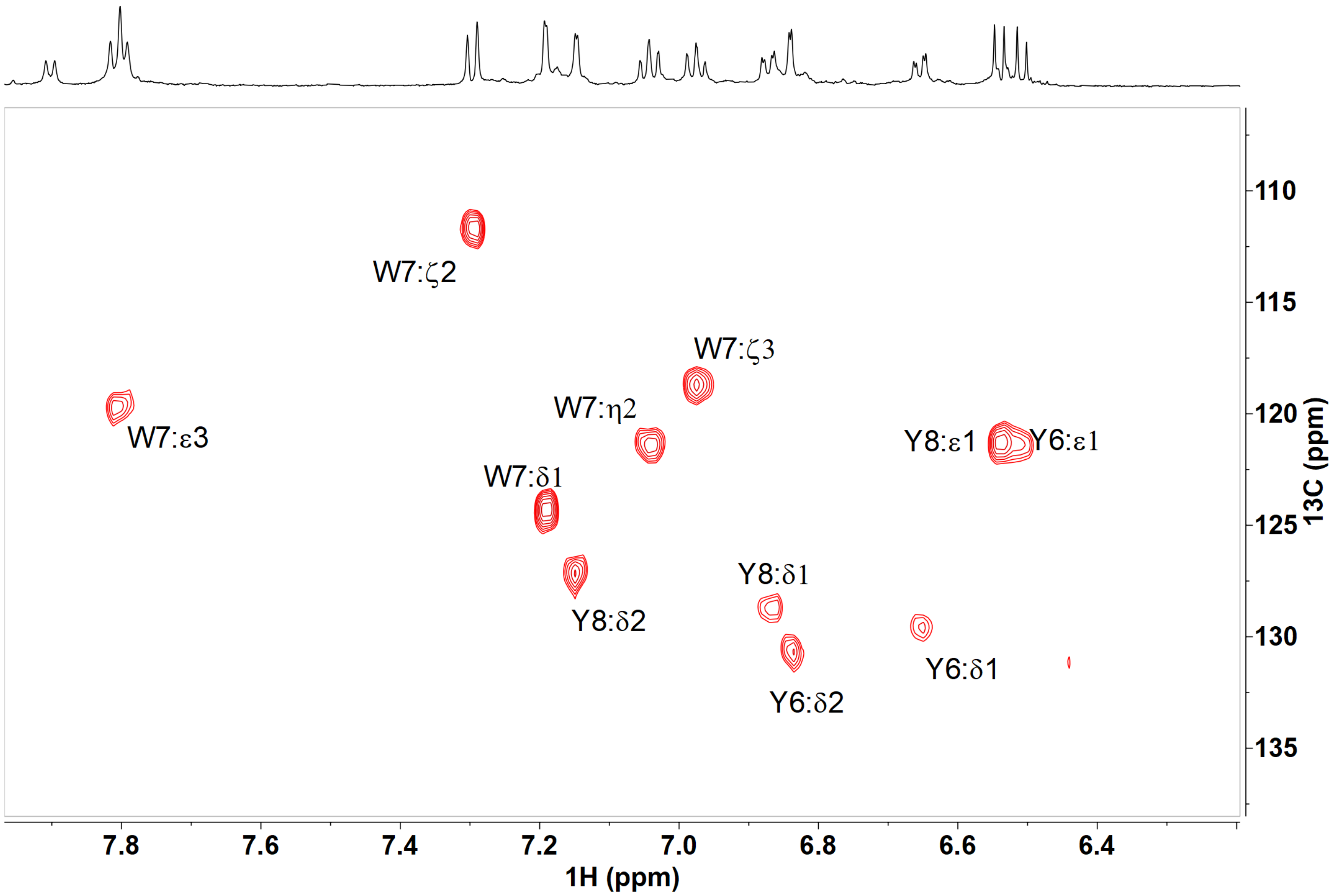

**Figure S27. The aromatic region of the 2D $^{1}H$-$^{13}C$ HSQC of GluC-digested, ApyO-modified ApyA-L7W isomer-1 in DMSO-$d_6$ and 0.2% dFA.** Three aromatic protons on each Tyr ring were observed. The assignment of the Trp7 aromatic side chain protons is also shown.

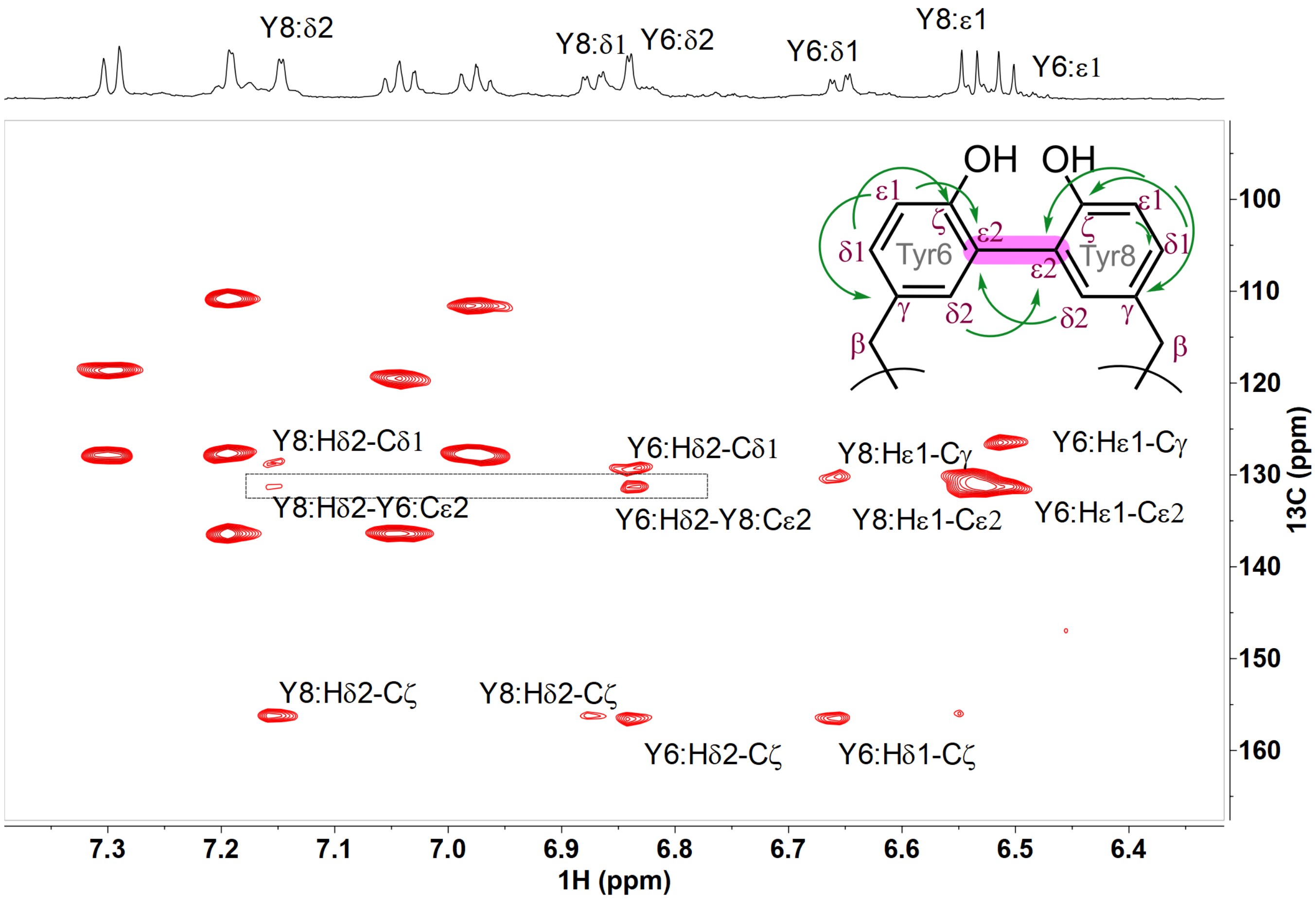

**Figure S28. The aromatic region of 2D $^{1}H$-$^{13}C$ HMBC of the GluC-digested, ApyO-modified ApyA-L7W isomer-1 in DMSO-$d_6$ and 0.2% dFA.** Critical cross peaks involving the Tyr6 and Tyr8 residues are annotated, highlighting the ring patterns and the linkage between the two Tyr rings via a C–C bond.

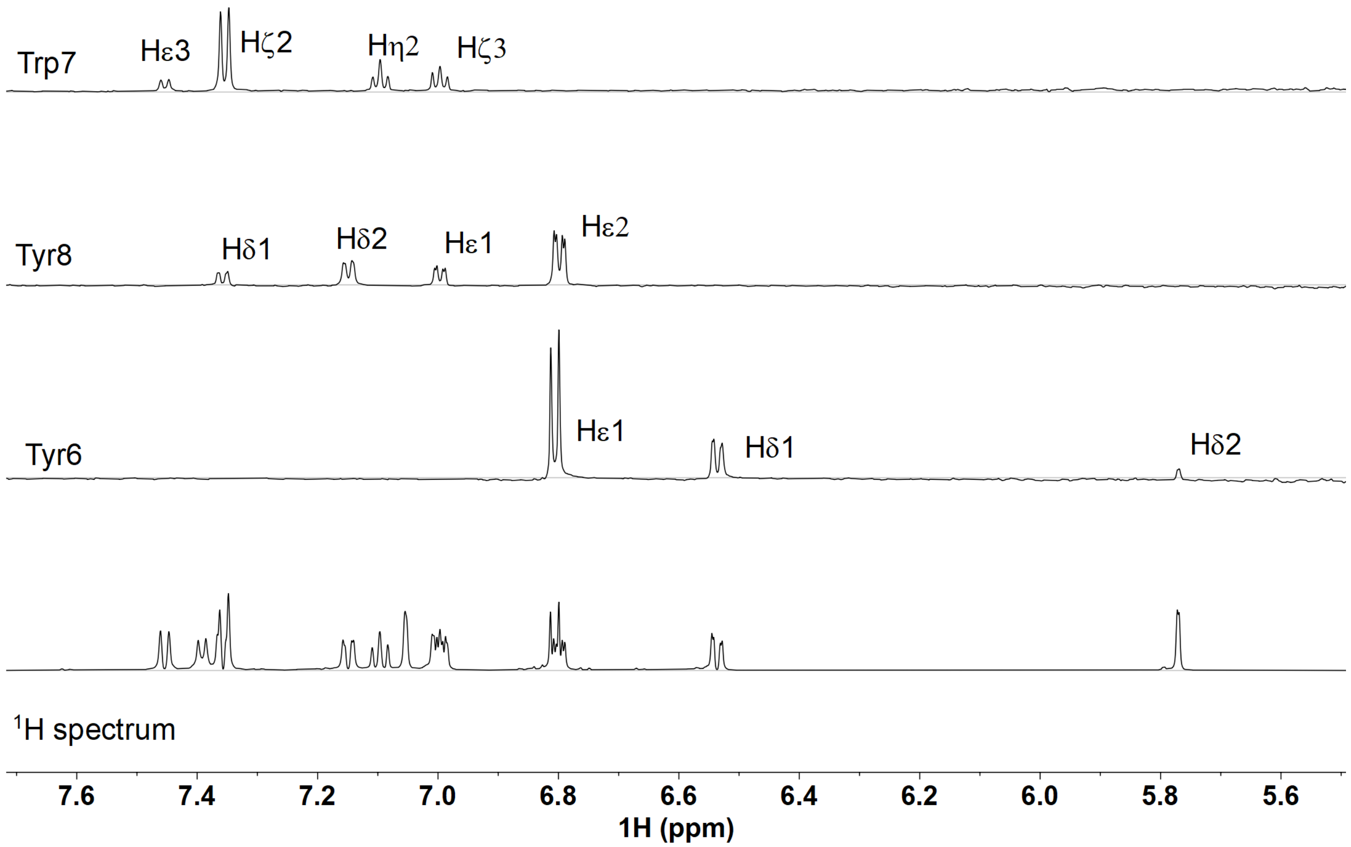

**Figure S29. 1D $^1H$-$^1H$ TOCSY spectra showing spin systems of the aromatic region of Tyr6, Trp7 and Tyr8 of the GluC-digested ApyO-modified ApyA-L7W isomer-2 in 90% $H_2O$, 10% $D_2O$, and 0.2% dFA.** Three aromatic protons of Tyr6 and four aromatic protons of Tyr8 were observed. The coupled spin system of Trp7 is also shown.

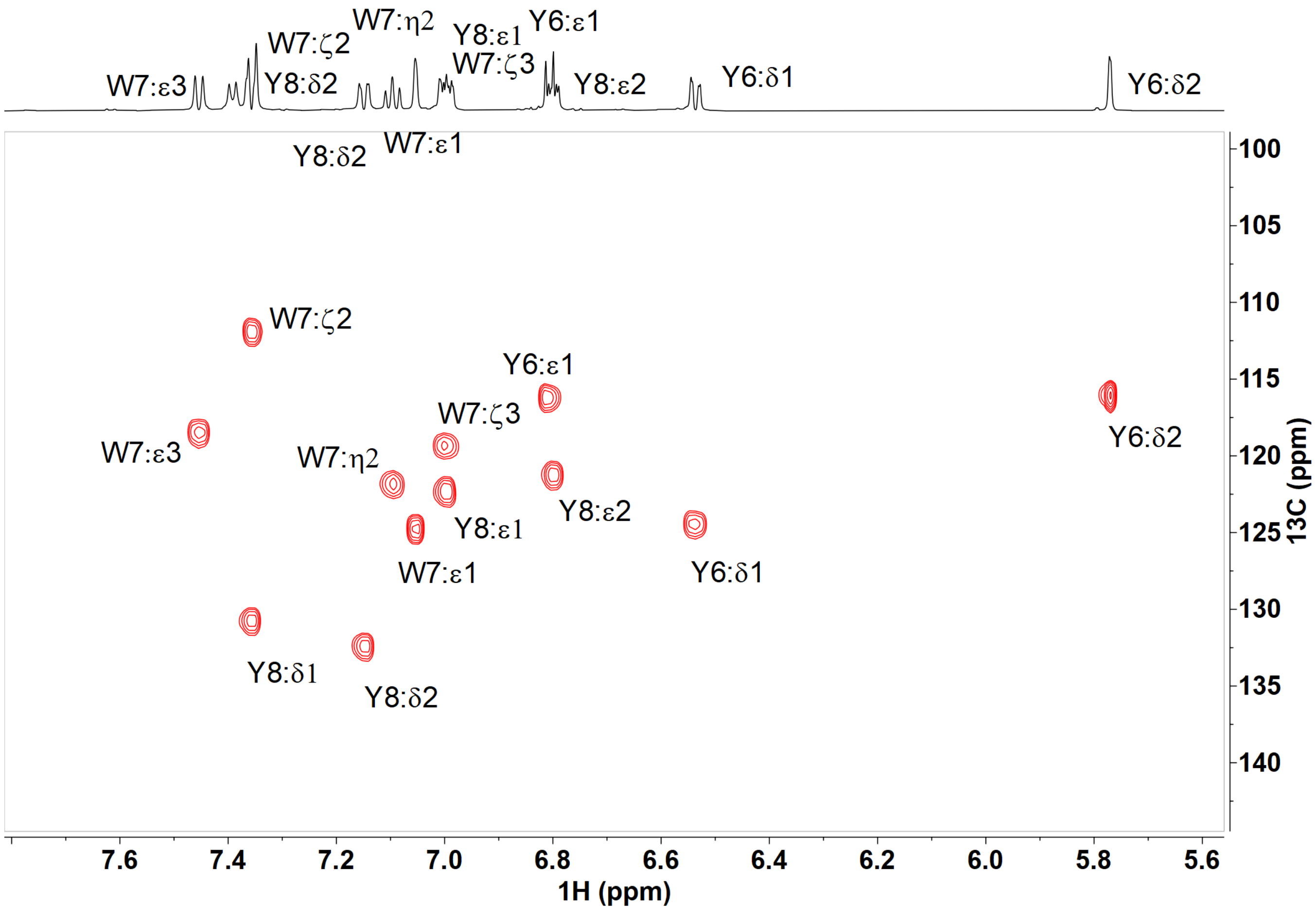

**Figure S30. 2D [1]H-[13]C HSQC data showing the aromatic region of the GluC-digested ApyO-modified ApyA-L7W isomer-2 in 90% $H_2O$, 10% $D_2O$, and 0.2% dFA.** Three and four aromatic proton-carbon cross peaks were observed for Tyr6 and Tyr8 residues, respectively.

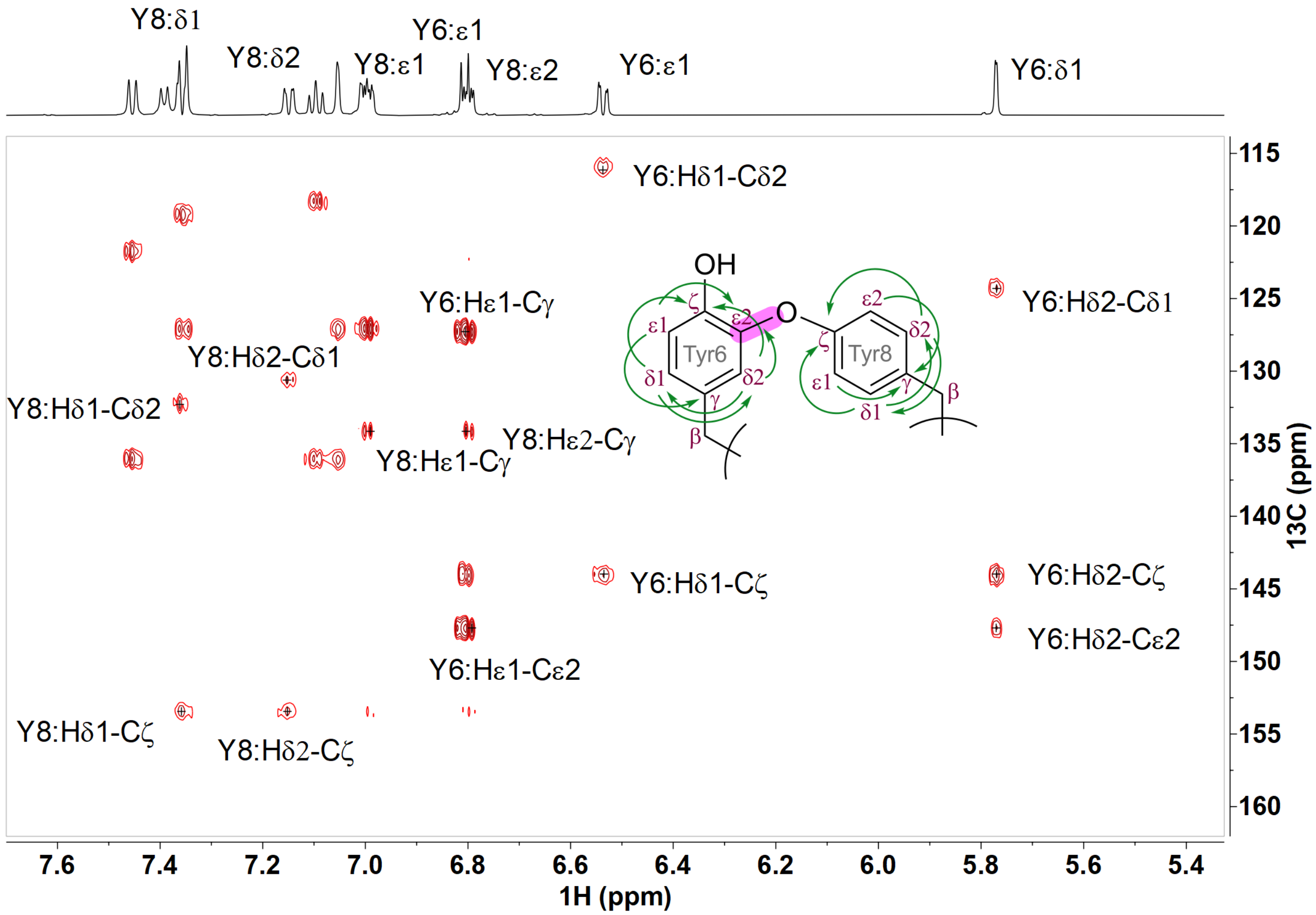

**Figure S31. 2D $^{1}H$-$^{13}C$ HMBC data showing the aromatic region of the GluC-digested ApyO-modified ApyA-L7W isomer-2 in 90% $H_2O$, 10% $D_2O$, and 0.2% dFA.** Select $^{1}H$-$^{13}C$ two and three-bond cross peaks within each aromatic side chain of Tyr6 and Tyr8 are annotated that confirmed the spin systems and the patterns of the two rings, consistent with a C–O linkage between the two residues.

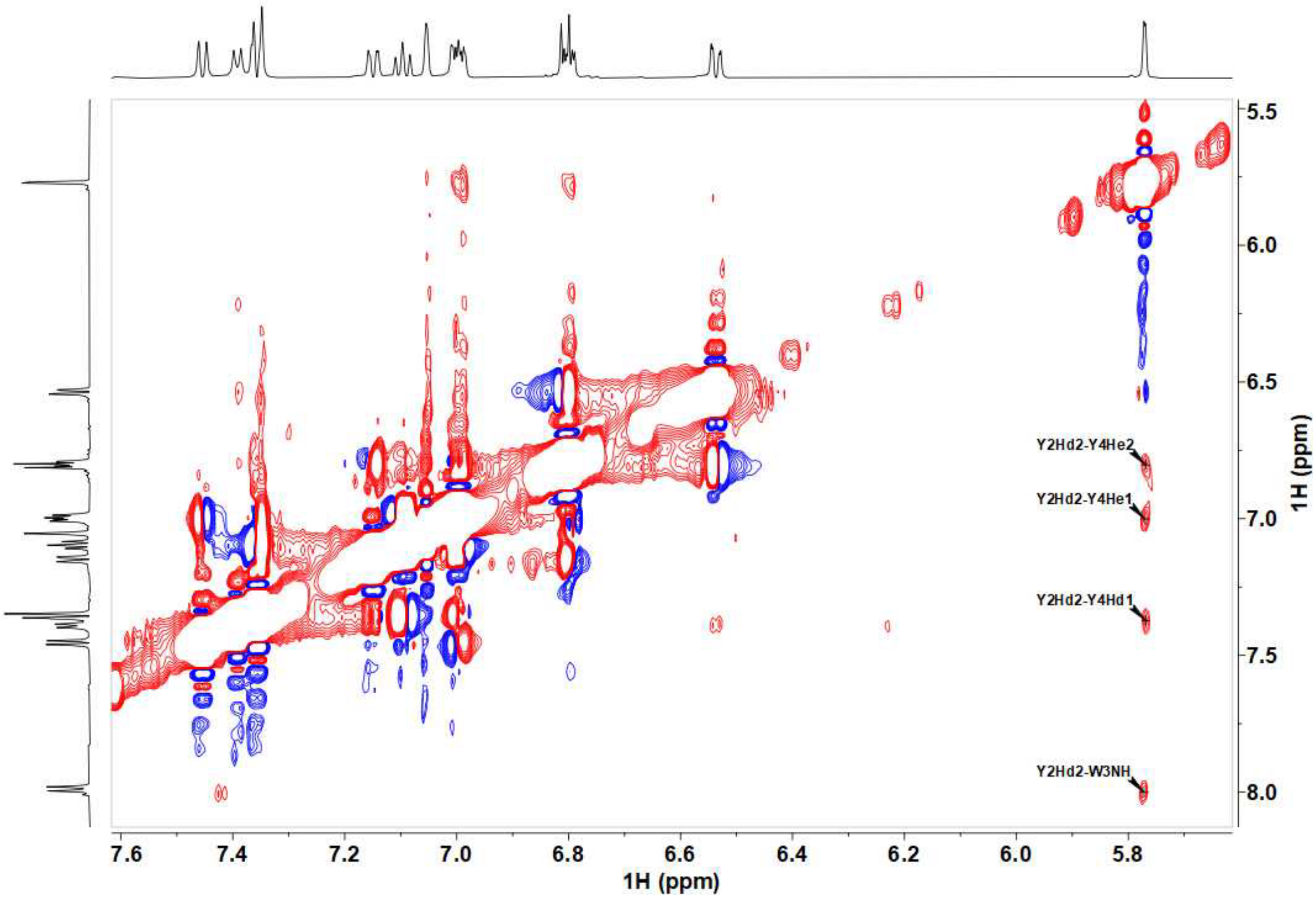

**Figure S32. 2D $^1H$-$^1H$ NOESY data showing the aromatic region of the GluC-digested ApyO-modified ApyA-L7W isomer-2 in 90% $H_2O$, 10 % $D_2O$, and 0.2% dFA.** Select NOESY cross peaks between Hδ2 proton of Tyr6 and the aromatic protons of Tyr8 shown in the figure is consistent with a C-O linkage between the two residues.

A

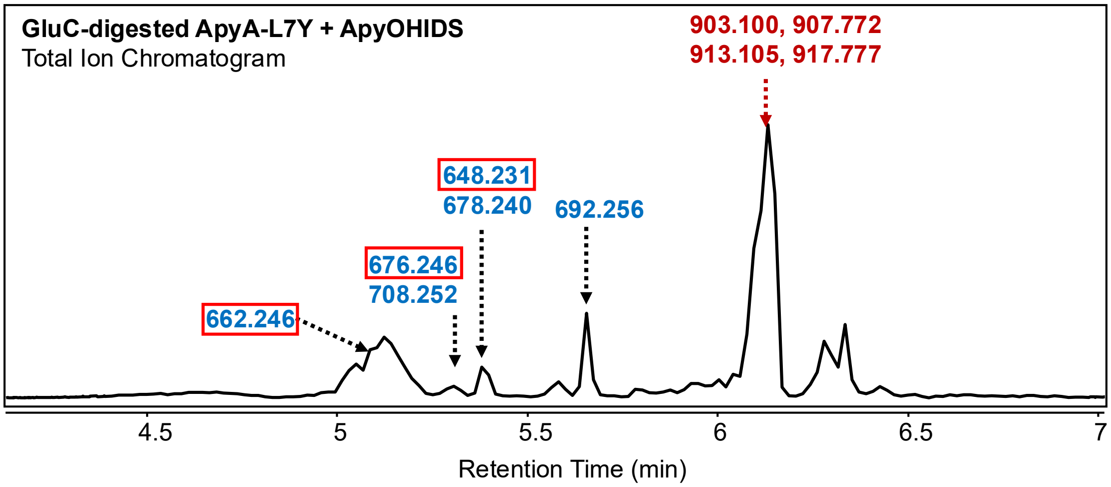

B

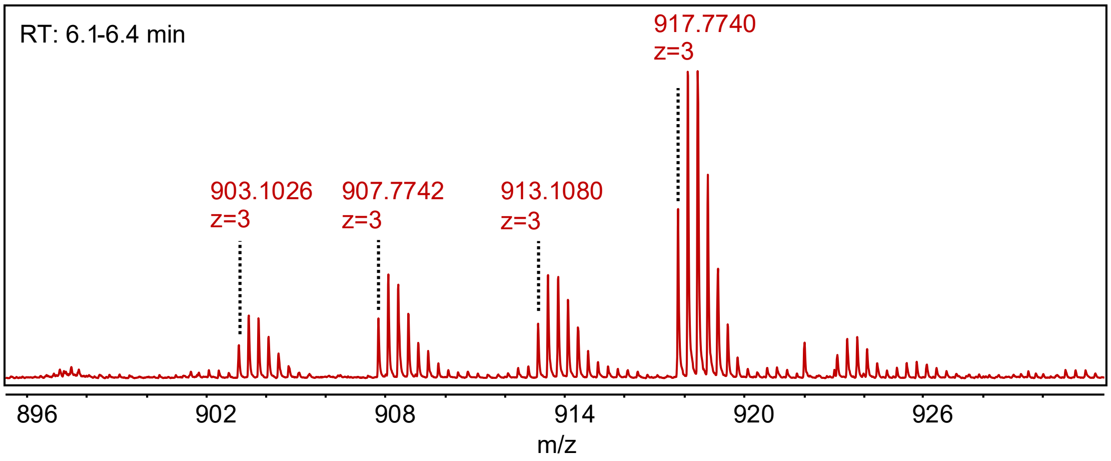

C

$[M+H]^{1+}$ for GYYYD

| | Unmodified | -2 Da | -30 Da | -32 Da |
|---|---|---|---|---|
| | 680.2562 | 678.2462 | 650.2562 | 648.2462 |
| +14 | 694.2562 | 692.2462 | 664.2562 | 662.2462 |
| +28 | 708.2562 | 706.2462 | 678.2562 | 676.2462 |

D

$[M+3H]^{3+}$ for NGKLTITMSKVDLAGGRMGEGYYYD

| | Unmodified | -2 Da | -30 Da | -32 Da |
|---|---|---|---|---|
| | 913.7772 | 913.1072 | 903.7772 | 903.1072 |
| +14 | 918.4439 | 917.7739 | 908.4439 | 907.7739 |
| +28 | 923.1106 | 922.4406 | 913.1106 | 912.4406 |

**Figure S33. LC-HRMS analysis of GluC-digested Apy-L7Y co-expressed with ApyOHIDS in *E. coli*.** (A) Total ion chromatogram of ApyO-, ApyHI-, ApyD- and ApyS-modified GluC-digested ApyA-L7Y showing a variety of products derived from the GYYYD sequence. Putative products containing the desired keto acid group are boxed. The experiment generated also incomplete GluC-digestion products (red font; $[M+3H]^{3+}$ ions). The latter were the major species and arise from GluC not cleaving at the Glu before the GYYYD sequence (red sequence in panel D), possibly due to inhibition of the enzyme by products of the pathway. (B) HRMS spectra of the partially digested products showing that the major product (917 Da) lacks the desired keto acid group. (C) Monoisotopic masses of different modifications of the pentapeptide GYYYD product shown as $[M+H]^{+}$ ions. PTMs are crosslinking (−2 Da, ApyO), oxidative decarboxylation to form the $\alpha$-keto-$\beta$-amino acid (−30 Da, ApyHI) and both (−32 Da), with or without methylation (+14) or bismethylation (+28 Da) by ApyD and ApyS. Observed masses are highlighted in yellow. (D) The partially digested products shown as $[M+3H]^{3+}$ ions. The most abundant apparent products in the blue and red series (692 and 917 Da) have been crosslinked (−2 Da) and methylated by ApyD (+14 Da).

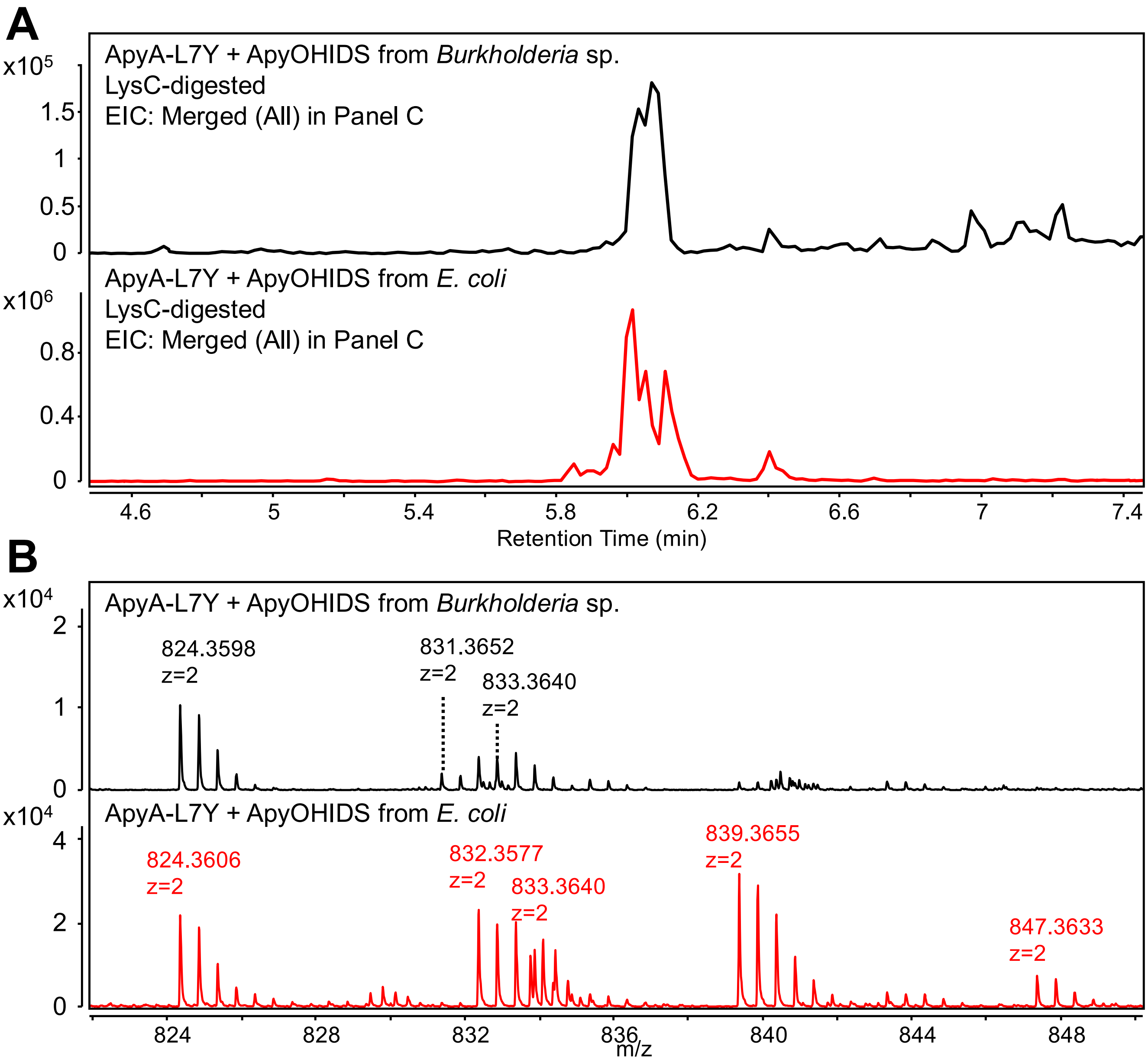

**C**

**$[M+2H]^{2+}$ for VDLAGGRMGEGYYYD**

| | Unmodified | -2 Da | -30 Da | -32 Da |
|---|---|---|---|---|
| | 833.3657 | 832.3607 | 818.3657 | 817.3607 |
| **+14** | 840.3657 | 839.3607 | 825.3657 | 824.3607 |
| **+28** | 847.3657 | 846.3607 | 832.3657 | 831.3607 |

**Figure S34. LC-HRMS analysis of LysC-digested Apy-L7Y variant co-expressed with ApyOHIDS in *E. coli* and *Burkholderia* sp.** (A) Extracted ion chromatograms of ApyO-, ApyHI-, ApyD- and ApyS-modified LysC-digested ApyA-L7Y variant produced in *Burkholderia* sp. FERM BP-3421 (black trace) and *E. coli* BL21 (DE3) TUNER (red trace). (B) The corresponding HRMS spectra shown as $[M+2H]^{2+}$ ions. (C) Monoisotopic masses of different modifications of the LysC-digested products shown as $[M+2H]^{2+}$ ions. Observed masses are highlighted in yellow. Mass shifts: Crosslink (ApyO), -2 Da; β-ketoacid (ApyHI), -30 Da; up to two methylations (ApyD and ApyS), +14 and +28 Da.

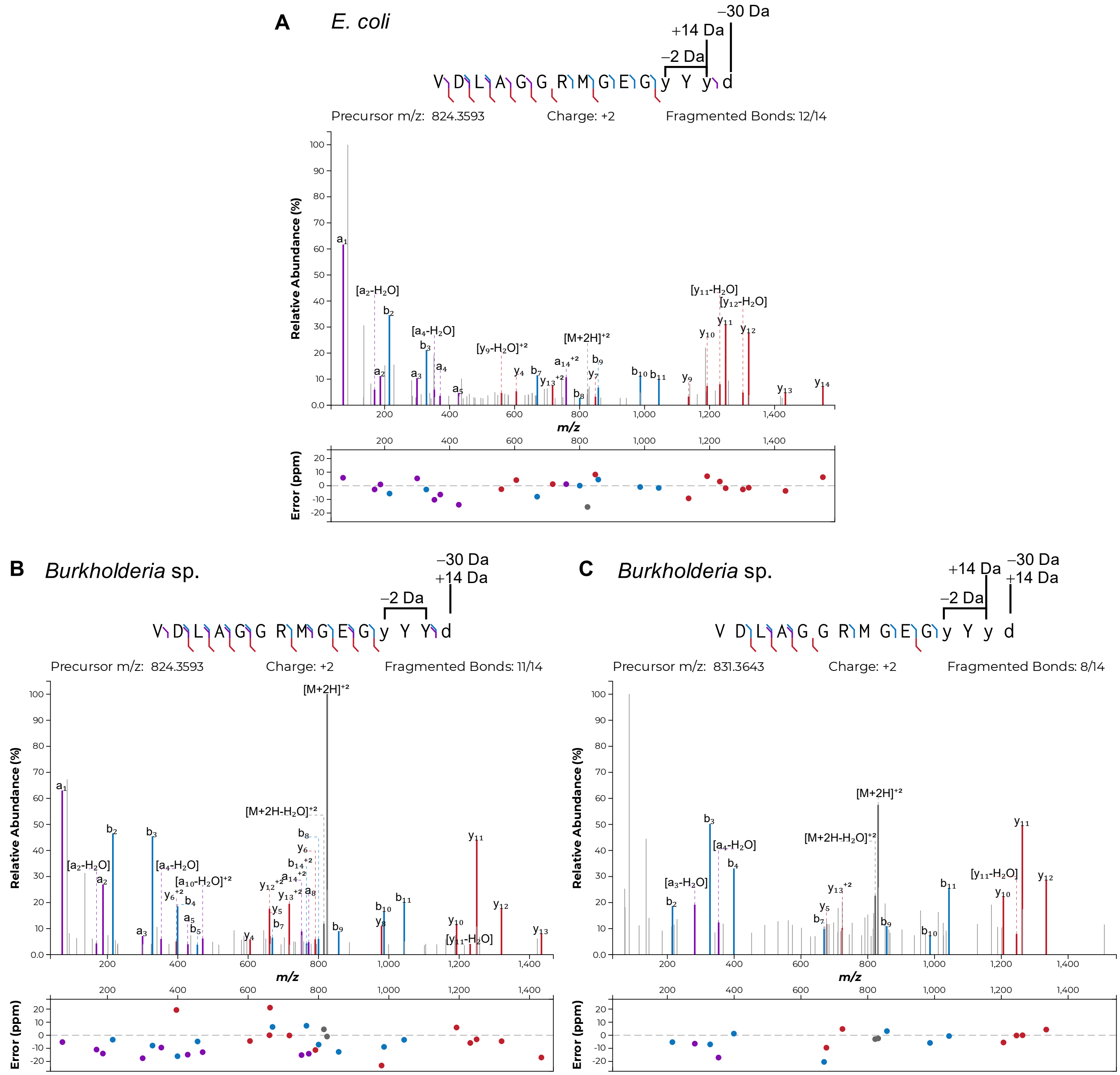

**Figure S35. Tandem MS analysis of LysC-digested Apy-L7Y variant co-expressed with ApyOHIDS.** (A) LysC-digested desmethyl-intermediate of ApyA-L7Y showing modifications installed by ApyO, ApyHI and ApyD obtained from *E. coli*. (B) LysC-digested desmethyl-intermediate of ApyA-L7Y variant showing modifications installed by ApyO, ApyHI and ApyS obtained from *Burkholderia* sp. (C) LysC-digested fully modified product obtained from *Burkholderia* sp.

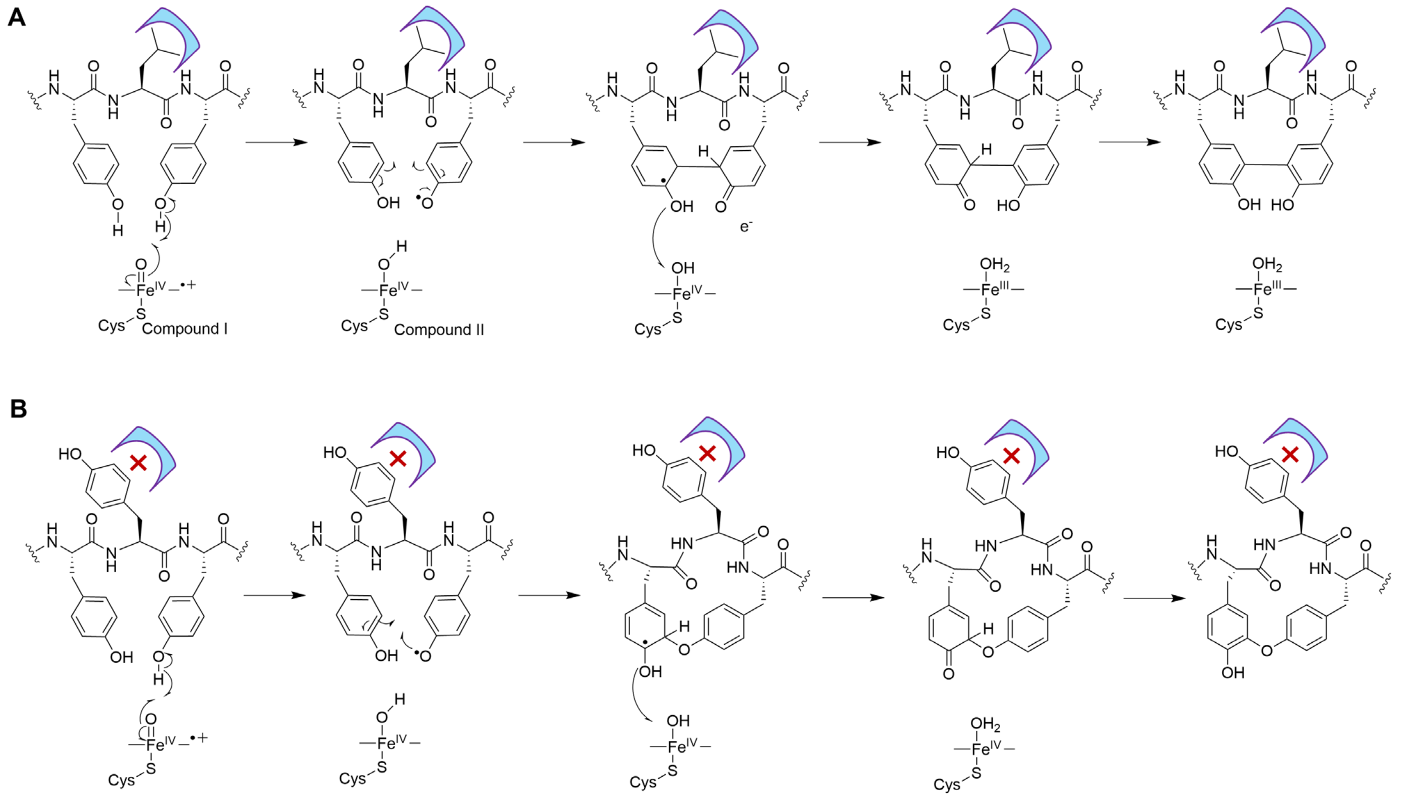

**Figure S36. Possible mechanism for the two observed constitutional isomers formed from ApyO catalysis.** (A) For the WT substrate, Compound I consisting of an Fe(IV)-oxo species and a porphyrin radical cation abstracts a hydrogen atom from one of the Tyr residues (shown for the C-terminal Tyr because it is the most conserved in Figure S1). Reaction of the Tyr radical at Cε, which carries significant spin density as a result of resonance delocalization, with the second Tyr results in the observed C-C crosslink. Electron and proton transfer to Compound II completes the catalytic cycle. (B) For the ApA-YYY mutant (L7Y), the Tyr at position 7 is proposed to perturb the orientation of Tyr6 and Tyr8 such that the initially formed Tyr radical reacts with Tyr6 through the oxygen to form a C-O crosslink. Without co-crystal structures, the subtle differences in orientation are not readily rationalized. Another possibility is that in WT vs Leu7 mutants, the initial hydrogen atom abstraction may occur from a different Tyr resulting in different catalytic outcomes.

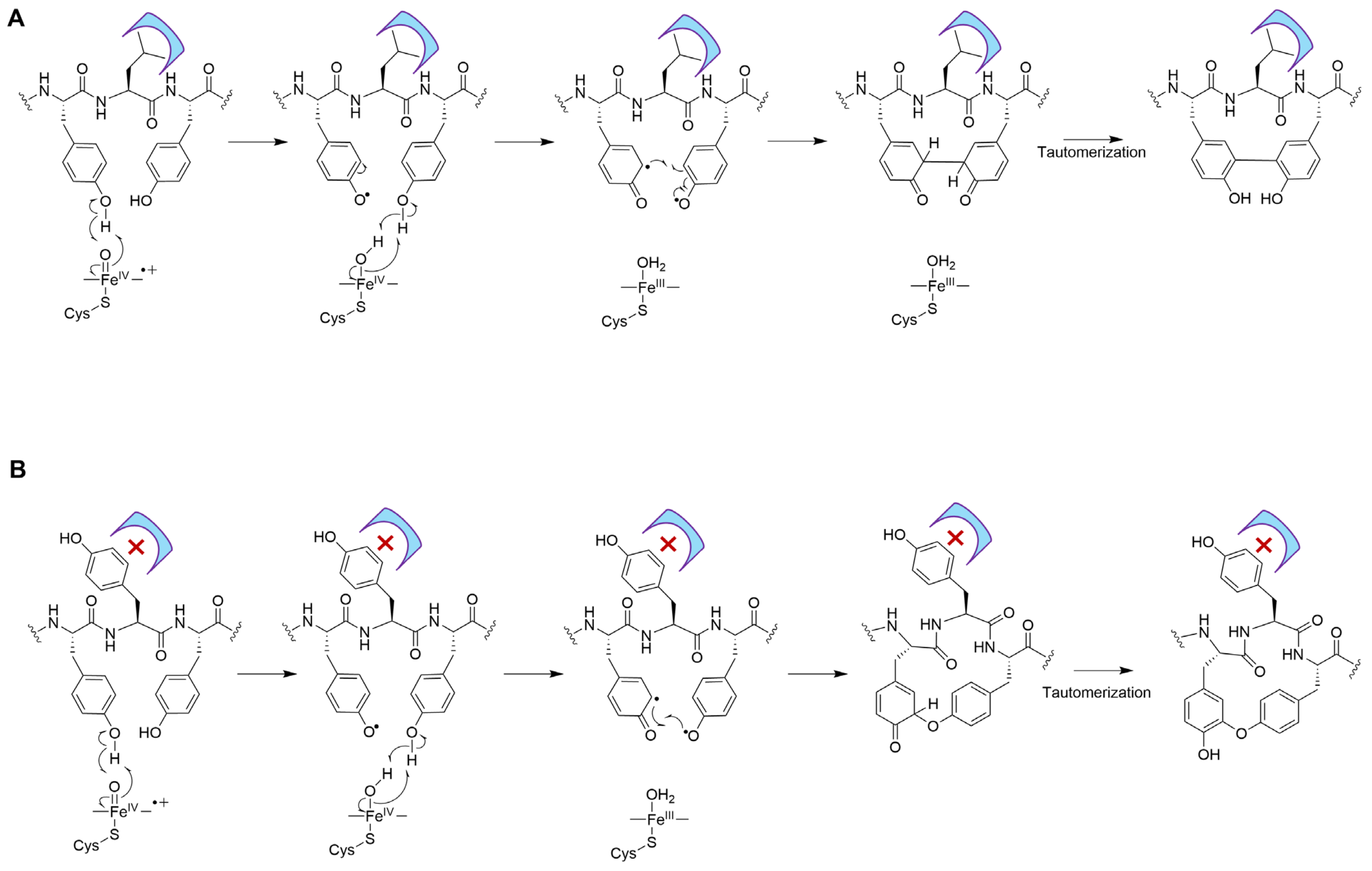

**Figure S37. Alternative mechanisms proposed in the literature for Tyr-Tyr crosslink formation that involves radical-radical couplings** [2-4]**.** Several studies have suggested that the crosslinking of two Tyr involves the coupling of two Tyr radicals, one formed by Compound I and one formed by Compound II. The mechanisms are arbitrarily drawn as first activating the N-terminal Tyr.

### Supplementary Tables

#### Table S1. Plasmid constructs used in this study.

Plasmid constructs and vector backbones used to amplify genes encoding precursors, 5’ and 3’ flanking sequences to identify location on the plasmid backbone, and the translated protein sequence of expressed genes. Precursor sequences are shown in green font. P450 coding sequence is shown in blue font. Peptide modification sites in the precursors are displayed in bold. N-terminal His6 tags and carried over amino acid sequences from the vector backbone due to the fusion are shown in purple font.

| Plasmid Construct | Vector Backbone | 5’ → 3’ sequence | | Flanking Sequences |
|---|---|---|---|---|
| **ApyA-ApyO** | pRSF-Duet | ORF | ATGGGCAGCAGCCATCACCATCATCACCACAGCCAGATGGCGACCAAACCAAAAAAGACCAAGGGCGTGAGCGTATCTATT<br>GAAGGGAAATTACCGAAAATGACGTTAGATATGCCTGTCGATGCAAAAAAAATCAAAGCTATTCAGAAATGTCTGGAGAACG<br>GTAAACTCACCATTACCATGTCTAAAGTAGATCTGGCAGGCGGTCGTATGGGGGAAGGTTATCTGTACGATTGAGATTCAAG<br>GAGATATACCATGAACGGTCTCCCGCCTTCTCTGCCTCGTGTGGACTCAACAGAAAGCCTGTTCGCGGAGCCTTTAGCATTC<br>TTAGCGCAAGCCCGTTCTCGTCACGGAGACGTGTTCGTTATGCGCGAGCACGGCCCCATCTTTTCCCGCGCATCGGATTGT<br>TCGGGTGTCATTGCTGTCTTTGGAGAGCACCGTTTACGTCAAATCCTTACGGACATCGACAACTTCGCGTTACCGATGAGTG<br>CCGCCGCCAAGATGGCATTACCTAAGAACCTCGTCAATTTAAACCGTGGCCTGCATAGTATGCGCGAGCCGGAGCATGGTC<br>GTCACAAGCGTTTGCTTACCGGTACAATTAACCGCGAGCTCTTTGACGCCCACCGTTTCGAAATTCGCGCAGCGCTCAACCG<br>CTTTTGTGAAATGCTCAAGGTAGATCGCCGTATCAGTGTGGTGAGCCGTATGCGCGAGCTGACGGTCGAGATGGCATCTCA<br>TATCTTTCTGGGTGCACAATGTCAGGAAGACGATGAGTTGGCGTTTCTTCTTTCAGCCTATTTCACCCTGCGCCGCGAGGCT<br>AGCTCATTAAACGCTCGTGACCCGTTGCTTTATCGTGACGAGCTGATCGGTGTGGGACAACAGCTGGACCGCACATTACGC<br>GAACGTATCCGCCGCTATCGTAAACGCCCCGTCGACGCCCGCGCCGGTCTGTTACAGCGTCTTGCTACGGCAGGTCCGCC<br>GGGCTCGCCTGCGTTGTCGGAAGATGAAATTGTGGGTCACGCCAACGTAATGTTCGTGTCTTCCACGGAGCCGGTTGCCAT<br>GTCTCTTACGTGGTTGTTGCTGGTCTTAAGTCAACTGCCTGACTTACGTCGTGCGCTGCGTGCGGAGATCGCGGACCGTGC<br>GTCTATGCCGGCCAGTACTAATGGCGCGTCGTGGTTAGAGAACGTTGTCAACGAGACGTTGCGCTTATTGACCCCAAATGC<br>CCTGATGGTCCGCGCCACAACACGCGCAGTGTCACTGCAAGGTGTTGCACTCCCGGCCCGCTGTGAGATTGTGGTATGTCC<br>ATTTCTGGCTCACCGCGAGGCCAAACCATTTCCCGACCCCCACGCATTCTCACCATCGCGCTGGGAGACCGCGCGTCCCAG<br>TCCTTATGAGTACTTTCCTTTCGGGGCGGGCGGACACTTCTGTGCGGGGCGTAATTTGGCCCTCTCACTTATTCGCGAAGTG<br>TTGTCGACGCTGTTAAGCCGTTTCGATTTCGTTCTTGACGGTGAGCAAAGCATCGACTGGCGCATTCATATTATGCTGATGCC<br>CAAGGGAGACCCGGCTTTGATTGCTCACCCTGTGGATGAACGTGGCGATACCCCGAGTCCAAAGTGGCGTGGGCCCATTAC<br>CGATTTATTTCACTTCGCCCCTGGGCTTTCATGA | 5’ Flank<br><br>TGTTTAACTT<br>TAATAAGGAG<br>ATAT<br><br>3’ Flank<br><br>CTCGAGTCTG<br>GTAAAGAAAC<br>CGCT |
| Translated sequence | ApyA: MGSSHHHHHHSQMATKPKKTKGVSVSIEGKLPKMTLDMPVDAKKIKAIQKCLENGKLTITMSKVDLAGGRMGEG**Y**L**Y**D<br>ApyO:<br>MNGLPPSLPRVDSTESLFAEPLAFLAQARSRHGDVFVMREHGPIFSRASDCSGVIAVFGEHRLRQILTDIDNFALPMSAAAKMALPKNLVNLNRGLHSMREPEHGRHKRLLTGTINR<br>ELFDAHRFEIRAALNRFCEMLKVDRRISVVSRMRELTVEMASHIFLGAQCQEDDELAFLLSAYFTLRREASSLNARDPLLYRDELIGVGQQLDRTLRERIRRYRKRPVDARAGLLQRL<br>ATAGPPGSPALSEDEIVGHANVMFVSSTEPVAMSLTWLLLVLSQLPDLRRALRAEIADRASMPASTNGASWLENVVNETLRLLTPNALMVRATTRAVSLQGVALPARCEIVVCPFLA<br>HREAKPFPDPHAFSPSRWETARPSPYEYFPFGAGGHFCAGRNLALSLIREVLSTLLSRFDFVLDGEQSIDWRIHIMLMPKGDPALIAHPVDERGDTPSPKWRGPITDLFHFAPGLS | | | |
| **ApyA-Y6W ApyO** | pRSF-Duet | ORF | ATGGGCAGCAGCCATCACCATCATCACCACAGCCAGATGGCGACCAAACCAAAAAAGACCAAGGGCGTGAGCGTATCTATT<br>GAAGGGAAATTACCGAAAATGACGTTAGATATGCCTGTCGATGCAAAAAAAATCAAAGCTATTCAGAAATGTCTGGAGAACG<br>GTAAACTCACCATTACCATGTCTAAAGTAGATCTGGCAGGCGGTCGTATGGGGGAAGGTTGGCTGTACGATTGAGATTCAAG<br>GAGATATACCATGAACGGTCTCCCGCCTTCTCTGCCTCGTGTGGACTCAACAGAAAGCCTGTTCGCGGAGCCTTTAGCATTC<br>TTAGCGCAAGCCCGTTCTCGTCACGGAGACGTGTTCGTTATGCGCGAGCACGGCCCCATCTTTTCCCGCGCATCGGATTGT<br>TCGGGTGTCATTGCTGTCTTTGGAGAGCACCGTTTACGTCAAATCCTTACGGACATCGACAACTTCGCGTTACCGATGAGTG<br>CCGCCGCCAAGATGGCATTACCTAAGAACCTCGTCAATTTAAACCGTGGCCTGCATAGTATGCGCGAGCCGGAGCATGGTC<br>GTCACAAGCGTTTGCTTACCGGTACAATTAACCGCGAGCTCTTTGACGCCCACCGTTTCGAAATTCGCGCAGCGCTCAACCG<br>CTTTTGTGAAATGCTCAAGGTAGATCGCCGTATCAGTGTGGTGAGCCGTATGCGCGAGCTGACGGTCGAGATGGCATCTCA<br>TATCTTTCTGGGTGCACAATGTCAGGAAGACGATGAGTTGGCGTTTCTTCTTTCAGCCTATTTCACCCTGCGCCGCGAGGCT<br>AGCTCATTAAACGCTCGTGACCCGTTGCTTTATCGTGACGAGCTGATCGGTGTGGGACAACAGCTGGACCGCACATTACGC<br>GAACGTATCCGCCGCTATCGTAAACGCCCCGTCGACGCCCGCGCCGGTCTGTTACAGCGTCTTGCTACGGCAGGTCCGCC<br>GGGCTCGCCTGCGTTGTCGGAAGATGAAATTGTGGGTCACGCCAACGTAATGTTCGTGTCTTCCACGGAGCCGGTTGCCAT<br>GTCTCTTACGTGGTTGTTGCTGGTCTTAAGTCAACTGCCTGACTTACGTCGTGCGCTGCGTGCGGAGATCGCGGACCGTGC<br>GTCTATGCCGGCCAGTACTAATGGCGCGTCGTGGTTAGAGAACGTTGTCAACGAGACGTTGCGCTTATTGACCCCAAATGC<br>CCTGATGGTCCGCGCCACAACACGCGCAGTGTCACTGCAAGGTGTTGCACTCCCGGCCCGCTGTGAGATTGTGGTATGTCC<br>ATTTCTGGCTCACCGCGAGGCCAAACCATTTCCCGACCCCCACGCATTCTCACCATCGCGCTGGGAGACCGCGCGTCCCAG<br>TCCTTATGAGTACTTTCCTTTCGGGGCGGGCGGACACTTCTGTGCGGGGCGTAATTTGGCCCTCTCACTTATTCGCGAAGTG<br>TTGTCGACGCTGTTAAGCCGTTTCGATTTCGTTCTTGACGGTGAGCAAAGCATCGACTGGCGCATTCATATTATGCTGATGCC<br>CAAGGGAGACCCGGCTTTGATTGCTCACCCTGTGGATGAACGTGGCGATACCCCGAGTCCAAAGTGGCGTGGGCCCATTAC<br>CGATTTATTTCACTTCGCCCCTGGGCTTTCATGA | 5’ Flank<br><br>TGTTTAACTT<br>TAATAAGGAG<br>ATAT<br><br>3’ Flank<br><br>CTCGAGTCTG<br>GTAAAGAAAC<br>CGCT |
| Translated sequence | ApyA-Y6W: MGSSHHHHHHSQMATKPKKTKGVSVSIEGKLPKMTLDMPVDAKKIKAIQKCLENGKLTITMSKVDLAGGRMGEG**W**L**Y**D<br>ApyO:<br>MNGLPPSLPRVDSTESLFAEPLAFLAQARSRHGDVFVMREHGPIFSRASDCSGVIAVFGEHRLRQILTDIDNFALPMSAAAKMALPKNLVNLNRGLHSMREPEHGRHKRLLTGTINR<br>ELFDAHRFEIRAALNRFCEMLKVDRRISVVSRMRELTVEMASHIFLGAQCQEDDELAFLLSAYFTLRREASSLNARDPLLYRDELIGVGQQLDRTLRERIRRYRKRPVDARAGLLQRL<br>ATAGPPGSPALSEDEIVGHANVMFVSSTEPVAMSLTWLLLVLSQLPDLRRALRAEIADRASMPASTNGASWLENVVNETLRLLTPNALMVRATTRAVSLQGVALPARCEIVVCPFLA<br>HREAKPFPDPHAFSPSRWETARPSPYEYFPFGAGGHFCAGRNLALSLIREVLSTLLSRFDFVLDGEQSIDWRIHIMLMPKGDPALIAHPVDERGDTPSPKWRGPITDLFHFAPGLS | | | |
| **ApyA-Y8W ApyO** | pRSF-Duet | ORF | ATGGGCAGCAGCCATCACCATCATCACCACAGCCAGATGGCGACCAAACCAAAAAAGACCAAGGGCGTGAGCGTATCTATT<br>GAAGGGAAATTACCGAAAATGACGTTAGATATGCCTGTCGATGCAAAAAAAATCAAAGCTATTCAGAAATGTCTGGAGAACG<br>GTAAACTCACCATTACCATGTCTAAAGTAGATCTGGCAGGCGGTCGTATGGGGGAAGGTTATCTGTGGGATTGAGATTCAAG<br>GAGATATACCATGAACGGTCTCCCGCCTTCTCTGCCTCGTGTGGACTCAACAGAAAGCCTGTTCGCGGAGCCTTTAGCATTC<br>TTAGCGCAAGCCCGTTCTCGTCACGGAGACGTGTTCGTTATGCGCGAGCACGGCCCCATCTTTTCCCGCGCATCGGATTGT<br>TCGGGTGTCATTGCTGTCTTTGGAGAGCACCGTTTACGTCAAATCCTTACGGACATCGACAACTTCGCGTTACCGATGAGTG<br>CCGCCGCCAAGATGGCATTACCTAAGAACCTCGTCAATTTAAACCGTGGCCTGCATAGTATGCGCGAGCCGGAGCATGGTC<br>GTCACAAGCGTTTGCTTACCGGTACAATTAACCGCGAGCTCTTTGACGCCCACCGTTTCGAAATTCGCGCAGCGCTCAACCG<br>CTTTTGTGAAATGCTCAAGGTAGATCGCCGTATCAGTGTGGTGAGCCGTATGCGCGAGCTGACGGTCGAGATGGCATCTCA<br>TATCTTTCTGGGTGCACAATGTCAGGAAGACGATGAGTTGGCGTTTCTTCTTTCAGCCTATTTCACCCTGCGCCGCGAGGCT<br>AGCTCATTAAACGCTCGTGACCCGTTGCTTTATCGTGACGAGCTGATCGGTGTGGGACAACAGCTGGACCGCACATTACGC<br>GAACGTATCCGCCGCTATCGTAAACGCCCCGTCGACGCCCGCGCCGGTCTGTTACAGCGTCTTGCTACGGCAGGTCCGCC<br>GGGCTCGCCTGCGTTGTCGGAAGATGAAATTGTGGGTCACGCCAACGTAATGTTCGTGTCTTCCACGGAGCCGGTTGCCAT<br>GTCTCTTACGTGGTTGTTGCTGGTCTTAAGTCAACTGCCTGACTTACGTCGTGCGCTGCGTGCGGAGATCGCGGACCGTGC<br>GTCTATGCCGGCCAGTACTAATGGCGCGTCGTGGTTAGAGAACGTTGTCAACGAGACGTTGCGCTTATTGACCCCAAATGC<br>CCTGATGGTCCGCGCCACAACACGCGCAGTGTCACTGCAAGGTGTTGCACTCCCGGCCCGCTGTGAGATTGTGGTATGTCC<br>ATTTCTGGCTCACCGCGAGGCCAAACCATTTCCCGACCCCCACGCATTCTCACCATCGCGCTGGGAGACCGCGCGTCCCAG<br>TCCTTATGAGTACTTTCCTTTCGGGGCGGGCGGACACTTCTGTGCGGGGCGTAATTTGGCCCTCTCACTTATTCGCGAAGTG<br>TTGTCGACGCTGTTAAGCCGTTTCGATTTCGTTCTTGACGGTGAGCAAAGCATCGACTGGCGCATTCATATTATGCTGATGCC<br>CAAGGGAGACCCGGCTTTGATTGCTCACCCTGTGGATGAACGTGGCGATACCCCGAGTCCAAAGTGGCGTGGGCCCATTAC<br>CGATTTATTTCACTTCGCCCCTGGGCTTTCATGA | 5’ Flank<br><br>TGTTTAACTT<br>TAATAAGGAG<br>ATAT<br><br>3’ Flank<br><br>CTCGAGTCTG<br>GTAAAGAAAC<br>CGCT |
| Translated sequence | ApyA-Y8W: MGSSHHHHHHSQMATKPKKTKGVSVSIEGKLPKMTLDMPVDAKKIKAIQKCLENGKLTITMSKVDLAGGRMGEG**Y**L**W**D<br>ApyO:<br>MNGLPPSLPRVDSTESLFAEPLAFLAQARSRHGDVFVMREHGPIFSRASDCSGVIAVFGEHRLRQILTDIDNFALPMSAAAKMALPKNLVNLNRGLHSMREPE | | | |

| | | | | |
|---|---|---|---|---|
| | HGRHKRLLTGTINRELFDAHRFEIRAALNRFCEMLKVDRRISVVSRMRELTVEMASHIFLGAQCQEDDELAFLLSAYFTLRREASSLNARDPLLYRDELIGVGQQ<br>LDRTLRERIRRYRKRPVDARAGLLQRLATAGPPGSPALSEDEIVGHANVMFVSSTEPVAMSLTWLLLVLSQLPDLRRALRAEIADRASMPASTNGASWLENVV<br>NETLRLLTPNALMVRATTRAVSLQGVALPARCEIVVCPFLAHREAKPFPDPHAFSPSRWETARPSPYEYFPFGAGGHFCAGRNLALSLIREVLSTLLSRFDFVL<br>DGEQSIDWRIHIMLMPKGDPALIAHPVDERGDTPSPKWRGPITDLFHFAPGLS | | | |
| **ApyA-Y6W/Y8W ApyO** | pRSF-Duet | ORF | ATGGGCAGCAGCCATCACCATCATCACCACAGCCAGATGGCGACCAAACCAAAAAAGACCAAGGGCGTGAGCGTATCTATT<br>GAAGGGAAATTACCGAAAATGACGTTAGATATGCCTGTCGATGCAAAAAAAATCAAAGCTATTCAGAAATGTCTGGAGAACG<br>GTAAACTCACCATTACCATGTCTAAAGTAGATCTGGCAGGCGGTCGTATGGGGGAAGGTTGGCTGTGGGATTGAGATTCAAG<br>GAGATATACCATGAACGGTCTCCCGCCTTCTCTGCCTCGTGTGGACTCAACAGAAAGCCTGTTCGCGGAGCCTTTAGCATTC<br>TTAGCGCAAGCCCGTTCTCGTCACGGAGACGTGTTCGTTATGCGCGAGCACGGCCCCATCTTTTCCCGCGCATCGGATTGT<br>TCGGGTGTCATTGCTGTCTTTGGAGAGCACCGTTTACGTCAAATCCTTACGGACATCGACAACTTCGCGTTACCGATGAGTG<br>CCGCCGCCAAGATGGCATTACCTAAGAACCTCGTCAATTTAAACCGTGGCCTGCATAGTATGCGCGAGCCGGAGCATGGTC<br>GTCACAAGCGTTTGCTTACCGGTACAATTAACCGCGAGCTCTTTGACGCCCACCGTTTCGAAATTCGCGCAGCGCTCAACCG<br>CTTTTGTGAAATGCTCAAGGTAGATCGCCGTATCAGTGTGGTGAGCCGTATGCGCGAGCTGACGGTCGAGATGGCATCTCA<br>TATCTTTCTGGGTGCACAATGTCAGGAAGACGATGAGTTGGCGTTTCTTCTTTCAGCCTATTTCACCCTGCGCCGCGAGGCT<br>AGCTCATTAAACGCTCGTGACCCGTTGCTTTATCGTGACGAGCTGATCGGTGTGGGACAACAGCTGGACCGCACATTACGC<br>GAACGTATCCGCCGCTATCGTAAACGCCCCGTCGACGCCCGCGCCGGTCTGTTACAGCGTCTTGCTACGGCAGGTCCGCC<br>GGGCTCGCCTGCGTTGTCGGAAGATGAAATTGTGGGTCACGCCAACGTAATGTTCGTGTCTTCCACGGAGCCGGTTGCCAT<br>GTCTCTTACGTGGTTGTTGCTGGTCTTAAGTCAACTGCCTGACTTACGTCGTGCGCTGCGTGCGGAGATCGCGGACCGTGC<br>GTCTATGCCGGCCAGTACTAATGGCGCGTCGTGGTTAGAGAACGTTGTCAACGAGACGTTGCGCTTATTGACCCCAAATGC<br>CCTGATGGTCCGCGCCACAACACGCGCAGTGTCACTGCAAGGTGTTGCACTCCCGGCCCGCTGTGAGATTGTGGTATGTCC<br>ATTTCTGGCTCACCGCGAGGCCAAACCATTTCCCGACCCCCACGCATTCTCACCATCGCGCTGGGAGACCGCGCGTCCCAG<br>TCCTTATGAGTACTTTCCTTTCGGGGCGGGCGGACACTTCTGTGCGGGGCGTAATTTGGCCCTCTCACTTATTCGCGAAGTG<br>TTGTCGACGCTGTTAAGCCGTTTCGATTTCGTTCTTGACGGTGAGCAAAGCATCGACTGGCGCATTCATATTATGCTGATGCC<br>CAAGGGAGACCCGGCTTTGATTGCTCACCCTGTGGATGAACGTGGCGATACCCCGAGTCCAAAGTGGCGTGGGCCCATTAC<br>CGATTTATTTCACTTCGCCCCTGGGCTTTCATGA | 5' Flank<br>TGTTTAACTTTAATAAGGAGATAT<br><br>3' Flank<br>CTCGAGTCTGGTAAAGAAACCGCT |
| Translated sequence | ApyA-Y6W/Y8W: MGSSHHHHHHSQMATKPKKTKGVSVSIEGKLPKMTLDMPVDAKKIKAIQKCLENGKLTITMSKVDLAGGRMGEG**W**L**W**D<br>ApyO:<br>MNGLPPSLPRVDSTESLFAEPLAFLAQARSRHGDVFVMREHGPIFSRASDCSGVIAVFGEHRLRQILTDIDNFALPMSAAAKMALPKNLVNLNRGLHSMREPEHGRHKRLLTGTINR<br>ELFDAHRFEIRAALNRFCEMLKVDRRISVVSRMRELTVEMASHIFLGAQCQEDDELAFLLSAYFTLRREASSLNARDPLLYRDELIGVGQQLDRTLRERIRRYRKRPVDARAGLLQRL<br>ATAGPPGSPALSEDEIVGHANVMFVSSTEPVAMSLTWLLLVLSQLPDLRRALRAEIADRASMPASTNGASWLENVVNETLRLLTPNALMVRATTRAVSLQGVALPARCEIVVCPFLA<br>HREAKPFPDPHAFSPSRWETARPSPYEYFPFGAGGHFCAGRNLALSLIREVLSTLLSRFDFVLDGEQSIDWRIHIMLMPKGDPALIAHPVDERGDTPSPKWRGPITDLFHFAPGLS | | | |
| **ApyA-L7Y ApyO** | pRSF-Duet | ORF | ATGGGCAGCAGCCATCACCATCATCACCACAGCCAGATGGCGACCAAACCAAAAAAGACCAAGGGCGTGAGCGTATCTATT<br>GAAGGGAAATTACCGAAAATGACGTTAGATATGCCTGTCGATGCAAAAAAAATCAAAGCTATTCAGAAATGTCTGGAGAACG<br>GTAAACTCACCATTACCATGTCTAAAGTAGATCTGGCAGGCGGTCGTATGGGGGAAGGTTACTACTATGATTGAGATTCAAG<br>GAGATATACCATGAACGGTCTCCCGCCTTCTCTGCCTCGTGTGGACTCAACAGAAAGCCTGTTCGCGGAGCCTTTAGCATTC<br>TTAGCGCAAGCCCGTTCTCGTCACGGAGACGTGTTCGTTATGCGCGAGCACGGCCCCATCTTTTCCCGCGCATCGGATTGT<br>TCGGGTGTCATTGCTGTCTTTGGAGAGCACCGTTTACGTCAAATCCTTACGGACATCGACAACTTCGCGTTACCGATGAGTG<br>CCGCCGCCAAGATGGCATTACCTAAGAACCTCGTCAATTTAAACCGTGGCCTGCATAGTATGCGCGAGCCGGAGCATGGTC<br>GTCACAAGCGTTTGCTTACCGGTACAATTAACCGCGAGCTCTTTGACGCCCACCGTTTCGAAATTCGCGCAGCGCTCAACCG<br>CTTTTGTGAAATGCTCAAGGTAGATCGCCGTATCAGTGTGGTGAGCCGTATGCGCGAGCTGACGGTCGAGATGGCATCTCA<br>TATCTTTCTGGGTGCACAATGTCAGGAAGACGATGAGTTGGCGTTTCTTCTTTCAGCCTATTTCACCCTGCGCCGCGAGGCT<br>AGCTCATTAAACGCTCGTGACCCGTTGCTTTATCGTGACGAGCTGATCGGTGTGGGACAACAGCTGGACCGCACATTACGC<br>GAACGTATCCGCCGCTATCGTAAACGCCCCGTCGACGCCCGCGCCGGTCTGTTACAGCGTCTTGCTACGGCAGGTCCGCC<br>GGGCTCGCCTGCGTTGTCGGAAGATGAAATTGTGGGTCACGCCAACGTAATGTTCGTGTCTTCCACGGAGCCGGTTGCCAT<br>GTCTCTTACGTGGTTGTTGCTGGTCTTAAGTCAACTGCCTGACTTACGTCGTGCGCTGCGTGCGGAGATCGCGGACCGTGC<br>GTCTATGCCGGCCAGTACTAATGGCGCGTCGTGGTTAGAGAACGTTGTCAACGAGACGTTGCGCTTATTGACCCCAAATGC<br>CCTGATGGTCCGCGCCACAACACGCGCAGTGTCACTGCAAGGTGTTGCACTCCCGGCCCGCTGTGAGATTGTGGTATGTCC<br>ATTTCTGGCTCACCGCGAGGCCAAACCATTTCCCGACCCCCACGCATTCTCACCATCGCGCTGGGAGACCGCGCGTCCCAG<br>TCCTTATGAGTACTTTCCTTTCGGGGCGGGCGGACACTTCTGTGCGGGGCGTAATTTGGCCCTCTCACTTATTCGCGAAGTG<br>TTGTCGACGCTGTTAAGCCGTTTCGATTTCGTTCTTGACGGTGAGCAAAGCATCGACTGGCGCATTCATATTATGCTGATGCC<br>CAAGGGAGACCCGGCTTTGATTGCTCACCCTGTGGATGAACGTGGCGATACCCCGAGTCCAAAGTGGCGTGGGCCCATTAC<br>CGATTTATTTCACTTCGCCCCTGGGCTTTCATGA | 5' Flank<br>TGTTTAACTTTAATAAGGAGATAT<br><br>3' Flank<br>CTCGAGTCTGGTAAAGAAACCGCT |
| Translated sequence | ApyA-L7Y: MGSSHHHHHHSQMATKPKKTKGVSVSIEGKLPKMTLDMPVDAKKIKAIQKCLENGKLTITMSKVDLAGGRMGEG**YYY**D<br>ApyO:<br>MNGLPPSLPRVDSTESLFAEPLAFLAQARSRHGDVFVMREHGPIFSRASDCSGVIAVFGEHRLRQILTDIDNFALPMSAAAKMALPKNLVNLNRGLHSMREPE<br>HGRHKRLLTGTINRELFDAHRFEIRAALNRFCEMLKVDRRISVVSRMRELTVEMASHIFLGAQCQEDDELAFLLSAYFTLRREASSLNARDPLLYRDELIGVGQQ<br>LDRTLRERIRRYRKRPVDARAGLLQRLATAGPPGSPALSEDEIVGHANVMFVSSTEPVAMSLTWLLLVLSQLPDLRRALRAEIADRASMPASTNGASWLENVV<br>NETLRLLTPNALMVRATTRAVSLQGVALPARCEIVVCPFLAHREAKPFPDPHAFSPSRWETARPSPYEYFPFGAGGHFCAGRNLALSLIREVLSTLLSRFDFVL<br>DGEQSIDWRIHIMLMPKGDPALIAHPVDERGDTPSPKWRGPITDLFHFAPGLS | | | |
| **ApyA-L7W ApyO** | pRSF-Duet | ORF | ATGGGCAGCAGCCATCACCATCATCACCACAGCCAGATGGCGACCAAACCAAAAAAGACCAAGGGCGTGAGCGTATCTATT<br>GAAGGGAAATTACCGAAAATGACGTTAGATATGCCTGTCGATGCAAAAAAAATCAAAGCTATTCAGAAATGTCTGGAGAACG<br>GTAAACTCACCATTACCATGTCTAAAGTAGATCTGGCAGGCGGTCGTATGGGGGAAGGTTACTGGTATGATTGAGATTCAAG<br>GAGATATACCATGAACGGTCTCCCGCCTTCTCTGCCTCGTGTGGACTCAACAGAAAGCCTGTTCGCGGAGCCTTTAGCATTC<br>TTAGCGCAAGCCCGTTCTCGTCACGGAGACGTGTTCGTTATGCGCGAGCACGGCCCCATCTTTTCCCGCGCATCGGATTGT<br>TCGGGTGTCATTGCTGTCTTTGGAGAGCACCGTTTACGTCAAATCCTTACGGACATCGACAACTTCGCGTTACCGATGAGTG<br>CCGCCGCCAAGATGGCATTACCTAAGAACCTCGTCAATTTAAACCGTGGCCTGCATAGTATGCGCGAGCCGGAGCATGGTC<br>GTCACAAGCGTTTGCTTACCGGTACAATTAACCGCGAGCTCTTTGACGCCCACCGTTTCGAAATTCGCGCAGCGCTCAACCG<br>CTTTTGTGAAATGCTCAAGGTAGATCGCCGTATCAGTGTGGTGAGCCGTATGCGCGAGCTGACGGTCGAGATGGCATCTCA<br>TATCTTTCTGGGTGCACAATGTCAGGAAGACGATGAGTTGGCGTTTCTTCTTTCAGCCTATTTCACCCTGCGCCGCGAGGCT<br>AGCTCATTAAACGCTCGTGACCCGTTGCTTTATCGTGACGAGCTGATCGGTGTGGGACAACAGCTGGACCGCACATTACGC<br>GAACGTATCCGCCGCTATCGTAAACGCCCCGTCGACGCCCGCGCCGGTCTGTTACAGCGTCTTGCTACGGCAGGTCCGCC<br>GGGCTCGCCTGCGTTGTCGGAAGATGAAATTGTGGGTCACGCCAACGTAATGTTCGTGTCTTCCACGGAGCCGGTTGCCAT<br>GTCTCTTACGTGGTTGTTGCTGGTCTTAAGTCAACTGCCTGACTTACGTCGTGCGCTGCGTGCGGAGATCGCGGACCGTGC<br>GTCTATGCCGGCCAGTACTAATGGCGCGTCGTGGTTAGAGAACGTTGTCAACGAGACGTTGCGCTTATTGACCCCAAATGC<br>CCTGATGGTCCGCGCCACAACACGCGCAGTGTCACTGCAAGGTGTTGCACTCCCGGCCCGCTGTGAGATTGTGGTATGTCC<br>ATTTCTGGCTCACCGCGAGGCCAAACCATTTCCCGACCCCCACGCATTCTCACCATCGCGCTGGGAGACCGCGCGTCCCAG<br>TCCTTATGAGTACTTTCCTTTCGGGGCGGGCGGACACTTCTGTGCGGGGCGTAATTTGGCCCTCTCACTTATTCGCGAAGTG<br>TTGTCGACGCTGTTAAGCCGTTTCGATTTCGTTCTTGACGGTGAGCAAAGCATCGACTGGCGCATTCATATTATGCTGATGCC<br>CAAGGGAGACCCGGCTTTGATTGCTCACCCTGTGGATGAACGTGGCGATACCCCGAGTCCAAAGTGGCGTGGGCCCATTAC<br>CGATTTATTTCACTTCGCCCCTGGGCTTTCATGA | 5' Flank<br>TGTTTAACTTTAATAAGGAGATAT<br><br>3' Flank<br>CTCGAGTCTGGTAAAGAAACCGCT |
| Translated sequence | ApyA-L7W: MGSSHHHHHHSQMATKPKKTKGVSVSIEGKLPKMTLDMPVDAKKIKAIQKCLENGKLTITMSKVDLAGGRMGEG**Y**W**Y**D<br>ApyO:<br>MNGLPPSLPRVDSTESLFAEPLAFLAQARSRHGDVFVMREHGPIFSRASDCSGVIAVFGEHRLRQILTDIDNFALPMSAAAKMALPKNLVNLNRGLHSMREPEHGRHKRLLTGTINR<br>ELFDAHRFEIRAALNRFCEMLKVDRRISVVSRMRELTVEMASHIFLGAQCQEDDELAFLLSAYFTLRREASSLNARDPLLYRDELIGVGQQLDRTLRERIRRYRKRPVDARAGLLQRL<br>ATAGPPGSPALSEDEIVGHANVMFVSSTEPVAMSLTWLLLVLSQLPDLRRALRAEIADRASMPASTNGASWLENVVNETLRLLTPNALMVRATTRAVSLQGVALPARCEIVVCPFLA<br>HREAKPFPDPHAFSPSRWETARPSPYEYFPFGAGGHFCAGRNLALSLIREVLSTLLSRFDFVLDGEQSIDWRIHIMLMPKGDPALIAHPVDERGDTPSPKWRGPITDLFHFAPGLS | | | |
| **pRSF-6xHis-TEV-ApyO** | pRSF-Duet | ORF | ATGGGCAGCAGCCATCACCATCATCACCACAGCCAGGAGAACCTGTACTTCCAATCCATGAACGGTCTCCCGCCTTCTCTGC<br>CTCGTGTGGACTCAACAGAAAGCCTGTTCGCGGAGCCTTTAGCATTCTTAGCGCAAGCCCGTTCTCGTCACGGAGACGTGTT<br>CGTTATGCGCGAGCACGGCCCCATCTTTTCCCGCGCATCGGATTGTTCGGGTGTCATTGCTGTCTTTGGAGAGCACCGTTTA<br>CGTCAAATCCTTACGGACATCGACAACTTCGCGTTACCGATGAGTGCCGCCGCCAAGATGGCATTACCTAAGAACCTCGTCA<br>ATTTAAACCGTGGCCTGCATAGTATGCGCGAGCCGGAGCATGGTCGTCACAAGCGTTTGCTTACCGGTACAATTAACCGCGA<br>GCTCTTTGACGCCCACCGTTTCGAAATTCGCGCAGCGCTCAACCGCTTTTGTGAAATGCTCAAGGTAGATCGCCGTATCAGT<br>GTGGTGAGCCGTATGCGCGAGCTGACGGTCGAGATGGCATCTCATATCTTTCTGGGTGCACAATGTCAGGAAGACGATGAG<br>TTGGCGTTTCTTCTTTCAGCCTATTTCACCCTGCGCCGCGAGGCTAGCTCATTAAACGCTCGTGACCCGTTGCTTTATCGTGA<br>CGAGCTGATCGGTGTGGGACAACAGCTGGACCGCACATTACGCGAACGTATCCGCCGCTATCGTAAACGCCCCGTCGACG<br>CCCGCGCCGGTCTGTTACAGCGTCTTGCTACGGCAGGTCCGCCGGGCTCGCCTGCGTTGTCGGAAGATGAAATTGTGGGT<br>CACGCCAACGTAATGTTCGTGTCTTCCACGGAGCCGGTTGCCATGTCTCTTACGTGGTTGTTGCTGGTCTTAAGTCAACTGC<br>CTGACTTACGTCGTGCGCTGCGTGCGGAGATCGCGGACCGTGCGTCTATGCCGGCCAGTACTAATGGCGCGTCGTGGTTA<br>GAGAACGTTGTCAACGAGACGTTGCGCTTATTGACCCCAAATGCCCTGATGGTCCGCGCCACAACACGCGCAGTGTCACTG<br>CAAGGTGTTGCACTCCCGGCCCGCTGTGAGATTGTGGTATGTCCATTTCTGGCTCACCGCGAGGCCAAACCATTTCCCGAC<br>CCCCACGCATTCTCACCATCGCGCTGGGAGACCGCGCGTCCCAGTCCTTATGAGTACTTTCCTTTCGGGGCGGGCGGACAC | 5' Flank<br>AAATAATTTTGTTTAACTTTAATAAGGAGATATACC<br><br>3' Flank<br>CTCGAGTCTGGTAAAGAAACCGCTGCTGCGAAATTTG |

| | | | | |
|---|---|---|---|---|
| | | | TTCTGTGCGGGGCGTAATTTGGCCCTCTCACTTATTCGCGAAGTGTTGTCGACGCTGTTAAGCCGTTTCGATTTCGTTCTTGA<br>CGGTGAGCAAAGCATCGACTGGCGCATTCATATTATGCTGATGCCCAAGGGAGACCCGGCTTTGATTGCTCACCCTGTGGA<br>TGAACGTGGCGATACCCCGAGTCCAAAGTGGCGTGGGCCCATTACCGATTTATTTCACTTCGCCCCTGGGCTTTCATGA | |
| Translated sequence | MGSSHHHHHHSQENLYFQSMNGLPPSLPRVDSTESLFAEPLAFLAQARSRHGDVFVMREHGPIFSRASDCSGVIAVFGEHRLRQILTDIDNFALPMSAAAKMALPKNLVNLNRGLH<br>SMREPEHGRHKRLLTGTINRELFDAHRFEIRAALNRFCEMLKVDRRISVVSRMRELTVEMASHIFLGAQCQEDDELAFLLSAYFTLRREASSLNARDPLLYRDELIGVGQQLDRTLRE<br>RIRRYRKRPVDARAGLLQRLATAGPPGSPALSEDEIVGHANVMFVSSTEPVAMSLTWLLLVLSQLPDLRRALRAEIADRASMPASTNGASWLENVVNETLRLLTPNALMVRATTRAV<br>SLQGVALPARCEIVVCPFLAHREAKPFPDPHAFSPSRWETARPSPYEYFPFGAGGHFCAGRNLALSLIREVLSTLLSRFDFVLDGEQSIDWRIHIMLMPKGDPALIAHPVDERGDTPS<br>PKWRGPITDLFHFAPGLS | | | |
| **pRSF-6xHis-MBP-ApyO** | pRSF-Duet | ORF | ATGGGCAGCAGCCATCATCATCATCATCACAGCAGCGGCCTGGTGCCGCGCGGCAGCCATATGAAAATCGAAGAAGGTAAA<br>CTGGTAATCTGGATTAACGGCGATAAAGGCTATAACGGTCTCGCTGAAGTCGGTAAGAAATTCGAGAAAGATACCGGAATTA<br>AAGTCACCGTTGAGCATCCGGATAAACTGGAAGAGAAATTCCCACAGGTTGCGGCAACTGGCGATGGCCCTGACATTATCTT<br>CTGGGCACACGACCGCTTTGGTGGCTACGCTCAATCTGGCCTGTTGGCTGAAATCACCCCGGACAAAGCGTTCCAGGACAA<br>GCTGTATCCGTTTACCTGGGATGCCGTACGTTACAACGGCAAGCTGATTGCTTACCCGATCGCTGTTGAAGCGTTATCGCTG<br>ATTTATAACAAAGATCTGCTGCCGAACCCGCCAAAAACCTGGGAAGAGATCCCGGCGCTGGATAAAGAACTGAAAGCGAAA<br>GGTAAGAGCGCGCTGATGTTCAACCTGCAAGAACCGTACTTCACCTGGCCGCTGATTGCTGCTGACGGGGGTTATGCGTTC<br>AAGTATGAAAACGGCAAGTACGACATTAAAGACGTGGGCGTGGATAACGCTGGCGCGAAAGCGGGTCTGACCTTCCTGGTT<br>GACCTGATTAAAAACAAACACATGAATGCAGACACCGATTACTCCATCGCAGAAGCTGCCTTTAATAAAGGCGAAACAGCGA<br>TGACCATCAACGGCCCGTGGGCATGGTCCAACATCGACACCAGCAAAGTGAATTATGGTGTAACGGTACTGCCGACCTTCA<br>AGGGTCAACCATCCAAACCGTTCGTTGGCGTGCTGAGCGCAGGTATTAACGCCGCCAGTCCGAACAAAGAGCTGGCGAAAG<br>AGTTCCTCGAAAACTATCTGCTGACTGATGAAGGTCTGGAAGCGGTTAATAAAGACAAACCGCTGGGTGCCGTAGCGCTGAA<br>GTCTTACGAGGAAGAGTTGGCGAAAGATCCACGTATTGCCGCCACCATGGAAAACGCCCAGAAAGGTGAAATCATGCCGAA<br>CATCCCGCAGATGTCCGCTTTCTGGTATGCCGTGCGTACTGCGGTGATCAACGCCGCCAGCGGTCGTCAGACTGTCGATGA<br>AGCCCTGAAAGACGCGCAGACTAATTCGAGCTCCCACCATCACCATCACCACGCGAATTCGGTACCGCTGGTTCCGCGTGG<br>ATCTGAGAACCTGTACTTCCAATCCGGATCCATGAACGGTCTCCCGCCTTCTCTGCCTCGTGTGGACTCAACAGAAAGCCTG<br>TTCGCGGAGCCTTTAGCATTCTTAGCGCAAGCCCGTTCTCGTCACGGAGACGTGTTCGTTATGCGCGAGCACGGCCCCATC<br>TTTTCCCGCGCATCGGATTGTTCGGGTGTCATTGCTGTCTTTGGAGAGCACCGTTTACGTCAAATCCTTACGGACATCGACA<br>ACTTCGCGTTACCGATGAGTGCCGCCGCCAAGATGGCATTACCTAAGAACCTCGTCAATTTAAACCGTGGCCTGCATAGTAT<br>GCGCGAGCCGGAGCATGGTCGTCACAAGCGTTTGCTTACCGGTACAATTAACCGCGAGCTCTTTGACGCCCACCGTTTCGA<br>AATTCGCGCAGCGCTCAACCGCTTTTGTGAAATGCTCAAGGTAGATCGCCGTATCAGTGTGGTGAGCCGTATGCGCGAGCT<br>GACGGTCGAGATGGCATCTCATATCTTTCTGGGTGCACAATGTCAGGAAGACGATGAGTTGGCGTTTCTTCTTTCAGCCTATT<br>TCACCCTGCGCCGCGAGGCTAGCTCATTAAACGCTCGTGACCCGTTGCTTTATCGTGACGAGCTGATCGGTGTGGGACAAC<br>AGCTGGACCGCACATTACGCGAACGTATCCGCCGCTATCGTAAACGCCCCGTCGACGCCCGCGCCGGTCTGTTACAGCGT<br>CTTGCTACGGCAGGTCCGCCGGGCTCGCCTGCGTTGTCGGAAGATGAAATTGTGGGTCACGCCAACGTAATGTTCGTGTCT<br>TCCACGGAGCCGGTTGCCATGTCTCTTACGTGGTTGTTGCTGGTCTTAAGTCAACTGCCTGACTTACGTCGTGCGCTGCGTG<br>CGGAGATCGCGGACCGTGCGTCTATGCCGGCCAGTACTAATGGCGCGTCGTGGTTAGAGAACGTTGTCAACGAGACGTTG<br>CGCTTATTGACCCCAAATGCCCTGATGGTCCGCGCCACAACACGCGCAGTGTCACTGCAAGGTGTTGCACTCCCGGCCCGC<br>TGTGAGATTGTGGTATGTCCATTTCTGGCTCACCGCGAGGCCAAACCATTTCCCGACCCCCACGCATTCTCACCATCGCGCT<br>GGGAGACCGCGCGTCCCAGTCCTTATGAGTACTTTCCTTTCGGGGCGGGCGGACACTTCTGTGCGGGGCGTAATTTGGCC<br>CTCTCACTTATTCGCGAAGTGTTGTCGACGCTGTTAAGCCGTTTCGATTTCGTTCTTGACGGTGAGCAAAGCATCGACTGGC<br>GCATTCATATTATGCTGATGCCCAAGGGAGACCCGGCTTTGATTGCTCACCCTGTGGATGAACGTGGCGATACCCCGAGTC<br>CAAAGTGGCGTGGGCCCATTACCGATTTATTTCACTTCGCCCCTGGGCTTTCATGA | Cloning sites<br><br>5': NcoI<br>3': XhoI<br><br>MBP Tag in green font |
| Translated sequence | MGSSHHHHHHSSGLVPRGSHMKIEEGKLVIWINGDKGYNGLAEVGKKFEKDTGIKVTVEHPDKLEEKFPQVAATGDGPDIIFWAHDRFGGYAQSGLLAEITPDKAFQDKLYPFTWD<br>AVRYNGKLIAYPIAVEALSLIYNKDLLPNPPKTWEEIPALDKELKAKGKSALMFNLQEPYFTWPLIAADGGYAFKYENGKYDIKDVGVDNAGAKAGLTFLVDLIKNKHMNADTDYSIAEA<br>AFNKGETAMTINGPWAWSNIDTSKVNYGVTVLPTFKGQPSKPFVGVLSAGINAASPNKELAKEFLENYLLTDEGLEAVNKDKPLGAVALKSYEEELAKDPRIAATMENAQKGEIMPNI<br>PQMSAFWYAVRTAVINAASGRQTVDEALKDAQTNSSSHHHHHHANSVPLVPRGSENLYFQSGSMNGLPPSLPRVDSTESLFAEPLAFLAQARSRHGDVFVMREHGPIFSRASDCS<br>GVIAVFGEHRLRQILTDIDNFALPMSAAAKMALPKNLVNLNRGLHSMREPEHGRHKRLLTGTINRELFDAHRFEIRAALNRFCEMLKVDRRISVVSRMRELTVEMASHIFLGAQCQED<br>DELAFLLSAYFTLRREASSLNARDPLLYRDELIGVGQQLDRTLRERIRRYRKRPVDARAGLLQRLATAGPPGSPALSEDEIVGHANVMFVSSTEPVAMSLTWLLLVLSQLPDLRRALR<br>AEIADRASMPASTNGASWLENVVNETLRLLTPNALMVRATTRAVSLQGVALPARCEIVVCPFLAHREAKPFPDPHAFSPSRWETARPSPYEYFPFGAGGHFCAGRNLALSLIREVLS<br>TLLSRFDFVLDGEQSIDWRIHIMLMPKGDPALIAHPVDERGDTPSPKWRGPITDLFHFAPGLS | | | |
| ApyA-L7Y-ApyOHIDS | pRSF-Duet | ORF | ATGGGCAGCAGCCATCACCATCATCACCACAGCCAGATGGCGACCAAACCAAAAAAGACCAAGGGCGTGAGCGTATCTATT<br>GAAGGGAAATTACCGAAAATGACGTTAGATATGCCTGTCGATGCAAAAAAAATCAAAGCTATTCAGAAATGTCTGGAGAACG<br>GTAAACTCACCATTACCATGTCTAAAGTAGATCTGGCAGGCGGTCGTATGGGGGAAGGTTATTACTACGATTGAGATTCAAG<br>GAGATATACCATGAAACTGTTGCTGGTTGATAACTTGGTA<br>ATGCCGGAGGAGGGCTCGCTGGCATATTTGGACGTGCATCCGCATCTGGGACTGCTGGCGCTGGCGGCCGTCGCAGAAGC<br>AGACGGTCACCGCGTGCAAATTTACGATCCGAAACGCCTGATTCGTGATGGCACACTGGCTTATGATGCCACCCTTTATGAA<br>CGCGCATCTCAAGCGATTCTGGCAGCGCGTCCTGATGGCGTCGGTTTCACTGCACTGGGCTGTTCCTTCCTGTTCGCGTTG<br>AATGTGGCGGCGTTACTGAAACGTCGCGAACCCGATCTGCCGGTGTTGCTGGGCGGTCCGCACGCGACCATGCTGCATCG<br>CCAGATTCTCGAACGTTTTCCGCAGTTTGATATCATCGTTCGCTATGAAGCCGATGAAATTCTGCCGGCCGTCCTCGATTGC<br>CTGCCACACCGCACCTTCGACGTGATCCCTGGCCTGAGTTGGCGTGCAACCGGTCGCGGCAGCCCGCTTCGGTTCACGGA<br>TGGCAAGCCGAAAGTTGAGGATCTGGACCTGCTGCCCATCGCTAGTTACGACCATTATCCGGTTGAAGAACTGGGACTGTC<br>GATGCTGCGTATCGAAGCCGGCCGCGGCTGCCCATTTGCGTGCACCTTTTGTTCCACGGCCGGTTTCTTTCAGCGTTCCTTC<br>CGCCTGAAGTCCGCTGAACGGCTGGTGCGTGAACTTGATATCTTACACCAACGTTACCGTGTCTCCGATTTCAAACTGGATC<br>ATGATATGTTTACTGTGAATCGTCGTAAAGTCATGGAATTCTGTGAAGCGGTTGCAGGTCGTGATTATCGCTGGCGTGCCTC<br>GGCCCGCATCGACTGTGTGGATGAAGCCCTGCTGAAGAAAATGGCGGATGCGGGTTGCGTTAACCTGTACTTTGGTGTTGA<br>AACAGGTTCAGAACGCATGCAGAAACTGTGTAAAAAGCGTCTGGATCTTCAACGTGTGGAGCCAATCCTGGCGGCCGCGGA<br>CAGCTTCGGCATTGAAACCACCGCCTCGTTCATCACCGGCTACCCGGAAGAAACGGGTGAAGATCAAGACGATACGCTCGA<br>TATGATTGGCCGCTGCGCCCGTCGTGCTAGTTGTCTGACCCAGCTGCATATGCTGGCGCCGGAACCGGGGACGCCGTTGTT<br>CGACGAACGCGGCGCGGAAATCGCGTATGACGGTTGTGGCGGTCCCTATAACACGCGTTTATTATCTAGCTCAGATGAACG<br>TGCGGTACTCGGTCACCCTGATATTTTCCAGACTTACTACCACTATCCGGCAGCTTTACCGCGTGCACGTTATATCTTTGCTG<br>TTGGGGCTGCCGATCTCCTGCGTCGTGTGGGTCCAGTAATTTTTGCGTATGCGCTGCGTGGATTCGGCGGCCGTCTGTCCA<br>CTCTGATTAGTGCGTGGCGCCAGTTCGCCGACGAGACCAGTCCGGGAGCCCCGGCGGACGCAGCGGGTCTGGAAGCGTTT<br>ATCGCTGCTCGTTTTGGCCGGGGCCACCACCTGCATAGTCTGTTCCGTTATGCGTTGCGCCTGTACGCCGCCCGTGCCCCT<br>GCAGATGTGATCGTCGAAGGTCCATATGAGCCAGATCGTCCATGCATGCTTGCTGAAAACGTCGGGTTGTTGGAAGACATTC<br>ACAATTGTGATGCCTTGGTTGATCGTATTCGCCGTTCAACCGAAGGCGCCCCGATGCTGGGCGATGCCGATGTCGGGGGCT<br>TGGGCACGTATATCGTGCAGGGTCACGGCGAAGCCGCCACAGGTTACTGGGTCGAATCCGGTATTGCGACTCTGCTGGATC<br>TGTTTCGTTCTCCTCGTAGTTGCCGGGCTGTGGCGCAATGGCTGGCAGACGCAACGCGTAACGATGGCATTGATGCCAGCA<br>TCTTTGAACCTCTGCTCCGCCGCGGTATTCTCGTGAGCGCAGGGTAA<br>GATTCAAGGAGATATACC<br>ATGCGTCGCACGCTCGAGCGCTGGTCAGCATTAGACTTGGCTGAGGGTCTGCAATTAGCGCACGCGGTGGCTGCGCTTCA<br>GGCGTTGGGTGTCCTGGATGCGATGCGTGAGCCAGTTACGGCAGAGACATTGTCTGCCTCTCACGATCTCGACCCGGAATT<br>GCTGCGTGGAATTCTTGAATACGCCGCATCCCGCACTAATTTGGTGCGTAAGACGGACCGTGGATTCGCGATTTCTGAGCAT<br>TATACCGCCCAAGCCCGTTTTCTCCTGAATTTATATGCCGGTGCTTACGCAGACAACGCCAGCCGTTTGGCCACATTATTAC<br>GTCGTCCCGCACTGGCTGGGAGCATGATCGATCTTGTGGCGCACGCTCGTGCATTCGATGCGAGTGAGGCTGGTGGTGGT<br>GCTCTTGCGTCTATTGTCACCCAATTGCGTCTCAACCATGTGTTGGATCTTGGTTGCGGCGCGGGCGCACTGTTACGTGAGT<br>TGGCAGCGGATGATCGTGGCTTCGTCGGCTGGGGTCTCGATCGCAATCCTGCGATGTGTAAGGCCGCACGTGTCCGTGCA<br>CGCCAGGCTGGATTGGCCGCGCGCGTCAAGGTTTTCCACGGAGACGGTTGGAATCCTGGTGCCAGCGTGCCACCGCGCGT<br>TTTAGCGCGCGTACGTAACGTCGTTGCTAGTCAATTCGTGAACGAAATGTTCCGCGGCGGTACGTCCCGCGTTGAGACCTG<br>GTTGCGCCGTATGCGTCGCCTCTTGCCCGGCCGTATGTTGGTTATTAGTGACTATTACGGTCGTCTCGGGCACGGCACCCA<br>CACCATGCGTCGCGAGACGCTTTTGCATGATTACGTGCAGTTGATCTCAGGTCAAGGTGTGCCGCCACCTGACTCCGATGC<br>GTGGTTGGCTATGTACCGTGCCGCAGGTTGCCGTCCGCTTCATGTGATCGAGGACCGCGCGGCGACTACTTGCTTCGTTCA<br>TTTAGTTGGTTTATGA<br>GTAGCGAAGCGGTCTAGCATAACCCCTTGGGGCCTCTAAACGGGTCTTGAGGGGTTTTTTGCCCTTAATACGACTCACTATA<br>GGGGAATTGTGAGCGGATAACAATTCCAAGGAGATATACA<br>ATGAACGCAGGCAAAATTCACGCGGTTATTGCGGCTGGTGCTAAAGACCCACGGTTGATTGAACATTGGCGTCGCCGTCCA<br>GATCTGCTGCGGCGCGCGGGTATCGAGCCCGATCATCTGGATCTTCGCGTCCTGGAGAATTTTGCGGGGCTGTCCCTTAAA<br>GTTCGCCACAATGGATTGCGCGGTGATTTGCCGTTAAGCTTTCGGTTGCTCTCTGTGGCGGGATTGGAAATTGAAGTTTTTG<br>CAGCGTACGCATCGGATTGCGCCGAATCAGGGCACGTCTTTGCCGCCACACCGCGTGAACGTGCTCGCGACCTCATGGCC<br>TTCCTTGACCGCTGGCTGGATCCGGCCCAGCGCAACCACGTGCTGTTATGGGACATCCTGCGCCATGAATACGCTCTCTCG<br>TGCCTGCGTCAGGAATCTGCGGCCCCGCTGCCGGCAGCGCGCCGTATTATTCGCCACCGTGATCGTGCAACTGGCGGTTG<br>CATTCCAATGCTGAATGGAAGTGTCGCCCAGCACGTTATGCATTGTAATCCAGAAGATCTGGCGTCGGCCTTACGCCAAAAA<br>GAACCGCCCCTGGATGCTGTTCCGGCCGCGGACACCTACTGTTGCTACTGGCGCAGTGCTGAGACGGGCGAGATCCGTTT<br>TCTCCAGGCAGATACCCTGGGTTATTTTGCAATTGAGGCAATCGATGGCCGCCGCTCTGCGGCGGACATCTGTGATCGCCT | MCS1<br><br>5'-NcoI--<br>His-ApyA-L7Y_<br>ApyD_ApyS--<br>3'-flank<br>GTAGCGAAG<br>CGGT--<br>Terminator<br><br>MCS2<br><br>5'-flank<br>AAGGAGATAT<br>ACA--<br>ApyH_ApyI_<br>ApyO—<br>3'-XhoI site<br><br>RBS |

| | | | |
|---|---|---|---|
| | | | GCTGGGCGGTCGTCGCCTCCTGCCCCAGTTTCTGCGCAGCCTGGACCAGCTCGCGGGCATTGGTATTGTACGGTTCGAGC<br>GTCCACTTGGAAGCGCGGCAGCATAA<br>GATTCAAGGAGATATACC<br>ATGAAAGCTTTTGATCGCGATGCATCAGAACCACGTATCGGCCTGGCCTATGGTCCGGGCATGGCGGAGTTTGCAGCCCGG<br>AACGCGCATCTCGTGGACTACATTGAAGTTCCGTTTGAACAACTGCGTTTCAGCCCAGCAGTAGCCGAGCTGCAGCAGACG<br>ATTCCGTTCGTGCTGCATTGCGCGAGCTTGAGCGTGGCTGGCTTTGTGCCGCCGGATGACTCGACTGTAGATGCAATTGAA<br>CGCACCGCAGTGCAAACTGGTACCCCGTGGATCGGCGAGCACCTGGCGTATATCTCGGCGGACCCCGTTGGTGAAGCATT<br>AGGTGGTACCGGGGAGCCGACTTCGCTGTCATACACCCTCTGTCCGCAGTTATCCGACGAGACCGTCCGTCGCGTGGTGG<br>ATAATTTAGCCGCACTGCGTCCTCACTTTCCGGTGCCCTTGATCGTAGAAAACTCGCCGCAATATTTTCCAATCCCCGGCAG<br>CACTATGGGCATGACCGATTTTATCCGTGCCATTACGGATCGCTGTGATGTTGGCTTATTACTTGATTTATCTCACTTTCTGAT<br>CACTGCCCACAATACGGGCGCAGAAGTTCACCGCGAACTGGCGCGCCTGCCACTGGAACGCGTGGTGGAGGTACACTTGT<br>CCGGCATGTCTGTGCAAAGCGGTACCGCCTGGGATGATCATTCATTACCGGCCTCACCTATTTTATTTGAACTGTTGGAACG<br>GTTGCTGGACGTGGCTCGCCCGCGCGCCCTGACATTTGAATACAATTGGAGTCCCTACTTTCCACTGAGCGTGTTGACTACC<br>CACATCGAACGCGCCCGTCAGCTGATGGGGCTGGCTTAA<br>GATTCAAGGAGATATACC<br>ATGAACGGTCTCCCGCCTTCTCTGCCTCGTGTGGACTCAACAGAAAGCCTGTTCGCGGAGCCTTTAGCATTCTTAGCGCAAG<br>CCCGTTCTCGTCACGGAGACGTGTTCGTTATGCGCGAGCACGGCCCCATCTTTTCCCGCGCATCGGATTGTTCGGGTGTCA<br>TTGCTGTCTTTGGAGAGCACCGTTTACGTCAAATCCTTACGGACATCGACAACTTCGCGTTACCGATGAGTGCCGCCGCCAA<br>GATGGCATTACCTAAGAACCTCGTCAATTTAAACCGTGGCCTGCATAGTATGCGCGAGCCGGAGCATGGTCGTCACAAGCG<br>TTTGCTTACCGGTACAATTAACCGCGAGCTCTTTGACGCCCACCGTTTCGAAATTCGCGCAGCGCTCAACCGCTTTTGTGAA<br>ATGCTCAAGGTAGATCGCCGTATCAGTGTGGTGAGCCGTATGCGCGAGCTGACGGTCGAGATGGCATCTCATATCTTTCTG<br>GGTGCACAATGTCAGGAAGACGATGAGTTGGCGTTTCTTCTTTCAGCCTATTTCACCCTGCGCCGCGAGGCTAGCTCATTAA<br>ACGCTCGTGACCCGTTGCTTTATCGTGACGAGCTGATCGGTGTGGGACAACAGCTGGACCGCACATTACGCGAACGTATCC<br>GCCGCTATCGTAAACGCCCCGTCGACGCCCGCGCCGGTCTGTTACAGCGTCTTGCTACGGCAGGTCCGCCGGGCTCGCCT<br>GCGTTGTCGGAAGATGAAATTGTGGGTCACGCCAACGTAATGTTCGTGTCTTCCACGGAGCCGGTTGCCATGTCTCTTACGT<br>GGTTGTTGCTGGTCTTAAGTCAACTGCCTGACTTACGTCGTGCGCTGCGTGCGGAGATCGCGGACCGTGCGTCTATGCCGG<br>CCAGTACTAATGGCGCGTCGTGGTTAGAGAACGTTGTCAACGAGACGTTGCGCTTATTGACCCCAAATGCCCTGATGGTCC<br>GCGCCACAACACGCGCAGTGTCACTGCAAGGTGTTGCACTCCCGGCCCGCTGTGAGATTGTGGTATGTCCATTTCTGGCTC<br>ACCGCGAGGCCAAACCATTTCCCGACCCCCACGCATTCTCACCATCGCGCTGGGAGACCGCGCGTCCCAGTCCTTATGAGT<br>ACTTTCCTTTCGGGGCGGGCGGACACTTCTGTGCGGGGCGTAATTTGGCCCTCTCACTTATTCGCGAAGTGTTGTCGACGC<br>TGTTAAGCCGTTTCGATTTCGTTCTTGACGGTGAGCAAAGCATCGACTGGCGCATTCATATTATGCTGATGCCCAAGGGAGA<br>CCCGGCTTTGATTGCTCACCCTGTGGATGAACGTGGCGATACCCCGAGTCCAAAGTGGCGTGGGCCCATTACCGATTTATTT<br>CACTTCGCCCCTGGGCTTTCATGA |
| Translated sequence | ApyA-L7Y: MGSSHHHHHHSQMATKPKKTKGVSVSIEGKLPKMTLDMPVDAKKIKAIQKCLENGKLTITMSKVDLAGGRMGEGYYYD<br><br>ApyD:<br>MKLLLVDNLVMPEEGSLAYLDVHPHLGLLALAAVAEADGHRVQIYDPKRLIRDGTLAYDATLYERASQAILAARPDGVGFTALGCSFLFALNVAALLKRREPDLPVLLGGPHATMLHR<br>QILERFPQFDIIVRYEADEILPAVLDCLPHRTFDVIPGLSWRATGRGSPLRFTDGKPKVEDLDLLPIASYDHYPVEELGLSMLRIEAGRGCPFACTFCSTAGFFQRSFRLKSAERLVREL<br>DILHQRYRVSDFKLDHDMFTVNRRKVMEFCEAVAGRDYRWRASARIDCVDEALLKKMADAGCVNLYFGVETGSERMQKLCKKRLDLQRVEPILAAADSFGIETTASFITGYPEETG<br>EDQDDTLDMIGRCARRASCLTQLHMLAPEPGTPLFDERGAEIAYDGCGGPYNTRLLSSSDERAVLGHPDIFQTYYHYPAALPRARYIFAVGAADLLRRVGPVIFAYALRGFGGRLST<br>LISAWRQFADETSPGAPADAAGLEAFIAARFGRGHHLHSLFRYALRLYAARAPADVIVEGPYEPDRPCMLAENVGLLEDIHNCDALVDRIRRSTEGAPMLGDADVGGLGTYIVQGHG<br>EAATGYWVESGIATLLDLFRSPRSCRAVAQWLADATRNDGIDASIFEPLLRRGILVSAG<br><br>ApyS:<br>MRRTLERWSALDLAEGLQLAHAVAALQALGVLDAMREPVTAETLSASHDLDPELLRGILEYAASRTNLVRKTDRGFAISEHYTAQARFLLNLYAGAYADNASRLATLLRRPALAGSMI<br>DLVAHARAFDASEAGGGALASIVTQLRLNHVLDLGCGAGALLRELAADDRGFVGWGLDRNPAMCKAARVRARQAGLAARVKVFHGDGWNPGASVPPRVLARVRNVVASQFVNE<br>MFRGGTSRVETWLRRMRRLLPGRMLVISDYYGRLGHGTHTMRRETLLHDYVQLISGQGVPPPDSDAWLAMYRAAGCRPLHVIEDRAATTCFVHLVGL<br><br>ApyH:<br>MNAGKIHAVIAAGAKDPRLIEHWRRRPDLLRRAGIEPDHLDLRVLENFAGLSLKVRHNGLRGDLPLSFRLLSVAGLEIEVFAAYASDCAESGHVFAATPRERARDLMAFLDRWLDPA<br>QRNHVLLWDILRHEYALSCLRQESAAPLPAARRIIRHRDRATGGCIPMLNGSVAQHVMHCNPEDLASALRQKEPPLDAVPAADTYCCYWRSAETGEIRFLQADTLGYFAIEAIDGRR<br>SAADICDRLLGGRRLLPQFLRSLDQLAGIGIVRFERPLGSAAA<br><br>ApyI:<br>MKAFDRDASEPRIGLAYGPGMAEFAARNAHLVDYIEVPFEQLRFSPAVAELQQTIPFVLHCASLSVAGFVPPDDSTVDAIERTAVQTGTPWIGEHLAYISADPVGEALGGTGEPTSLS<br>YTLCPQLSDETVRRVVDNLAALRPHFPVPLIVENSPQYFPIPGSTMGMTDFIRAITDRCDVGLLLDLSHFLITAHNTGAEVHRELARLPLERVVEVHLSGMSVQSGTAWDDHSLPASPI<br>LFELLERLLDVARPRALTFEYNWSPYFPLSVLTTHIERARQLMGLA<br><br>ApyO:<br>MNGLPPSLPRVDSTESLFAEPLAFLAQARSRHGDVFVMREHGPIFSRASDCSGVIAVFGEHRLRQILTDIDNFALPMSAAAKMALPKNLVNLNRGLHSMREPEHGRHKRLLTGTINR<br>ELFDAHRFEIRAALNRFCEMLKVDRRISVVSRMRELTVEMASHIFLGAQCQEDDELAFLLSAYFTLRREASSLNARDPLLYRDELIGVGQQLDRTLRERIRRYRKRPVDARAGLLQRL<br>ATAGPPGSPALSEDEIVGHANVMFVSSTEPVAMSLTWLLLVLSQLPDLRRALRAEIADRASMPASTNGASWLENVVNETLRLLTPNALMVRATTRAVSLQGVALPARCEIVVCPFLA<br>HREAKPFPDPHAFSPSRWETARPSPYEYFPFGAGGHFCAGRNLALSLIREVLSTLLSRFDFVLDGEQSIDWRIHIMLMPKGDPALIAHPVDERGDTPSPKWRGPITDLFHFAPGLS | | |
| ApyA-L7Y-ApyOHIDS | pSCrha2 | ORF | ATGAACGGTCTACCTCCGTCGCTTCCGCGGGTCGACAGCACCGAGTCGCTTTTTGCGGAGCCGCTCGCGTTTCTCGCACAA<br>GCGCGCTCGCGGCATGGTGATGTCTTCGTCATGCGGGAGCATGGTCCGATTTTTCCCGCGCGAGCGATTGCAGCGGCGT<br>GATAGCGGTATTCGGCGAGCATCGGCTTCGACAGATATTGACCGATATAGACAATTTCGCGTTGCCGATGTCCGCGGCGGC<br>GAAAATGGCATTGCCGAAAAACCTCGTCAACCTCAATCGCGGCTTGCACAGCATGCGCGAGCCGGAGCATGGCCGGCACAA<br>GCGCCTCTTGACGGGAACGATCAATCGCGAGTTGTTCGACGCGCACCGCTTCGAAATACGAGCGGCTTTGAACCGCTTTTG<br>CGAAATGCTGAAAGTGGACCGCCGGATTTCAGTGGTCAGCCGGATGCGCGAGCTGACCGTGGAGATGGCGTCTCATATCTT<br>TCTTGGGGCGCAGTGCCAAGAGGATGACGAACTGGCGTTTCTTTTAAGCGCCTATTTTACTTTGCGACGCGAAGCGTCTTCG<br>CTGAACGCGCGCGATCCCCTGCTGTACCGGGACGAACTGATCGGCGTCGGGCAGCAACTCGACCGCACATTGCGGGAGCG<br>GATCAGGCGGTATCGAAAGCGTCCCGTCGACGCTCGTGCGGGGCTGCTGCAACGTCTGGCAACGGCCGGACCGCCGGGC<br>TCGCCCGCGCTTTCCGAGGACGAGATAGTCGGCCATGCCAACGTGATGTTCGTGTCGAGCACCGAGCCCGTCGCGATGTC<br>GCTGACGTGGCTGCTGCTCGTTCTGTCGCAGCTGCCCGATCTGAGGCGCGCGCTTCGCGCGGAAATTGCCGATCGCGCGT<br>CGATGCCGGCATCGACGAACGGCGCGTCGTGGCTCGAGAACGTCGTCAACGAGACGCTCCGGCTCCTGACGCCCAACGC<br>GCTGATGGTTCGCGCGACGACGCGCGCGGTCTCGCTGCAAGGCGTCGCGCTTCCGGCGCGTTGCGAGATCGTCGTGTGTC<br>CGTTCCTCGCGCATCGCGAAGCCAAGCCGTTTCCGGACCCGCATGCATTCTCGCCGTCCCGGTGGGAGACGGCCAGGCCG<br>TCTCCGTACGAGTATTTTCCGTTCGGCGCCGGCGGTCATTTTTGCGCGGGGCGAAATCTCGCGCTCTCGCTGATTCGCGAA<br>GTACTGTCAACGCTGCTATCGCGCTTCGATTTCGTTTTGGATGGCGAACAGTCTATCGACTGGCGCATTCATATCATGTTGAT<br>GCCGAAAGGCGACCCGGCACTCATCGCGCATCCCGTCGATGAGCGCGGTGACACGCCGTCGCCGAAATGGCGCGGGCCG<br>ATAACCGATCTATTCCATTTCGCGCCGGGGCTTTCTTGA<br>GGTACCCATTTACCAGGGCGCATCCTTTAATCAGAAGCGAATCGGCATGCATGCCCATCGGGATTTTTTCCAGTCATTGATC<br>CGCATGGTTCTTTTCACGGGTTATTCCGCGCTCACACGCTAAGCGGCGCGTCCTGCCTACAACAAAGGAACAATC<br>ATGCATCACCATCACCATCACATGGCTACGAAACCCAAAAAAACAAAGGGCGTTTCTGTTTCCATCGAAGGGAAGCTGCCCA<br>AGATGACCCTGGATATGCCGGTGGACGCGAAAAAGATCAAGGCCATCCAGAAATGCCTGGAAAACGGCAAGCTGACCATCA<br>CGATGAGCAAGGTGGATCTTGCCGGCGGCCGCATGGGCGAGGGCTATTACTACGACTGA<br>GGATCCTGTCCGGCGGCGCTCTGGTTTTCGCCGGCGGCGTCGCGATCTCGCCGCCTCGATGCTCGGGCCTCGGAGGGAA<br>CGTGCCGCCGAACGCGTGCGTTTGGCGCGCGGTCGCGCGATAACGGGCGATGAATTCCGGGCTTCGGCAGTTCGACCTTA<br>AGGGCGCGGGAAGCCT<br>ATGAAAGCCTTCGATCGCGATGCATCGGAGCCTCGGATCGGTCTCGCGTATGGGCCCGGGATGGCGGAATTCGCCGCACG<br>GAACGCGCACCTCGTCGATTACATCGAAGTGCCGTTCGAGCAATTGCGATTCTCGCCGGCGGTCGCCGAGCTTCAGCAAAC<br>CATTCCGTTCGTTCTGCACTGCGCCAGCTTGTCCGTCGCGGGATTCGTGCCGCCCGACGATTCGACGGTCGACGCAATCGA<br>GCGCACGGCCGTGCAGACCGGCACGCCATGGATAGGCGAACATCTCGCGTACATTTCGGCCGATCCCGTTGGCGAAGCGC<br>TGGGCGGGACCGGTGAGCCCACCTCGTTGTCGTACACGTTGTGCCCTCAGTTGAGCGACGAGACGGTACGCCGCGTCGTC<br>GACAATCTCGCCGCGCTTCGTCCGCATTTTCCGGTTCCACTGATCGTCGAAAACTCCCCGCAGTACTTTCCGATACCGGGCA<br>GCACGATGGGAATGACGGACTTCATTCGCGCGATCACGGACCGATGCGACGTCGGTTTGCTCCTGGACCTGAGTCACTTCC<br>TGATCACGGCGCACAACACGGGCGCAGAGGTGCATCGGGAACTCGCCCGGCTTCCGCTGGAGCGCGTGGTCGAAGTTCAT<br>CTGTCCGGAATGAGCGTGCAGTCCGGCACGGCCTGGGACGACCATTCGCTGCCGGCATCGCCGATCCTGTTCGAACTGCT<br>CGAGCGCCTGCTCGACGTGGCGCGGCCTCGCGCGCTGACCTTCGAATACAACTGGTCGCCATACTTCCCGCTGTCGGTCC<br>TGACGACGCATATCGAGCGCGCACGCCAGTTGATGGGGCTCGCATGAACGCAGGGAAAATACACGCCGTGATCGCCGCCG<br>GCGCCAAGGACCCGCGGCTCATCGAGCATTGGCGGCGGCGTCCGGACCTGCTGCGCCGCGCAGGAATCGAGCCGGATCA<br>TCTCGATTTGCGCGTATTGGAGAATTTCGCCGGGTTGAGCCTGAAGGTGCGTCACAACGGCCTGCGCGGGGATCTGCCGCT |

| | | | |
|---|---|---|---|
| | | | GAGTTTCAGGTTGTTGAGCGTGGCCGGCCTCGAGATCGAGGTCTTCGCCGCGTATGCGTCGGACTGCGCGGAAAGCGGCC<br>ATGTATTCGCCGCCACGCCGCGGGAACGCGCGCGCGATCTGATGGCGTTCCTCGACAGATGGCTGGACCCCGCGCAACGA<br>AATCACGTTCTGCTGTGGGACATCCTTCGTCATGAATATGCGCTTTCGTGCCTGCGGCAAGAAAGCGCCGCGCCGCTACCC<br>GCCGCTCGCCGCATCATCCGCCATCGGGATCGCGCGACAGGCGGCTGCATTCCGATGCTCAACGGCTCCGTCGCTCAGCA<br>CGTGATGCATTGCAATCCGGAAGACCTGGCGTCGGCGCTGCGTCAGAAGGAGCCGCCGCTCGATGCGGTTCCCGCCGCCG<br>ACACTTACTGCTGCTATTGGCGCTCCGCCGAAACGGGCGAAATCCGGTTTCTGCAGGCCGATACGCTCGGCTATTTCGCGA<br>TCGAAGCGATCGACGGCCGGCGATCCGCCGCCGACATCTGCGATCGGCTGCTCGGCGGCCGCCGTCTGCTGCCGCAGTT<br>CCTCCGGTCGCTCGACCAACTGGCCGGAATCGGGATCGTGCGGTTCGAGAGGCCGCTCGGAAGCGCTGCGGCATGAAACT<br>GCTGCTCGTCGACAACCTCGTCATGCCGGAAGAGGGCTCGCTCGCGTACCTCGACGTGCATCCGCATCTCGGGCTGCTCG<br>CGCTGGCCGCGGTCGCCGAAGCCGACGGCCATCGCGTGCAGATCTACGATCCGAAACGACTGATCAGGGACGGCACGCT<br>GGCTTACGATGCGACGCTTTACGAGCGGGCGAGCCAAGCGATTCTCGCGGCGCGTCCGGACGGGGTCGGCTTCACGGCG<br>CTCGGCTGCAGCTTTCTGTTTGCGCTCAATGTCGCGGCGCTGTTGAAGCGGCGCGAGCCCGACCTGCCCGTCTTGCTCGG<br>CGGGCCGCACGCGACGATGCTGCACCGGCAGATCCTCGAGCGGTTCCCGCAGTTCGACATCATCGTCCGGTACGAGGCGG<br>ATGAAATCCTGCCCGCCGTGCTCGACTGCCTCCCGCACCGGACATTCGACGTGATTCCCGGCTTGAGCTGGCGCGCGACG<br>GGCCGCGGCTCGCCGTTGCGCTTCACCGACGGCAAGCCCAAGGTCGAGGATCTGGACCTGCTGCCGATCGCCTCGTACGA<br>CCATTATCCGGTTGAGGAACTGGGGCTCTCGATGCTGCGCATCGAGGCCGGCCGAGGGTGTCCGTTCGCATGCACATTCTG<br>CTCGACGGCCGGTTTCTTTCAACGCAGCTTCCGCCTCAAATCGGCCGAGCGCCTGGTTCGCGAACTGGATATCCTTCACCA<br>GCGTTATCGCGTGTCGGACTTCAAGCTCGACCACGACATGTTCACCGTGAATCGCCGCAAGGTAATGGAGTTTTGCGAGGC<br>GGTCGCCGGGCGCGACTATCGGTGGCGCGCGTCGGCGCGGATCGACTGCGTCGACGAAGCATTGTTGAAGAAGATGGCC<br>GACGCGGGCTGCGTGAATCTTTATTTCGGCGTGGAGACGGGCTCCGAGCGCATGCAAAAGCTCTGCAAGAAGCGGCTGGA<br>TCTGCAACGGGTCGAGCCCATTCTGGCGGCGGCGGATTCATTCGGAATCGAGACGACCGCGTCGTTCATCACCGGCTATCC<br>CGAGGAAACCGGGGAGGATCAGGACGACACGCTGGACATGATCGGGCGCTGCGCTCGGCGAGCGTCTTGCCTCACGCAA<br>CTGCACATGCTCGCGCCGGAGCCCGGCACACCGCTGTTCGACGAACGCGGCGCGGAAATCGCCTACGACGGCTGTGGCG<br>GGCCTTACAACACGCGCCTGCTGAGTTCGTCGGACGAGCGTGCGGTGCTCGGCCATCCCGACATATTCCAAACCTACTATC<br>ACTACCCCGCCGCGCTGCCGCGTGCGCGATACATCTTTGCCGTCGGGGCGGCGGATCTGCTGCGCCGGGTCGGGCCGGT<br>CATTTTCGCGTATGCGTTGCGGGGGTTCGGAGGGCGGCTGAGCACGCTGATATCGGCGTGGCGACAATTCGCCGACGAAA<br>CGTCGCCCGGCGCGCCCGCGGACGCGGCGGGGCTCGAGGCCTTCATTGCCGCGCGGTTCGGCCGCGGCCATCACCTGC<br>ATTCTCTTTTTCGGTACGCGTTGCGGCTCTATGCGGCCCGCGCTCCGGCGGACGTAATCGTCGAGGGGCCCTATGAGCCG<br>GACCGGCCTTGCATGCTGGCCGAAAACGTCGGTCTGCTGGAGGACATTCACAATTGCGACGCATTGGTCGACCGGATCAGG<br>CGATCGACCGAAGGCGCGCCGATGCTCGGCGACGCGGATGTCGGCGGGCTCGGAACATATATCGTTCAGGGGCACGGCG<br>AGGCCGCCACGGGCTATTGGGTGGAGTCCGGCATCGCTACGCTTCTCGATCTGTTCAGGTCGCCCCGTTCGTGCCGCGCC<br>GTTGCGCAATGGCTGGCGGACGCCACCCGCAATGACGGCATAGACGCTTCGATCTTCGAGCCGCTGCTGCGCAGAGGCAT<br>CCTGGTGTCGGCCGGTTAAATCCATTTCGTCCATGGTGCGTCGCACACTCGAACGTTGGTCCGCGCTGGATCTCGCGGAAG<br>GGCTTCAACTGGCGCATGCCGTCGCCGCGCTGCAGGCGCTCGGCGTTCTGGACGCGATGAGGGAGCCGGTCACGGCGGA<br>AACGTTGAGCGCCAGCCATGATCTCGATCCCGAGTTGCTGCGCGGCATCCTCGAATATGCGGCGTCGAGAACGAATCTGGT<br>TCGCAAGACGGACCGCGGATTCGCGATTTCCGAGCATTACACCGCGCAAGCGCGCTTTCTGTTGAACCTGTACGCCGGCGC<br>TTATGCCGATAACGCTTCACGGCTGGCGACGCTGCTCCGGCGTCCGGCATTGGCGGGCTCCATGATCGACCTCGTCGCGC<br>ACGCGCGCGCCTTCGACGCGAGCGAGGCCGGCGGTGGAGCGCTGGCGTCGATCGTCACGCAATTGCGTCTCAATCATGTG<br>CTGGATCTGGGATGCGGCGCCGGCGCGCTGCTGCGCGAGTTGGCGGCGGACGATCGCGGATTCGTCGGCTGGGGACTGG<br>ATCGCAATCCCGCGATGTGCAAGGCGGCGCGGGTTCGAGCGCGGCAAGCGGGGCTCGCCGCGCGCGTGAAGGTGTTCCA<br>CGGCGACGGCTGGAATCCCGGGGCGTCGGTCCCGCCGCGCGTGCTGGCGCGCGTGCGCAACGTCGTCGCGAGCCAGTTC<br>GTCAACGAGATGTTTCGCGGCGGCACATCGCGAGTGGAAACGTGGCTTCGGCGCATGCGGCGCTTGCTGCCCGGCCGCAT<br>GCTCGTGATATCGGACTATTACGGCCGCCTCGGCCACGGCACTCACACGATGCGCCGCGAGACGCTGCTGCATGACTATGT<br>GCAGCTCATATCGGGGCAGGGCGTGCCGCCGCCCGATTCCGACGCGTGGTTGGCCATGTATCGAGCAGCCGGTTGCCGG<br>CCTCTGCACGTGATCGAAGACCGGGCGGCCACGACATGCTTCGTCCATCTGGTGGGGCTATAG |
| Translated sequence | Same as above (pRSF construct) | | |

**Table S2. NMR assignments of ApyO-modified, GluC-digested ApyA-Y6W**

| number | AA | NH/C=O | αH/C | βH/C | γH/C | δH/C and others |
|---|---|---|---|---|---|---|
| 1 | G5 | <br>173.0 | 3.29<br>46.1 | | | |
| 2 | W6 | 8.12 (br)<br>172.9 | 4.80<br>56.1 | 3.14, 3.05<br>30.4 | | Cγ: 114.7<br>δ1: 7.01 (s), 132.2<br>Cε2: 140.2<br>Cδ2: 131.8<br>ε3: 7.45 (d), 121.6<br>ζ3: 7.04 (t), 122.2<br>η2: 7.08 (t), 124.5<br>ζ2: 7.14 (d), 115.3<br>ε1: N- (linker) |
| 3 | L7 | 8.29<br>174.8 | 4.64<br>54.3 | 1.56, 1.44<br>45.5 | 1.56<br>27.6 | 0.86, 25.6<br>0.89, 25.9 |
| 4 | Y8 | 8.85<br>172.8 | 4.39<br>57.3 | 2.97, 2.80<br>37.7 | | Cγ: 133.7<br>δ1: 7.045 (d), 131.9<br>ε1: 6.95 (d), 119.5<br>δ2: 6.87 (s), 131.3<br>Cε2 (linker): 129.8<br>Cζ-OH: 153.3 |
| 5 | D9 | 7.55<br>175.1 | 4.09<br>52.1 | 2.41<br>42.9 | | Cγ=O: 176.1 |

Reference: DMSO with $^{1}$H: 2.48 ppm, $^{13}$C: 42.8 ppm

#### Table S3. Primers used for KLENOW fragment extension.

| Primer | Sequence |
|---|---|
| FP_T7p-RBS | GGCGTAATACGACTCACTATAGGGTTAACTTTAACAAGGAGAAAAAC |
| RP_R1Y | CGAAGCTCAATCGTACAGATAACCTTCCCCCATGTACATGTTTTTCTCCTTGTTAAAGTT |
| RP_R1W | CGAAGCTCAATCGTACAGATAACCTTCCCCCATCCACATGTTTTTCTCCTTGTTAAAGTT |
| RP_R1V | CGAAGCTCAATCGTACAGATAACCTTCCCCCATGACCATGTTTTTCTCCTTGTTAAAGTT |
| RP_R1T | CGAAGCTCAATCGTACAGATAACCTTCCCCCATTGTCATGTTTTTCTCCTTGTTAAAGTT |
| RP_R1S | CGAAGCTCAATCGTACAGATAACCTTCCCCCATGCTCATGTTTTTCTCCTTGTTAAAGTT |
| RP_R1Q | CGAAGCTCAATCGTACAGATAACCTTCCCCCATTTGCATGTTTTTCTCCTTGTTAAAGTT |
| RP_R1P | CGAAGCTCAATCGTACAGATAACCTTCCCCCATGGGCATGTTTTTCTCCTTGTTAAAGTT |
| RP_R1N | CGAAGCTCAATCGTACAGATAACCTTCCCCCATGTTCATGTTTTTCTCCTTGTTAAAGTT |
| RP_R1M | CGAAGCTCAATCGTACAGATAACCTTCCCCCATCATCATGTTTTTCTCCTTGTTAAAGTT |
| RP_R1L | CGAAGCTCAATCGTACAGATAACCTTCCCCCATTAACATGTTTTTCTCCTTGTTAAAGTT |
| RP_R1K | CGAAGCTCAATCGTACAGATAACCTTCCCCCATTTTCATGTTTTTCTCCTTGTTAAAGTT |
| RP_R1I | CGAAGCTCAATCGTACAGATAACCTTCCCCCATGATCATGTTTTTCTCCTTGTTAAAGTT |
| RP_R1H | CGAAGCTCAATCGTACAGATAACCTTCCCCCATATGCATGTTTTTCTCCTTGTTAAAGTT |
| RP_R1G | CGAAGCTCAATCGTACAGATAACCTTCCCCCATGCCCATGTTTTTCTCCTTGTTAAAGTT |
| RP_R1F | CGAAGCTCAATCGTACAGATAACCTTCCCCCATGAACATGTTTTTCTCCTTGTTAAAGTT |
| RP_R1E | CGAAGCTCAATCGTACAGATAACCTTCCCCCATTTCCATGTTTTTCTCCTTGTTAAAGTT |
| RP_R1D | CGAAGCTCAATCGTACAGATAACCTTCCCCCATATCCATGTTTTTCTCCTTGTTAAAGTT |
| RP_R1C | CGAAGCTCAATCGTACAGATAACCTTCCCCCATGCACATGTTTTTCTCCTTGTTAAAGTT |
| RP_R1A | CGAAGCTCAATCGTACAGATAACCTTCCCCCATGGCCATGTTTTTCTCCTTGTTAAAGTT |
| RP_L7Y | CGAAGCTCAATCGTAATAATAACCTTCCCCCATACGCATGTTTTTCTCCTTGTTAAAGTT |
| RP_L7W | CGAAGCTCAATCGTACCAATAACCTTCCCCCATACGCATGTTTTTCTCCTTGTTAAAGTT |
| RP_L7V | CGAAGCTCAATCGTACACATAACCTTCCCCCATACGCATGTTTTTCTCCTTGTTAAAGTT |
| RP_L7T | CGAAGCTCAATCGTATGTATAACCTTCCCCCATACGCATGTTTTTCTCCTTGTTAAAGTT |
| RP_L7S | CGAAGCTCAATCGTAAGAATAACCTTCCCCCATACGCATGTTTTTCTCCTTGTTAAAGTT |
| RP_L7R | CGAAGCTCAATCGTAGCGATAACCTTCCCCCATACGCATGTTTTTCTCCTTGTTAAAGTT |
| RP_L7Q | CGAAGCTCAATCGTACTGATAACCTTCCCCCATACGCATGTTTTTCTCCTTGTTAAAGTT |
| RP_L7P | CGAAGCTCAATCGTATGGATAACCTTCCCCCATACGCATGTTTTTCTCCTTGTTAAAGTT |
| RP_L7N | CGAAGCTCAATCGTAATTATAACCTTCCCCCATACGCATGTTTTTCTCCTTGTTAAAGTT |
| RP_L7M | CGAAGCTCAATCGTACATATAACCTTCCCCCATACGCATGTTTTTCTCCTTGTTAAAGTT |
| RP_L7K | CGAAGCTCAATCGTACTTATAACCTTCCCCCATACGCATGTTTTTCTCCTTGTTAAAGTT |
| RP_L7I | CGAAGCTCAATCGTAAATATAACCTTCCCCCATACGCATGTTTTTCTCCTTGTTAAAGTT |
| RP_L7H | CGAAGCTCAATCGTAATGATAACCTTCCCCCATACGCATGTTTTTCTCCTTGTTAAAGTT |
| RP_L7G | CGAAGCTCAATCGTAGCCATAACCTTCCCCCATACGCATGTTTTTCTCCTTGTTAAAGTT |
| RP_L7F | CGAAGCTCAATCGTAGAAATAACCTTCCCCCATACGCATGTTTTTCTCCTTGTTAAAGTT |
| RP_L7E | CGAAGCTCAATCGTATTCATAACCTTCCCCCATACGCATGTTTTTCTCCTTGTTAAAGTT |
| RP_L7D | CGAAGCTCAATCGTAATCATAACCTTCCCCCATACGCATGTTTTTCTCCTTGTTAAAGTT |
| RP_L7C | CGAAGCTCAATCGTAGCAATAACCTTCCCCCATACGCATGTTTTTCTCCTTGTTAAAGTT |
| RP_L7A | CGAAGCTCAATCGTAAGCATAACCTTCCCCCATACGCATGTTTTTCTCCTTGTTAAAGTT |
| RP_E4K | CGAAGCTCAATCGTACAGATAACCTTTCCCCATACGCATGTTTTTCTCCTTGTTAAAGTT |
| RP_E4A | CGAAGCTCAATCGTACAGATAACCTGCGCCCATACGCATGTTTTTCTCCTTGTTAAAGTT |
| RP_Y6/8H | CGAAGCTCAATCGTGCAGATGACCTTCCCCCATACGCATGTTTTTCTCCTTGTTAAAGTT |
| RP_Y6H | CGAAGCTCAATCGTACAGATGACCTTCCCCCATACGCATGTTTTTCTCCTTGTTAAAGTT |
| RP_Y8H | CGAAGCTCAATCGTGCAGATAACCTTCCCCCATACGCATGTTTTTCTCCTTGTTAAAGTT |
| RP_8-mer | CGAAGCTCAATCGTACAGATAACCTTCACCCATGTTTTTCTCCTTGTTAAAGTT |

##### Table S4. NMR assignments of ApyO-modified, GluC-digested ApyA-L7Y Isomer–1

| number | AA | NH/C=O | αH/C | βH/C | Others |
|---|---|---|---|---|---|
| 1 | G5 | <br>166.7 | 4.08<br>41.4 | | |
| 2 | Y6 | 8.43 (br)<br>169.8 | 5.06<br>54.2 | 3.19, 3.16<br>37.0 | Cγ: 130.2<br>δ1: 7.02 (d), 130.7<br>ε1: 6.90 (d), 116.0<br>δ2: 6.76 (s), 133.8<br>Cε2: (linker): 128.2<br>Cζ-OH: 154.9 |
| 3 | Y7 | 8.74<br>172.4 | 4.92<br>55.4 | 2.91<br>38.2 | Cγ: 131.4<br>6.87 (br), 130.6<br>6.79 (d), 115.8<br>Cζ-OH: 157.5 |
| 4 | Y8 | 8.68 (br)<br>171.2 | 4.33<br>N/A | 2.58 (br)<br>N/A | Cγ: 134.3<br>δ1: 7.09 (d), 130.2<br>ε1: 6.82 (d), 116.0<br>δ2: 6.84 (s), 131.0<br>Cε2: linker: 128.4<br>Cζ-OH: 154.1 |
| 5 | D9 | 8.17<br>175.5 | 4.55<br>51.0 | 2.96, 2.87<br>37.0 | Cγ=O: 175.5 |

##### Table S5. NMR assignments of ApyO-modified, GluC-digested ApyA-L7Y Isomer–2

| number | AA | NH/C=O | αH/C | βH/C | Others |
|---|---|---|---|---|---|
| 1 | G5 | <br>166.0 | 3.66<br>40.5 | | |
| 2 | Y6 | 7.42<br>170.2 | 4.67<br>52.5 | 2.80, 2.70<br>37.2 | Cγ: 127.2<br>δ1: 6.53 (dd), 124.3<br>ε1: 6.79 (d), 116.2<br>δ2: 5.72 (d, 1.5Hz), 115.9<br>Cε2: 147.8 (linker)<br>Cζ-OH: 144.0 |
| 3 | Y7 | 8.09<br>171.4 | 4.45<br>53.7 | 2.90, 2.53<br>38.4 | Cγ: 128.1<br>6.91, 131.0<br>6.64, 115.5<br>C-OH: 154.5 |
| 4 | Y8 | 8.49<br>172.1 | 4.74<br>57.7 | 3.32, 2.68<br>37.7 | Cγ: 134.3<br>δ1: 7.39 (dd), 130.7<br>δ2: 7.18 (dd), 132.4<br>ε1: 6.99 (dd), 122.5<br>ε2: 6.80 (dd), 121.2<br>Cζ-O-: 153.3 (linker) |
| 5 | D9 | 8.29<br>175.1 | 4.58<br>50.5 | 2.86<br>37.0 | Cγ=O: 175.7 |

**Table S6. NMR assignments of ApyO-modified, GluC-digested ApyA-L7W Isomer–1**

| number | AA | NH/C=O | αH/C | βH/C | Others |
|---|---|---|---|---|---|
| 1 | G5 | | 3.43<br>41.7 | | |
| 2 | Y6 | 7.90 | 4.62<br>53.6 | 3.09, 2.89<br>37.4 | Cγ: 126.4<br>δ1: 6.65, 129.6<br>ε1: 6.51, 121.5<br>δ2: 6.84, 130.7<br>Cε2: (linker): 131.3<br>Cζ-OH: 156.3 |
| 3 | W7 | 8.82 | 4.96<br>53.4 | 3.31, 2.99<br>29.2 | Cγ: 110.7<br>δ1: 7.19 (s), 124.3<br>ε1: 10.80(NH)<br>Cε2: 136.4<br>Cδ2: 127.7<br>ε3: 7.80 (d), 119.6<br>ζ3: 6.98 (t), 118.7<br>η2: 7.04 (t), 121.4<br>ζ2: 7.30 (d), 111.7 |
| 4 | Y8 | 9.29 | 4.59<br>54.4 | 3.05, 2.91<br>36.0 | Cγ:<br>δ1: 6.88 (dd), 128.8<br>ε1: 6.44 (d), 121.3<br>δ2: 7.15 (s), 127.2<br>Cε2: linker: 131.4<br>Cζ-OH: 156.2 |
| 5 | D9 | 7.797<br>173.2 | 4.14<br>49.4 | 2.51<br>40.1 | Cγ=O: 173.7 |

Referenced to $^{1}$H (2.50 ppm) and $^{13}$C (40.3 ppm) of DMSO solvent peaks.

**Table S7. NMR assignments of ApyO-modified, GluC-digested ApyA-L7W Isomer–2**

| number | AA | NH/C=O | αH/C | βH/C | Others |
|---|---|---|---|---|---|
| 1 | G5 | <br>166.1 | 3.66, 3.59<br>40.5 | | |
| 2 | Y6 | 7.39<br>170.1 | 4.55<br>53.1 | 2.83, 2.72<br>36.5 | Cγ: 127.3<br>δ1: 6.54 (dd), 124.5<br>ε1: 6.81 (d), 116.2<br>δ2: 5.77 (d, 1.5Hz), 116.0<br>Cε2: 147.8 (linker)<br>Cζ-OH: 144.0 |
| 3 | W7 | 7.99<br>171.7 | 4.45<br>53.4 | 3.11, 2.99<br>29.0 | Cγ: 108.8<br>δ1: 7.05 (s), 124.7<br>ε1: 9.94 (NH)<br>Cε2: 136.1<br>Cδ2: 127.1<br>ε3: 7.4 (d), 118.4<br>ζ3: 6.99 (t), 119.4<br>η2: 7.10 (t), 121.9<br>ζ2: 7.35 (d), 111.8 |
| 4 | Y8 | 8.33<br>172.0 | 4.65<br>54.9 | 3.28, 2.64<br>37.9 | Cγ: 134.2<br>δ1: 7.36 (dd), 130.7<br>ε1: 6.99 (dd), 122.5<br>δ2: 7.15 (dd), 132.4<br>ε2: 6.80 (dd), 121.2<br>Cζ-O-: 153.3 (linker) |
| 5 | D9 | 8.30<br>175.2 | 4.57<br>50.3 | 2.86<br>36.5 | Cγ=O: 175.3 |

### Materials and Methods

#### Plasmid constructs

Wild type ApyA-ApyO, ApyA-ApyOIHIDS co-expression plasmids in the pRSF-Duet and pSCrha2 backbones were previously reported.[5] Mutations in *apyA* were conducted by PCR amplification of the backbone leading to construction of pRSF-ApyA-Y6W_ApyO, pRSF-ApyA-Y8W_ApyO, pRSF-ApyA-Y6W/Y8W_ApyO, pRSF-ApyA-L7Y_ApyO, pRSF-ApyA-L7W_ApyO, pRSF-ApyA-L7Y_ApyOHIDS and pSCrha2-ApyA-L7Y_ApyOHIDS (Table S1). Clones were maintained in *E. coli* DH5α and expressions were conducted in *E. coli* BL21 (DE3) TUNER cells or *Burkholderia* sp. FERM BP-2431 (for pSCrha2) unless otherwise mentioned.

#### Heterologous expression and purification of peptides

Bacterial growth and protein production in *E. coli* BL21(DE3) Tuner using the pRSF-constructs was conducted using a modified Terrific Broth medium according to a previous report[6] (yeast extract 24 g/L, tryptone 20 g/L, 17 mM $KH_2PO_4$, and 72 mM $K_2HPO_4$) supplemented with 2% glycerol, appropriate antibiotics and 1x final concentration of trace metal mix (Teknova). For cells co-expressing ApyHI, ApyD and ApyS in *E. coli*, the pIGB240 plasmid was introduced to assist in [Fe-S] cluster formation and vitamin B12 uptake as described previously[7] along with a pACYC-Duet plasmid ($Cam^R$) encoding the chaperonins Cpn10 and Cpn60 of *Oleispira antarctica*[8]. An overnight starter culture of the bacteria containing the appropriate plasmids was used to inoculate a 1 L subculture in 2 L flasks as 1% inoculum and grown at 37 ºC until the optical density at 600 nm ($OD_{600}$) reached ~1.2. The cells were then transferred to ice for 15 min followed by addition of 0.5 mM IPTG to induce gene expression, 1 mM freshly prepared ferrous citrate (0.1 mg $FeSO_4$ in 1 mL $Na_3C_6H_5O_7$) and 1 mM δ-aminolevulinic acid to support heme production (Catalog Number: 01433, CHEMIMPEX). For cells expressing ApyOHIDS, vitamin B12a (2 µM final) was also added during induction. Induced cells were cultivated at 16 ºC for the next 36 h at 150 rpm.

Protein production from *Burkholderia* sp. FERM BP-2431 (for pSCrha2) was conducted as previously reported in 2S4G medium composed of 20 g/L phytone peptone (Gibco), 40 g/L glycerol (Fisher Scientific), 2 g/L ammonium sulfate (Sigma), 2 g/L $CaCO_3$ (Sigma), and 405 µM $MgSO_4$.[5] Cells were grown in Erlenmeyer Baffled Cell Culture Flasks with vented caps containing a 0.22 µM filter at 30 ºC, shaking at 180 rpm until OD reached 0.8-1. L-Rhamnose (0.2% final) was added to induce gene expression along with iron(II) citrate (255 µM), L-cysteine (200 µM) and vitamin B12a (2 µM) followed by shaking at 30 ºC at 180 rpm for 24 h.

*E. coli* cells were then harvested by centrifugation at 5,000 × g at 4 ºC whereas *Burkholderia* cultures were harvested in two steps, a 1000 × g centrifugation step for 2 min to separate the $CaCO_3$ followed by re-centrifugation of the supernatant at 5,000 × g to harvest the cells. Cells were then re-suspended in 50 mM Tris-HCl buffer containing 300 mM NaCl, 10% glycerol, and 20 mM imidazole at pH 8.0 ($NPI_{20}$). For purification of peptides modified by ApyHI, 5 mM pyruvate was added to all buffers except the lysis buffer which contained 10 mM pyruvate. DNase I (Sigma), lysozyme (GoldBio), $MgCl_2$ (0.01 mg/mL, 0.1 mg/mL and 0.5 mM, respectively) and protease inhibitor cocktail (Pierce, A32955; 4 tablets per 100 mL of lysis buffer) were added.

The *E. coli* cell mixture was stirred for 1 h at 4 ºC followed by sonication at an amplitude of 50% for 20 cycles of 5 s on/off with a 12 mm probe. *Burkholderia* cells were resuspended in 50 mL of lysis buffer (6 M guanidinium hydrochloride, 50 mM Tris, 300 mM NaCl, pH 8) per 10 g of wet cells. The cell paste was lysed using a high-pressure homogenizer (Avestin, Inc) at an operating pressure between 15-20k psi. The homogenization was repeated 3-4 times.

The lysates were centrifuged at 22,000 × g at 4 ºC for 30 min. We observed a drop in pH to approx. pH 6 after cell lysis which was adjusted back to approx. pH 8 by addition of NaOH to facilitate Ni-NTA binding. After pH adjustment, Ni-NTA beads (MCLAB) were added to the supernatant and rotated 4 ºC for 1 h to facilitate batch binding. The pellet from the previous step was also resuspended in denaturation buffer (50 mM Tris-HCl buffer containing 300 mM NaCl, 10% glycerol, and 6 M guanidine hydrochloride at pH 8.0) followed by sonication and centrifugation as mentioned above. The denatured supernatant was collected and Ni-NTA beads were added. The mixtures were rotated at 4 ºC for 1 h before loading on to a column. At this point, Ni-NTA beads collected from both native and denaturing purification were loaded on the same column. The beads were washed 2 times with 3 × column volume (CV) of denaturing buffer, then 3 × CV of $NPI_{20}$ followed by 3 × CV of $NPI_{40}$. The desired protein was eluted twice with 2 × CV of $NPI_{750}$ (i.e., lysis buffer containing 750 mM imidazole). These conditions were used for all expression conditions unless stated otherwise.

**Isolation of matured C-terminal core peptide**

Following IMAC purification, the peptide eluent was desalted and the buffer exchanged to exchange buffer (50 mM Tris-HCl, 100 mM NaCl at pH 8.0) using a Sephadex PD10 column. In addition, 5 mM pyruvate was added to the exchange buffer for desalting peptides modified by ApyHI. To isolate the matured core peptide, proteolysis was conducted overnight using GluC endopeptidase (NEB) at 1:5000 enzyme:substrate ratio. The digested peptide was then desalted by solid phase extraction (SPE) using a Chromabond® c18ec column and subsequently purified by high-performance liquid chromatography (HPLC) as mentioned below.

**HPLC purification of modified peptide core fragments**

SPE column-eluted samples were lyophilized followed by resuspension in 10% acetonitrile containing 0.1% formic acid. The peptide sample was centrifuged at 12,000 xg for 10 min and the supernatant was subjected to HPLC purification. All purifications were conducted on a Vanquish UHPLC system (Thermo Fisher Scientific). For the purification of all compounds, a Phenomenex Aeris™ peptide C18-XB LC column (Part nr. 00G-4632-N0; particle size: 5 µm; dimensions 250 x 10 mm; pore size 100 Å) was used. Mobile phase used was solvent A containing water + 0.1% formic acid and solvent B containing acetonitrile + 0.1% formic acid. The flow rate for all purifications was 2 mL/min, and the column oven was maintained at 45 ºC.

For GluC-cleaved ApyA-Y6W fragment (modified GWLYD), separation was achieved over a 30-min method consisting of an initial equilibration segment (5 min at 5% solvent B) followed by a gradient of solvent B from 15% to 37.5% for the next 11 min, a wash step at 90% solvent B for 3 min and a re-equilibration step at 5% solvent B over the next 11 min. The desired product eluted between 14-14.5 min.

For GluC-cleaved ApyA-L7Y fragment (modified GYYYD), separation was achieved over a 30 min method consisting of an initial equilibration segment (5 min at 5% solvent B) followed by a gradient of solvent B from 15% to 38% for the next 11 min, a wash step at 90% solvent B for 4 min and a re-equilibration step at 5% solvent B over the next 10 min. The L7Y isomer-1 eluted at 12.6 min and the L7Y isomer-2 eluted at 12.85 min.

For GluC-cleaved ApyA-L7W fragment (modified GYWYD), separation was achieved over a 30 min method consisting of an initial equilibration segment (5 min at 5% solvent B) followed by a gradient of solvent B from 15% to 45% for the next 15 min, a wash step at 90% solvent B for 10 min and a re-equilibration step at 5% solvent B over the next 10 min. The L7W isomer-1 eluted at 13.1 min and the L7W isomer-2 eluted at 14.2 min.

**High-resolution tandem mass spectrometry**

The desalted protease-digested peptides were injected onto an Agilent 1290 LC-MS QToF for ESI-HR-MS and MS/MS analysis. LC separation was conducted at 50 ºC on a 5%-95% gradient of acetonitrile-water (+0.1% formic acid) over 8 min at 1 mL/min flow rate on a Phenomenex Aeris Widepore XB-C18 LC column (part nr. 00D-4482-E0). Mass spectra were collected in positive mode (for Leu7 mutants) or negative mode (for Arg1 mutants) at 10 spectra/s and 100 ms/spectrum. Tandem-MS fragmentation was achieved at normalized collision energies of 15, 20 and 25 eV. HR-MS/MS analysis was performed using the Interactive Peptide Spectral Annotator (IPSA) tool[1] and verified manually.

**Expression and purification of the cytochrome P450 ApyO**

*E. coli* BL21 (DE3) was transformed with a pRSF-6xHis-TEV-ApyO or pRSF-6xHis-MBP-ApyO ($Kan^R$) and a pACYC-Duet plasmid ($Cam^R$) encoding the chaperonins Cpn10 and Cpn60 of *Oleispira antarctica*[8] on a Luria Broth containing 1.5% agar plate supplemented with 50 µg/mL kanamycin and 34 µg/mL chloramphenicol. A single colony was picked and inoculated in Luria Broth containing the same amount of antibiotics overnight before being subcultured to 1 L of Terrific Broth [0.4 % (w/v) Glycerol, yeast extract 24 (g/L), tryptone 12 (g/L), 17 mM $KH_2PO_4$, and 72 mM $K_2HPO_4$] supplemented with 50 µg/mL kanamycin and 25 µg/mL chloramphenicol. The culture was grown at 37 ºC and shaken at 200-220 rpm. Then 1 mM IPTG, 2 mM $MgCl_2$, 160 mg δ-aminolevulinic acid, and 255 µM iron (II) citrate [concentrated stock freshly prepared by mixing 100 mg/mL iron (II) ammonium sulfate heptahydrate and 1 M sodium citrate] was added when the $OD_{600}$ of the culture reached 1.5 - 1.7. The culture was kept at 12 ºC and 200-220 rpm for 24 h. Afterwards, the cells were collected through centrifugation at 5000 x g for 15 min, resuspended in 50 mL of lysis buffer [50 mM HEPES-NaOH pH 7.5, 500 mM NaCl, 5% glycerol (v/v), 0.1% Triton X-100] per 10 g of estimated weight of wet cells. Here, the lysis buffer also contained 4 mg/mL lysozyme, 2 µM leupeptin, 2 µM benzamidine, and 2 µM E64. The tubes/cups containing the cells were placed in an ice-water bath and then lysed through sonication. The mixture was centrifuged at 30000 x g for 1 h and the supernatant (lysate) was collected and applied to a column containing Ni-NTA resin (2 mL of resin per L of cells, His-Pur, Thermo Scientific) pre-equilibrated with lysis buffer. After the Ni-NTA resin had been resuspended with the lysates, the lysate was allowed to flow through the column, and the Ni-NTA resin was washed with 10 resin volumes (RV) of lysis buffer (i.e., 20 mL lysis buffer for 2 mL resin). Afterwards, the Ni-NTA resin was washed with 15 RV of wash buffer [50 mM HEPES-NaOH pH 7.5, 1 M NaCl, 5% glycerol (v/v), 30 mM Imidazole]. His6-ApyO was eluted from the column using 5-6 RV of elution buffer (50 mM HEPES-NaOH pH 7.5, 300 mM NaCl, 5% glycerol (v/v), 250 mM Imidazole]. The buffer of the collected His6-ApyO solution was exchanged to 1000x volume of protein storage buffer (50 mM HEPES-NaOH pH 7.5, 300 mM NaCl, 2.5% glycerol) using a 30 kDa MWCO Amicon Ultra centrifugal filter (EMD Millipore) and concentrated until reaching a stock concentration of 50 µM. The tubes containing concentrated His6-ApyO were then flash frozen by liquid $N_2$ and stored at -80 ºC.

**Klenow fragment extension**

The coding DNA sequence for $ApyA_{ct}$ was synthesized in vitro using a DNA Polymerase I Klenow fragment-based polymerization strategy. Briefly, a universal 5’→3’ oligonucleotide fragment (FP; Table S3) was designed to contain a T7 promoter region followed by a ribosome binding sequence. Similarly, a 3’→5’ reverse primer (RP) sequence was designed to encode for the $ApyA_{ct}$ peptide sequence with an overlap on the forward primer at the 3’ region for annealing. The FP and RP (4 µM each, final concentration) were added to 1x NEB buffer 2 calculated for 50 µL reaction. Then 400 µM (final) of dNTP mix (NEB) was added to the mix followed by addition of 1 U of DNA polymerase I, Large (Klenow) fragment. The reaction volume was adjusted to 50 µL with nuclease free water (NFW) and incubated for 30 min at 25 ºC in a thermocycler. The reaction was stopped with addition of 10 mM (final) EDTA followed by heating the mix for 20 min at 75 ºC. The DNA product was then precipitated by addition of 2.5 volumes of ice-cold absolute ethanol and 0.3 M sodium acetate. Following centrifugation for 30 min at 13000 x g (4 ºC), the supernatant was discarded and the pellet was washed twice with ice-cold 75% ethanol. The resulting pellet was air-dried and resuspended in 20 µL of NFW.

***In vitro* assays**

The purified DNA fragment from the section above was used as a template in a PURExpress cell free translation (NEB) set up to obtain the desired substrate. CFE reactions were set up as per the manufacturer’s instructions in 10 µL scale and with addition of 500 ng of the DNA template. The CFE reactions were incubated at 37 ºC for 4 h to produce the peptide substrate.

After the CFE reaction was completed, the reaction mixture was split into two 5 µL aliquots. To both aliquots NADPH (3 µM, final concentration), *E. coli* ferredoxin reductase (10 µM, final), *E. coli* ferredoxin (25 µM, final), and HEPES buffer at pH 7.5 (50 mM, final) were added. To one of the aliquots ApyO purified from *E.*

*coli* (10 µM, final) was added while the second aliquot contained no ApyO (replaced by NFW) as a control. Total reaction volume was maintained at 15 µL for all reactions. The reaction was incubated at 25 ºC for 16 h followed by quenching with 50% methanol containing 0.1% formic acid (final). The mixture was then centrifuged, and the supernatant was dried under vacuum using a speedvac. The dried pellets were subsequently resuspended in 10 µL of 10% aqueous acetonitrile containing 0.1% formic acid and subjected to LCMS analysis.

**Kinetic experiments**

For kinetic experiments, a synthetic peptide was purchased from Lifetein (*N*-acetylated GRMGEGYLYD) and dissolved in water to make a 1.5 mM solution (confirmed by qNMR analysis). Substrate to product conversion was determined by LC-HRMS analysis by integrating the area under the curve (AUC) for the extracted ion chromatograms. To interpolate the AUC to concentration, we established a standard curve for the substrate with concentrations ranging from 0.2 µM to 12.8 µM. Purified MBP-ApyO was used for kinetic assays because of its higher solubility and yield. A substrate range of 10, 20, 40, 80, 160, 250, 320, 480 and 640 µM was used with 1 µM of enzyme. The redox partners ferredoxin (NCBI accession ID: WP_001124469.1), ferredoxin reductase (NCBI accession ID: WP_000796310.1) and NADPH were added at 25 µM, 10 µM and 1.5 mM final concentrations, respectively in 50 mM Hepes pH 7.5. Assays were conducted at 37 ºC on a 100 µL scale. At 2 min time intervals, 10 µL of the reaction was quenched with 10 µL of ice-cold acetonitrile (MeCN) containing 1% formic acid (final: 50% MeCN:0.5% FA). Quenched samples were centrifuged at 10000 x g for 5 min to remove the precipitated enzyme and the supernatant was analyzed by LC-HRMS.

**NMR data acquisition and analysis**

Sample information: five peptides were studied by NMR spectroscopy in this work. The peptides ApyA-Y6W (~ 1.8 mM) in DMSO-d6 at 55 ºC, ApyA-L7Y-isomer-1 (~ 1.0 mM) in 90% $H_2O$, 10% $D_2O$ and 0.2% formic acid-$d_2$ (dFA) at 45 ºC, ApyA-L7Y-isomer-2 (~ 2.7 mM) in 90% $H_2O$, 10% $D_2O$ and 0.2% dFA at 25 ºC, ApyA-L7W-isomer-1 in DMSO-d6 and 0.2% dFA (~ 0.3 mM) at 25 ºC, and ApyA-L7W-isomer-2 (~ 0.8 mM) in 90% $H_2O$ and 10% $D_2O$ and 0.2% dFA at 25 ºC were dissolved in ~ 550 µL volume in a 5-mm Wilmad 535-pp NMR tube. NMR data were collected at the temperatures listed on a Bruker Avance NEO 600 MHz spectrometer equipped with a 5-mm BBO prodigy probe or a Bruker Avance III 500 MHz spectrometer equipped with a 5-mm BBFO cryoprobe, or an Agilent VNMRS 750 MHz NMR spectrometer equipped with a room temperature indirect detection 5-mm HCN probe. One-dimensional (1D) $^1$H NMR, two-dimensional (2D) homonuclear $^1$H-$^1$H COSY (correlation spectroscopy which reveals the correlation between two neighboring protons), $^1$H-$^1$H TOCSY (total correlation spectroscopy which reveals the correlation of protons in the same spin system) at 80 ms mixing time, $^1$H-$^1$H NOESY (Nuclear Overhauser Effect spectroscopy which reveals close proximity in space between two protons) at 400 ms mixing time, $^1$H-$^{13}$C HSQC (heteronuclear single quantum coherence spectroscopy, revealing one-bond correlation between $^1$H and $^{13}$C), and $^1$H-$^{13}$C HMBC (heteronuclear multiple bond correlation, revealing long range $^1$H-$^{13}$C connectivity such as 2-bond, 3-bond, or 4 or more bonds) were collected.

Spectra were acquired for each sample using either pulse programs in Bruker Topspin 4.1.4, Topspin 3.7.0, or the Biopack pulse sequences in the VNMRJ 4.2A software. The spectra were processed and analyzed in Mnova (version 15.1.0.; Mestrelab Research). The sample concentrations were estimated by $^{1}H$ spectra collected under the qNMR condition on either the Bruker Avance NEO 600 MHz spectrometer compared with a standard sample (ERETIC 2 method) or an Agilent VNMRS 750 MHz spectrometer with a HCN probe that was calibrated with a known standard. On both instruments, 48.5 mM triphenylphosphate in $CDCl_3$ was used as the external calibration compound.

**Protease inhibition assays**

The concentration of all peptides was determined by NMR (see above). Test compounds were dissolved in deionized water. Protease inhibition assays were conducted using fluorescence detection for all enzymes as shown in the tables below. Test compounds were incubated with the enzyme for 15 min at room temperature before addition of the corresponding substrates (see Tables below). Assays were conducted in 96 well black opaque half area plates (3694, Corning) and kinetic readings were taken every 60 secs for 1 h. Measurements were conducted on an Agilent Synergy H1 plate reader using the extended gain functionality and auto read height adjustment. Slope function was used to calculate relative inhibition in the linear range of enzyme activity as previously described.[6] $IC_{50}$ calculations were conducted using GraphPad Prism v10.6.1 using a non-linear regression (curve fit) of the [inhibitor] *vs.* response variable slope functionality at 95% confidence interval (CI) of parameters and asymmetrical (profile-likelihood) CI option. Inhibitor controls used were leupeptin for cathepsin B, cathepsin L and papain or captopril for ACE at 15 µM concentration. SARS-CoV-2 Main Protease Inhibitor Screening Assay Kit (Item No. 701960; Cayman chemical) was used for $M^{pro}$ inhibition assays as per the manufacturer's protocol.

| Cathepsin L (16-12-030112; Athens research and technology) | | | |
|---|---|---|---|
| Substrate: Z-FR-AFC (866-75; Echelon Biosciences) | | | |
| Assay buffer:<br>50 mM sodium acetate, pH 5.5, 2.5 mM EDTA, 2.5 mM DTT | | | |
| Reaction temperature: 30 ºC | | | |
| **Component** | **Stock concentration** | **Volume added** | **Final concentration** |
| Enzyme | 1 nM | 10 µL | 200 pM |
| Substrate | 50 µM | 10 µL | 10 µM |
| Compound | 5× desired | 10 µL | variable |
| Assay buffer | – | 20 µL | – |
| **Total** | – | **50 µL** | – |

| Cathepsin B (16-12-030102; Athens research and technology) | | | |
|---|---|---|---|
| Substrate: Z-RR-4MBNA (866-31, Echelon Biosciences) | | | |
| Assay buffer:<br>100 mM sodium phosphate, pH 6.0, 1.33 mM EDTA, 2 mM DTT | | | |
| Reaction temperature: 30 ºC | | | |
| **Component** | **Stock concentration** | **Volume added** | **Final concentration** |
| Enzyme | 0.962 nM | 10 µL | 192.4 pM |
| Substrate | 100 µM | 10 µL | 20 µM |
| Compound | 5× desired final | 10 µL | variable |
| Assay buffer | – | 20 µL | – |
| **Total** | – | **50 µL** | – |

| Papain (P4762-25MG; Sigma) | | | |
|---|---|---|---|
| Substrate: Z-FR-AFC (866-75; Echelon Biosciences) | | | |
| Assay buffer:<br>50 mM sodium phosphate, pH 7, 1 mM EDTA, 2 mM DTT<br>Preactivated for 10 min at room temperature in assay buffer | | | |
| Reaction temperature: 30 ºC | | | |
| **Component** | **Stock concentration** | **Volume added** | **Final concentration** |
| Enzyme | 1 nM | 10 µL | 200 pM |
| Substrate | 50 µM | 10 µL | 10 µM |
| Compound | 5× desired | 10 µL | variable |
| Assay buffer | – | 20 µL | – |
| **Total** | – | **50 µL** | – |

| Angiotensin I-converting enzyme (ACE) (A6778; Sigma) | | | |
|---|---|---|---|
| Substrate: Abz-FRK(Dnp)-P-OH (HY-P1853; MedChemExpress) | | | |
| Assay buffer:<br>50 mM HEPES, pH 7.5, 300 mM NaCl, 10 µM $ZnCl_2$ | | | |
| Reaction temperature: 37 ºC | | | |
| **Component** | **Stock concentration** | **Volume added** | **Final concentration** |
| Enzyme | 2 mU/mL | 10 µL | 0.4 mU/mL |
| Substrate | 50 µM | 10 µL | 10 µM |
| Compound | 5× desired final | 10 µL | variable |
| Assay buffer | – | 20 µL | – |
| **Total** | – | **50 µL** | – |

| Trypsin (V5111; Promega) | | | |
|---|---|---|---|
| Substrate: Boc-QAR-AMC (HY-134432B; MedChemExpress) | | | |
| Assay buffer:<br>50 mM Tris HCl, pH 8, 100 mM NaCl, 2 mM $CaCl_2$<br>Preactivated for 15 min at 30 ºC | | | |
| Reaction temperature: 30 ºC | | | |
| **Component** | **Stock concentration** | **Volume added** | **Final concentration** |
| Enzyme | 2.5 nM | 10 µL | 500 pM |
| Substrate | 50 µM | 10 µL | 10 µM |
| Compound | 5× desired | 10 µL | variable |
| Assay buffer | – | 20 µL | – |
| **Total** | – | **50 µL** | – |

| Chymotrypsin (V1061; Promega) | | | |
|---|---|---|---|
| Substrate: Suc-AAPF-AMC (881-31; Echelon Biosciences) | | | |
| Assay buffer:<br>50 mM Tris HCl, pH 8, 100 mM NaCl, 2 mM $CaCl_2$ | | | |
| Reaction temperature: 30 ºC | | | |
| **Component** | **Stock concentration** | **Volume added** | **Final concentration** |
| Enzyme | 5 nM | 10 µL | 1 nM |
| Substrate | 50 µM | 10 µL | 10 µM |
| Compound | 5× desired final | 10 µL | variable |
| Assay buffer | – | 20 µL | – |
| **Total** | – | **50 µL** | – |
